# Restoration of PITPNA in Type 2 diabetic human islets reverses pancreatic beta-cell dysfunction

**DOI:** 10.1101/2022.07.06.498991

**Authors:** Yu-Te Yeh, Chandan Sona, Xin Yan, Adrija Pathak, Mark I. McDermott, Zhigang Xie, Liangwen Liu, Anoop Arunagiri, Yuting Wang, Amaury Cazenave-Gassiot, Adhideb Ghosh, Ferdinand von Meyenn, Sivarajan Kumarasamy, Sonia M. Najjar, Shiqi Jia, Markus R. Wenk, Alexis Traynor-Kaplan, Peter Arvan, Sebastian Barg, Vytas A. Bankaitis, Matthew N. Poy

**Author notes:** Equal contribution (Yu-Te Yeh, Chandan Sona). **Correspondence and reprint requests should be addressed to:** Matthew N. Poy, Ph.D., Johns Hopkins All Children’s Hospital, Johns Hopkins University School of Medicine, Department of Medicine, Division of Endocrinology, Diabetes and Metabolism, Institute for Fundamental Biomedical Research, 600 Sixth Avenue South, St. Petersburg, Florida 33701.

## Abstract

Defects in insulin processing and granule maturation are linked to pancreatic beta-cell failure during type 2 diabetes (T2D). Phosphatidylinositol transfer protein alpha (PITPNA) stimulates activity of phosphatidylinositol (PtdIns) 4-OH kinase to produce sufficient PtdIns-4-phosphate (PtdIns-4-P) in the trans-Golgi network to promote insulin granule maturation. *PITPNA* in beta-cells of T2D human subjects is markedly reduced suggesting its depletion accompanies beta-cell dysfunction. Conditional deletion of *Pitpna* in the beta-cells of *Ins*-Cre;*Pitpna^flox/flox^* mice leads to hyperglycemia resulting from decreased glucose-stimulated insulin secretion (GSIS) and reduced pancreatic beta-cell mass. Furthermore, *PITPNA* silencing in human islets confirmed its role in PtdIns-4-P synthesis and led to impaired insulin granule maturation and docking, GSIS, and proinsulin processing with evidence of ER stress. Restoration of *PITPNA* in islets of T2D human subjects reversed these beta-cell defects and identify *PITPNA* as a critical target linked to beta-cell failure in T2D.

## INTRODUCTION

Type 2 diabetes (T2D) is a non-autoimmune disease of impaired insulin signaling that afflicts ~10% of the population in the United State alone (Eizirik et al., 2020; Nolan and Prentki, 2019). Both impaired insulin release and reduced beta-cell mass contribute to the beta-cell failure that occurs during T2D (Campbell and Newgard, 2021; Porksen et al., 2002; Rhodes, 2005). Beta-cell failure is calculated to associate with a 24-65% loss of measurable beta-cell mass and a 50-97% loss of secretory capacity after disease onset (Chen et al., 2017; Liu et al., 2021). Prior to the events that cause this decline, beta-cells functionally accommodate peripheral insulin resistance for a limited time in two ways. First, beta-cells increase insulin production (Campbell and Newgard, 2021; Nolan and Prentki, 2019). Second, beta-cells increase their proliferation to expand the pool of insulin-producing cells in order to compensate for increased metabolic demand (Alejandro et al., 2015; Ferrannini, 2010; Muoio and Newgard, 2008). However, beta-cells in T2D patients ultimately succumb to multiple complications that include endoplasmic reticulum (ER) stress, glucotoxicity, and dedifferentiation (Back and Kaufman, 2012; Fonseca et al., 2011; Shrestha et al., 2021; Talchai et al., 2009). The precise mechanisms underlying the decline of both beta-cell secretion and mass remain unclear. As a result, worldwide efforts continue to focus on identifying the molecular bases of these defects (Rohm et al., 2022; van Raalte and Verchere, 2017; Yong et al., 2021).

Phosphoinositides define a set of chemically distinct phosphorylated derivatives of the glycerophospholipid phosphatidylinositol (Balla, 2013). The central importance of phosphoinositide signaling in regulating cellular homeostasis in eukaryotes is demonstrated in two ways. First, the diversity of cellular activities regulated by phosphoinositide metabolism is striking. Phosphoinositide signaling controls cellular functions that range from membrane trafficking to receptor signaling at the plasma membrane, autophagy, transcription, mRNA transport, cytoskeleton dynamics, and numerous other activities (De Camilli et al., 1996; Di Paolo and De Camilli, 2006; Hokin and Hokin, 1953; Martin, 1998). Second, even subtle derangements in phosphoinositide metabolism contribute instrumentally to many diseases -- including diabetes (Bridges and Saltiel, 2012; Rameh and Deeney, 2016; Wuttke, 2015). Phosphatidylinositol transfer proteins (PITPs) are highly conserved molecules that regulate the interface between lipid metabolism and cellular functions (Bankaitis et al., 2010; Cleves et al., 1991). PITPs promote the activity of phosphatidylinositol (PtdIns) 4-OH kinases and PtdIns-4 phosphate (PtdIns-4-P) synthesis in eukaryotic cells (Ashlin et al., 2021; Lete et al., 2020; Xie et al., 2018). There are at least three soluble PITPs expressed in mammals -- PITPα/PITPNA, PITPβ/PITPNB, and rdgBβ/PITPnc1 (Dickeson et al., 1989; Fullwood et al., 1999; Tanaka and Hosaka, 1994). PITPNA and PITPNB share ~77% sequence identity, are encoded by distinct genes, and both PITPNA and PITPNB are characterized as transfer proteins for several phospholipids including PtdIns in vitro (Helmkamp et al., 1974; Wirtz, 1991). However, rather than functioning as inter-organelle lipid transfer proteins in cells, all available data are more consistent with PITPs serving as metabolic sensors that facilitate the presentation of PtdIns to PtdIns 4-OH kinases in vivo -- thereby channeling PtdIns-4-P signaling to specific (and diverse) biological outcomes (Schaaf et al., 2008). In that regard, PITPs contribute to secretory vesicle formation from the trans-Golgi network (TGN), to Ca^2+^-activated secretion in permeabilized neuroendocrine cells, and to the regulation of Golgi dynamics in embryonic neural stem cells of the developing mouse neocortex (Bankaitis et al., 1990; Hay and Martin, 1993; Lete *et al*., 2020; Ohashi et al., 1995; Xie and Bankaitis, 2022; Xie *et al*., 2018).

Here we first demonstrate that functional ablation of *Pitpna* in murine beta-cells results in random-fed hyperglycemia due to both impaired glucose-stimulated insulin secretion (GSIS) and reduced beta-cell number. These defects are accompanied by induction of ER stress and deranged mitochondrial dynamics and performance. Consistent with the murine studies, we further show that expression of *PITPNA* (referred to as human *PITPNA* and mouse *Pitpna*) is markedly diminished in pancreatic islets of T2D human subjects compared to non-diabetic donors. Such a downregulation is of functional consequence as reduced *PITPNA* levels in isolated human islets compromised PtdIns-4-P synthesis in the Golgi system, impaired insulin granule maturation and docking, and induced both ER and mitochondrial stress. Finally, we demonstrate that restoration of PITPNA expression in isolated pancreatic islets from T2D human subjects rescued insulin secretory capacity and granule biogenesis and alleviated ER stress. Taken together, these results establish that diminished PITPNA function is a major cell-autonomous contributor to reduced beta-cell mass and insulin output and, ultimately, to the beta-cell failure that represents a cardinal feature of T2D pathogenesis.

## RESULTS

### *Pitpna* is a direct target of miR-375 in the pancreatic beta-cell

The microRNA miR-375 is a potent regulator of insulin secretion that directly targets expression of several genes including *Myotrophin, Cadm1, Gephyrin (Gphn)*, and *Elavl4/HuD* (Poy et al., 2004; Poy et al., 2009; Tattikota et al., 2014; Tattikota et al., 2013). An extended analysis using the TargetScan algorithm identified a candidate binding site for miR-375 in the 3’UTR of the gene *Pitpna* (Agarwal et al., 2018). This gene encodes a phosphatidylinositol transfer protein and is expressed in pancreatic beta-cells (Figure 1A). To determine whether *Pitpna* is a genuine miR-375 target, the full-length mouse *Pitpna* 3’UTR (2709-nt, *Pitpna* WT) was subcloned into a luciferase reporter construct. The effects of modulating miR-375 activity on expression of this reporter were then determined. As expected, luciferase expression was inhibited in the presence of the miR-375 mimic (375-mimic) relative to its expression when cells were incubated with a pool of scrambled control mimics (Ctrl-mimic) (Figure S1A). Moreover, miR-375 directly targets this specific site binding site as evidenced by our observation that site-directed mutagenesis of the candidate binding site in the 3’UTR (*Pitpna* MUT) abolished the inhibitory effect of the miR-375 mimic (Figure S1A). To test whether endogenous *Pitpna* expression is subject to regulation by miR-375 in vivo, murine insulinoma MIN6 cells were transfected with an inhibitory antisense RNA oligonucleotide directed against miR-375 (Antg-375) to reduce expression of this miRNA (Figure S1B). The Antg-375-mediated silencing of miR-375 resulted in increased *Pitpna* mRNA levels when compared to cells transfected with a control pool of scrambled antisense oligonucleotides (Antg-Ctrl) (Figure S1B). Immunoblot analyses confirmed that inhibition of miR-375 resulted in elevated steady-state levels of Pitpna as well as other miR-375 targets (i.e. Cadm1, Gphn). Conversely, transfection with the 375-mimic reduced the steady-state levels of all three of these proteins in a dose-dependent manner (Figures S1C, D). Direct binding of miR-375 with its target genes is mediated by the RNA-binding protein Argonaute2 (Ago2) (Tattikota *et al*., 2014). Consistent with the abolition of miR-375 action, conditional deletion of *Ago2* in pancreatic beta-cells (*Ins*-Cre, *Ago2^flox/flox^*) de-repressed *Pitpna, Cadm1* and *Gphn* expression (Figure S1E).

**Figure 1.**
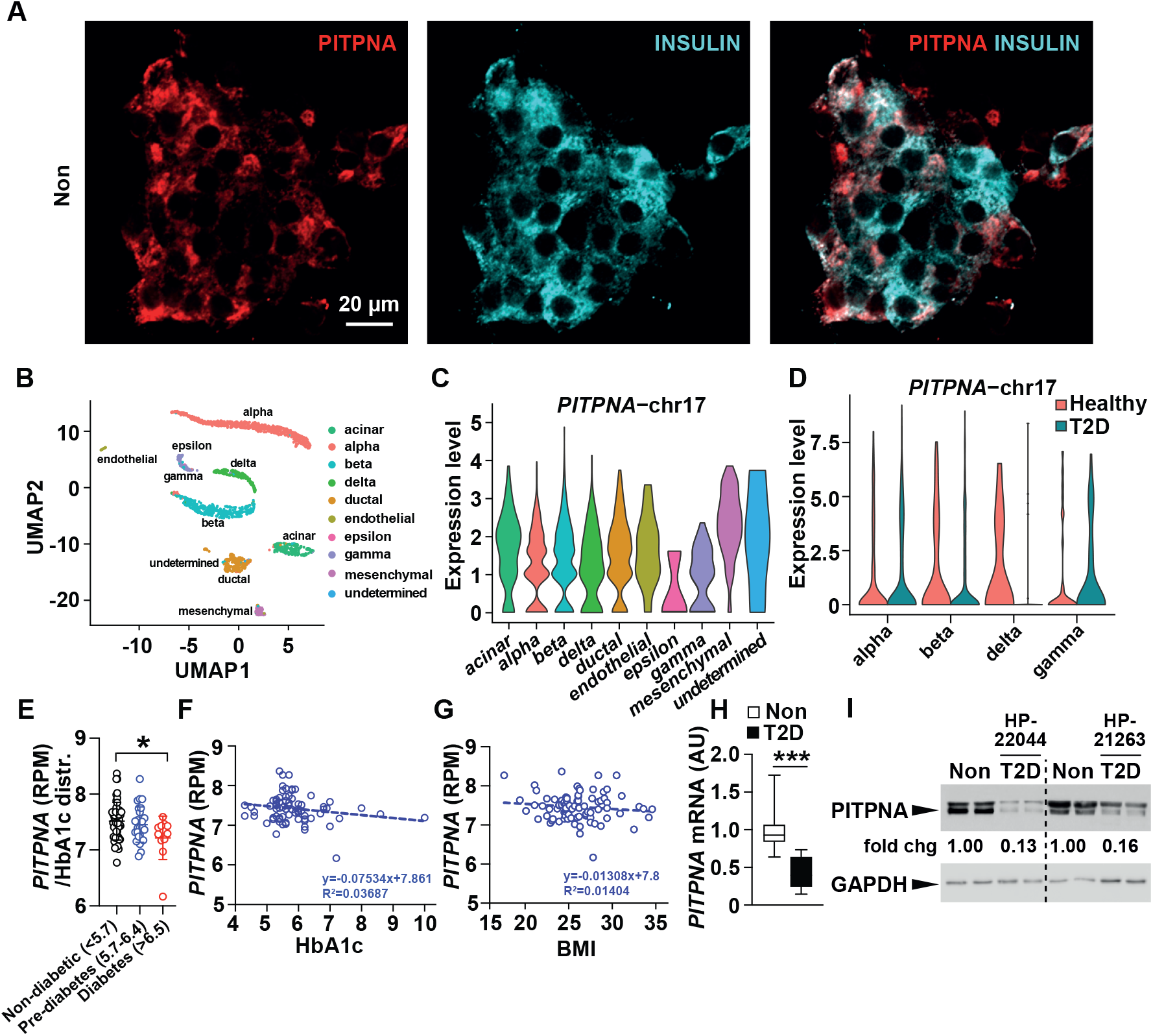
*PITPNA* expression is decreased in isolated islets of T2D human subjects. **A**, Immunostaining of endogenous INSULIN (cyan), and PITPNA (red) expression within pancreatic islets isolated from a non-diabetic human subject. Scale bar = 20μm. **B**, UMAP projection and graph-based clustering of scRNA-Seq analysis performed on isolated human pancreatic islet cell types. **C**, Relative abundance of *PITPNA* in islet cell clusters from human donors. **D**, Comparison of *PITPNA* expression in islet endocrine cell types from T2D (green) and non-diabetic donors (red). **E**, Normalized *PITPNA* expression from bulk RNA sequencing of isolated human islets across non-diabetic (HbA1c levels<5.7, n=51), pre-diabetic (HbA1c between 5.7 and 6.4, n=27), and T2D (HbA1c >6.5, n=11) human subjects. Normalized expression values are shown in reads per million (RPM). **F**, Correlation analysis between normalized islet *PITPNA* expression and HbA1c of human subjects (n=77). The R^2^ value indicates the correlation coefficient. **G**, Correlation analysis between normalized islet *PITPNA* expression and body mass index (BMI) of human subjects (n=89). The R^2^ value indicates correlation coefficient. **H**, qRT-PCR analysis of *PITPNA* mRNA expression in pancreatic islets isolated from non-diabetic (n=15) and T2D (n=5) human donors. **I**, Western blot analysis of PITPNA expression in isolated islets of non-diabetic human donors (Non) and T2D donors (T2D). Results presented as mean ± SEM. \**P*< 0.05.

Pitpna activity represents an interesting target for miR-375 control as it is an established mediator of PtdIns-4-P synthesis within the mammalian TGN (Lete *et al*., 2020; Xie *et al*., 2018), and PtdIns-4-P is required for the recruitment of budding factors and secretory granule formation (Cruz-Garcia et al., 2013). Further evidence in support of *Pitpna* expression being a physiologically relevant miR-375 target in beta-cells was provided by quantitative liquid chromatography-tandem mass spectrometry (LC/MSMS) analyses of the MIN6 murine insulinoma cell line lipidome as a function of Pitpna expression. Transfection of MIN6 cells with Antg-375 oligonucleotides to inhibit miR-375 resulted in the elevation of total bulk PtdIns in these cells as well as increased levels of multiple PtdIns molecular species (Figure S1F). Given the elevated insulin secretory output observed after inhibition of miR-375 (Poy *et al*., 2004), these results suggest that *Pitpna* functional status is linked to PtdIns metabolism in the murine beta-cell.

### Decreased PITPNA expression in isolated islets of T2D human subjects

We next examined whether PITPNA functional status is an important factor in human diabetes. Datasets obtained from transcriptomic RNA sequencing analyses performed on isolated human islet cells were interrogated for altered *PITPNA* expression (Fadista et al., 2014; Muraro et al., 2016; Xin et al., 2016). Notably, single cell RNA-seq analyses showed *PITPNA* expression was reduced in beta-cells from T2D donors in comparison to non-diabetic human donors with no change observed in alpha, and gamma-cell populations (Figures 1B-D) (Xin *et al*., 2016). Moreover, analyses of global transcriptomic RNA sequencing data from islets of human subjects stratified according to hemoglobin A1C (HbA1c) levels were also informative. HbA1c is a measure of long-term glycemia and the patients were classified as non-diabetic (HbA1c<5.7%), pre-diabetic (5.7-6.4), and diabetic (>6.5) (Fadista *et al*., 2014). *PITPNA* gene expression (reads per million) was reduced in islets of T2D subjects (HbA1c >6.5) relative to the expression levels recorded for islets of non-diabetic controls (HbA1c <5.7, Figure 1E). Indeed, *PITPNA* expression was inversely correlated with both HbA1c levels and body mass index (BMI) across all subjects (Figures 1F, G). This inverse correlation indicates that both body weight and glycemic status are parameters associated with changes in *PITPNA* expression. Expression analyses using quantitative real-time polymerase chain reaction (qRT-PCR) and immunoblotting further corroborated reduced PITPNA expression in isolated islets of T2D donors compared to non-diabetic control subjects (Figures 1H, I and Table S1). These collective data demonstrate that reduced *PITPNA* expression in pancreatic beta-cells of human subjects is associated with several hallmarks of predisposition to T2D.

### Whole body *Pitpna* knockout mice exhibit decreased pancreatic beta-cell mass

One of the signature phenotypes of *Pitpna* whole body knockout mice is reduced pancreatic islet numbers marked by shrunken islet morphologies and vacuolations (Alb et al., 2003). As the majority of *Pitpna* total-body knockout (*Pitpna* KO) mice die within the first 48 hours after birth, pancreata were isolated from these animals within the first 24 hours of birth and subjected to islet morphometric analysis. In addition to quantifying the reduction in overall islet number, we observed that the number of insulin^+^ cells per area of pancreas (mm^2^) appeared reduced in *Pitpna* KO mice compared to littermate controls (Figures S2A, B). These reductions in beta-cell number were accompanied by a proportional decline in total pancreatic insulin content (Figure S2C). By contrast, proinsulin levels were elevated in whole pancreas lysates derived from *Pitpna* null mice relative to controls and these data indicate that loss of *Pitpna* expression compromised the relative efficiency of proinsulin processing for insulin storage (Figure S2C). Analysis of *Pitpna* null beta-cells by transmission electron microscopy (TEM) showed reduced numbers of docked granules at the plasma membrane, a reduction in the number of mature granules, and reduced overall granule size in the mutant islets relative to littermate controls (Figures S2D-F). Lastly, terminal nucleotidyl transferase dUTP nick end labeling (TUNEL) experiments revealed a significant increase in the number of beta-cells undergoing apoptosis in *Pitpna* whole body knockout pancreas compared to controls (Figures S2G, H). The apoptotic phenotype was cell-specific. No changes were observed in TUNEL staining of the glucagon^+^ cell population (Figure S2I), or in Ki-67^+^ beta-cell numbers in *Pitpna* null mice (Figures S2J, K).

### Conditional deletion of *Pitpna* in beta-cells impairs glucose-stimulated insulin secretion (GSIS)

To more specifically assess the role of Pitpna in beta-cell physiology, two approaches were taken. First, we transfected the murine insulinoma cell line MIN6 with either a scrambled siRNA control pool (si-*Ctrl*) or an siRNA pool targeting *Pitpna* (si-*Pitpna*) designed to achieve a reduction in Pitpna expression at least as great as that seen in the beta-cells of T2D islets, in order to look directly at the role of *Pitpna* in pancreatic beta-cell function. The transfected cells were subsequently treated with glucose in concentrations ranging from 5.5mM to 25mM and insulin secretion responses were measured. Indeed, insulin release in response to 10 and 25mM glucose was markedly reduced in MIN6 cells inhibited for *Pitpna* expression (Figure S3A-C). Likewise, intracellular insulin content was also significantly reduced in MIN6 cells incubated in 25mM glucose -- indicating Pitpna contributes to insulin expression, its processing and/or insulin granule biogenesis (Figure S3D). Conversely, transfection of MIN6 cells with an expression construct encoding the *Pitpna* cDNA increased cellular *Pitpna* expression and elevated both GSIS and insulin content in cells challenged with 25mM glucose (Figure S3E-H).

Second, Pitpna function was specifically assessed in beta-cells of adult mice. To that end, conditional beta-cell-specific *Pitpna* null mice were generated by crossing *Pitpna*-floxed animals with mice expressing Cre recombinase under control of the mouse *Insulin1* promoter (*Ins*-Cre, *Pitpna^flox/flox^* mice) (Thorens et al., 2015). Immunoblot analyses confirmed a significant reduction in Pitpna expression in isolated islets of *Ins*-Cre, *Pitpna^flox/flox^* mice by age 10 weeks (Figure 2A). While blood glucose levels were unchanged after an 8 hour fast, *Ins*-Cre, *Pitpna^flox/flox^* mice exhibited elevated random-fed blood glucose in addition to reduced plasma insulin levels relative to control animals (Figure 2B). Moreover, *Ins*-Cre, *Pitpna^flox/flox^* mice exhibited reduced plasma insulin and elevated blood glucose levels in response to an intraperitoneal glucose bolus (Figures 2C, D). Taken together these data diagnose impaired insulin secretion in *Pitpna*-deficient mice relative to Pitpna-sufficient control littermate mice. These results were further corroborated by a significant blunting of the ex vivo secretory response to both 16.7mM glucose and 40mM KCl in *Pitpna*-null islets relative to control islets (Figure 2E). High resolution analyses of granule morphology by TEM reported increased numbers of immature and empty insulin secretory granules and reductions in the numbers of both mature secretory granules and docked granules in *Ins*-Cre, *Pitpna^flox/flox^* islets (Figures 2F-H). Moreover, genetic ablation of *Pitpna* in beta-cells showed significant reductions in islet and beta cell numbers and beta-cell mass (Figure 2I, J). These declines were in part attributed to apoptotic cell death as TUNEL staining of pancreata of 8-week-old *Ins*-Cre, *Pitpna^flox/flox^* mice showed an increase in the number of TUNEL^+^ beta-cells relative to littermate control mice (Figures 3A, B). No significant increases in TUNEL staining were detected in Gcg+ alpha cells of *Ins*-Cre, *Pitpna^flox/flox^* mice (Figure 3C). These phenotypes in the beta-cell-specific *Pitpna* gene eviction model were similar to the results obtained for *Pitpna* whole body knockout mice. These results demonstrate that Pitpna: (i) is a potent regulator of beta-cell viability, (ii) is required for insulin granule maturation and secretion in beta-cells, and (iii) beta-cell *Pitpna* deficiency is sufficient to disrupt systemic glucose homeostasis in an animal model.

**Figure 2.**
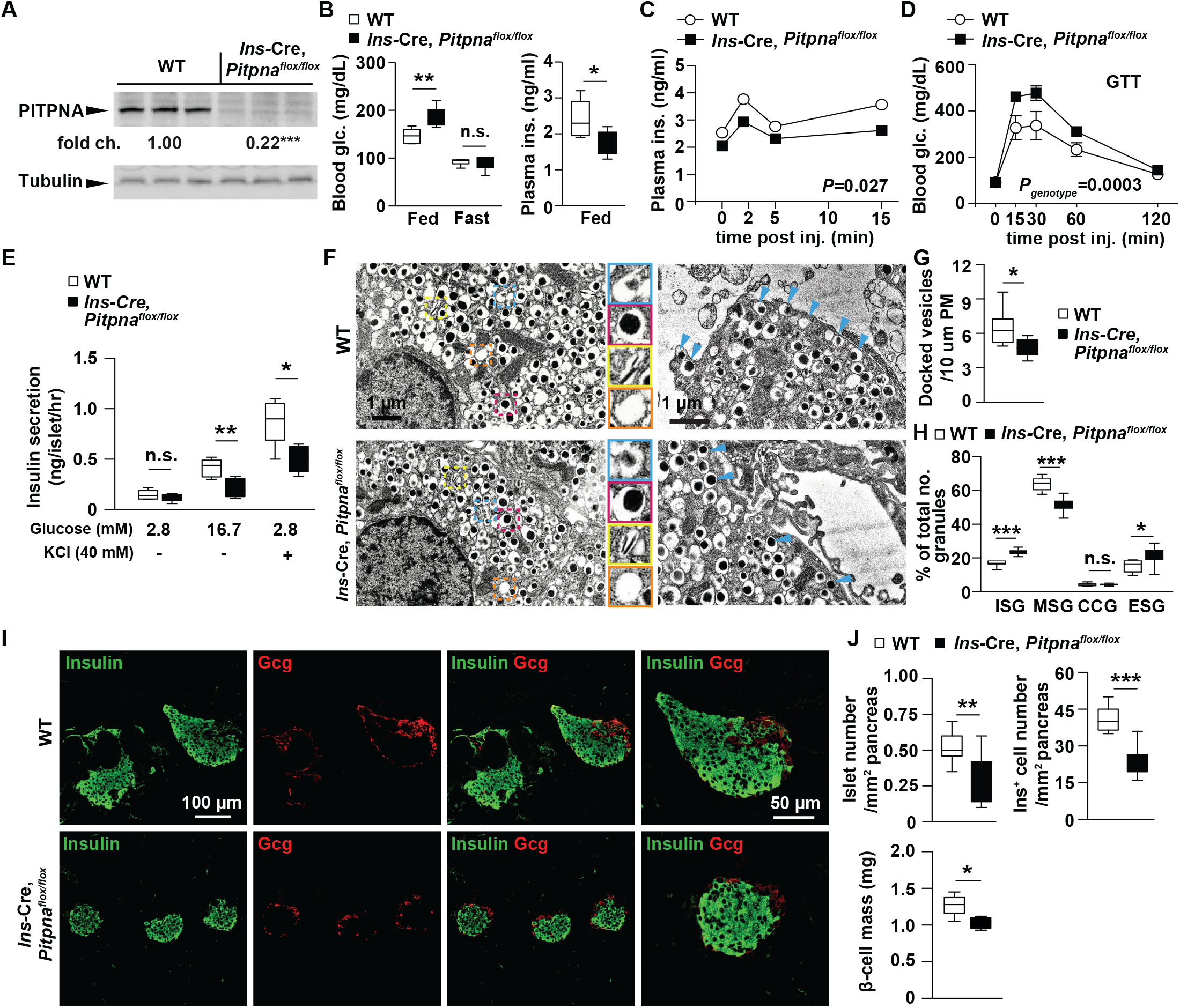
Conditional deletion of *Pitpna* in the pancreatic beta-cell impairs glucose-stimulated insulin secretion. **A**, Western blot analysis of Pitpna in isolated islets from *Ins*-Cre, *Pitpna^flox/flox^* and littermate control wildtype (WT) mice at age 8 weeks (n=3). Metabolic parameters were assessed in *Ins*-Cre, *Pitpna^flox/flox^* and WT mice at age 8 weeks including: **B**, random-fed and overnight 16-hour fasted blood glucose and plasma insulin (n=6), **C**, plasma insulin after glucose bolus (n=6), and **D**, blood glucose measurements after glucose bolus (n=6). **E**, Quantification of insulin release in response to 2.8mM and 16.7mM glucose concentrations and KCl (40mM) from isolated islets of 10-week-old *Ins*-Cre, *Pitpna^flox/flox^* and WT mice (n=5). **F**, Representative transmission electron micrographs of pancreatic beta-cells from 10-week-old *Ins*-Cre, *Pitpna^flox/flox^* and WT mice. Quantification of **G**, docked vesicles, and **H**, granule morphology (immature secretory granule (ISG, blue box), mature secretory granules (MSG, red box), crystal-containing granules (CCG, yellow box), and empty secretory granules (ESG, orange box)) in beta-cells of 10-week-old *Ins*-Cre, *Pitpna^flox/flox^* and WT mice (n=8). **I**, Immunostaining of insulin and glucagon (Gcg) in paraffin-embedded pancreata from 10-week-old *Ins*-Cre, *Pitpna^flox/flox^* and WT mice. Scale bar= 100μm. In far-right panel, scale bar= 50μm. **J**, Islet morphometric analysis including islet number per area pancreas (mm^2^), insulin^+^ cells per area pancreas and pancreatic beta-cell mass in 10-week old *Ins*-Cre, *Pitpna^flox/flox^* and WT mice (n=5). Results presented as mean ± SEM. \**P*<0.05, \*\**P*<0.01, \*\*\**P*<0.001, and n.s. denotes not significant. See also Figure S3.

**Figure 3.**
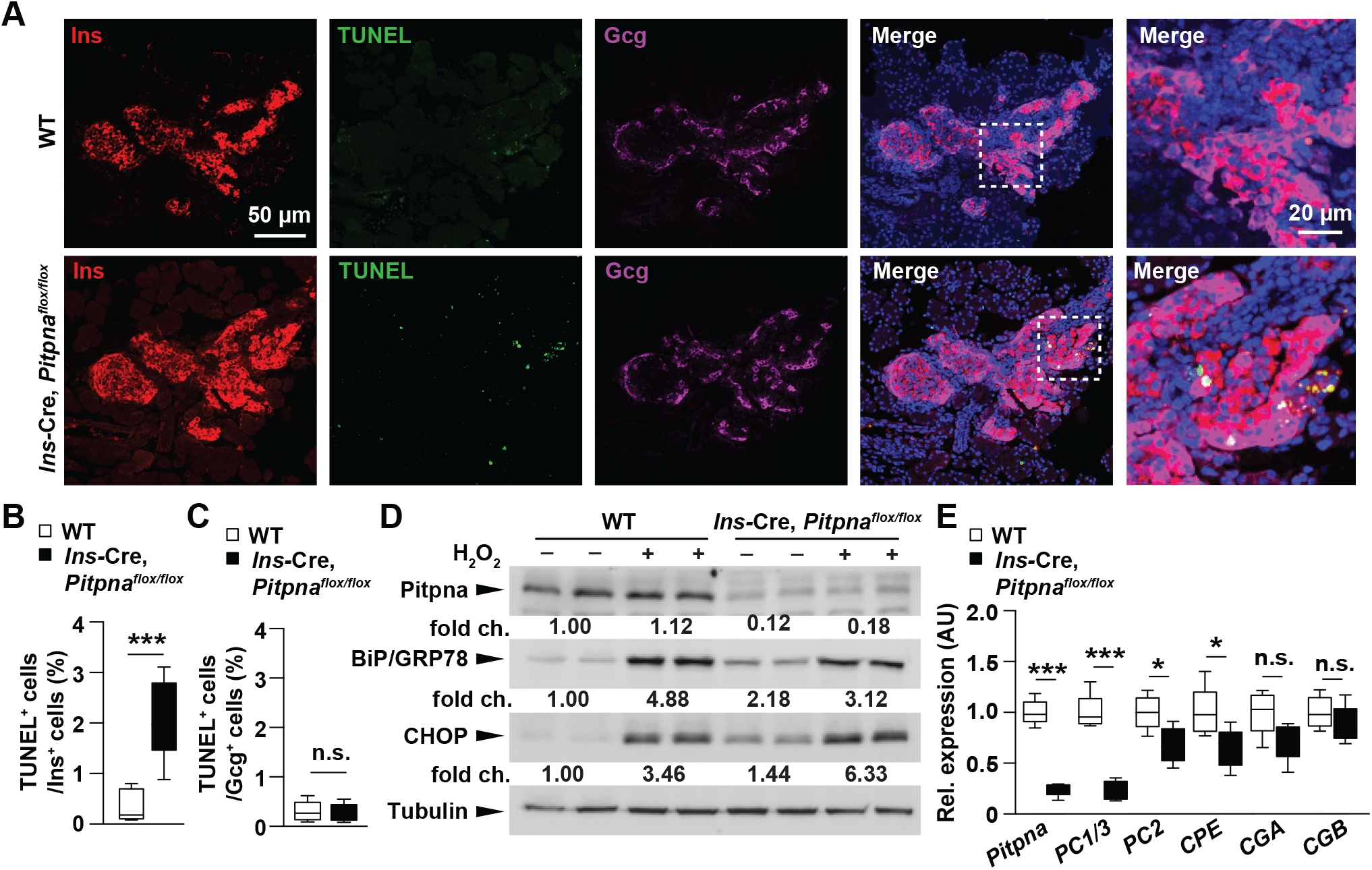
Loss of *Pitpna* increases beta-cell apoptosis and expression of endoplasmic reticulum (ER) stress markers. Immunostaining in paraffin-embedded pancreata from 10-week-old *Ins*-Cre, *Pitpna^flox/flox^* and littermate control wild-type (WT) mice was performed to assess: **A**, insulin (red), glucagon (Gcg, magenta) and apoptotic marker (TUNEL, green) Scale bar= 50μm. In far-right panel, scale bar= 20μm, **B**, TUNEL-positive beta cell number (n=6), and **C**, TUNEL-positive alpha cell number (n=6). **D**, Western blot analysis of Pitpna, BiP/GRP78, and CHOP after treatment of hydrogen peroxide (H_2_O_2_) in isolated islets of 10-week-old *Ins*-Cre, *Pitpna^flox/flox^* and WT mice. **E**, qRT-PCR analysis of *Pitpna, PC1/3, PC2, CPE, CGA* and *CGB* mRNA expression in islets of WT and *Ins*-Cre, *Pitpna^flox/flox^* mice at age 10 weeks (n=5). Results presented as mean ± SEM. \**P*< 0.05, \*\*\**P*<0.001, and n.s. denotes not significant.

### Endoplasmic reticulum (ER) stress in *Pitpna*-deficient beta-cells

Previous analysis in murine *Pitpna* null embryonic fibroblasts showed increased expression of the ER stress marker C/EBP homologous protein CHOP (Alb *et al*., 2003). Since elevated CHOP levels are consistent with ER stress-induced apoptosis (Harding and Ron, 2002), CHOP expression was examined in isolated islets of conditional *Pitpna* knockout mice. Indeed, we observed elevated basal CHOP levels at steady state and upon challenge of beta-cells with hydrogen peroxide -- an established inducer of oxidative and ER stress-mediated apoptosis (Figure 3D) (Back and Kaufman, 2012; Wright et al., 2013). Furthermore, steady-state levels of the unfolded protein response regulator GRP78/BiP were also increased ~2-fold in *Pitpna* null islets. These observations confirm that *Pitpna*-deficient beta-cells experience elevated chronic ER stress. We considered the possibility that impaired insulin granule synthesis, maturation, and exocytosis feeds back to induce ER stress as a result of continued high-level production of proinsulin in the face of a TGN trafficking ‘bottleneck’. With regard to insulin processing, qRT-PCR analysis of the insulin processing enzymes in isolated islets of *Ins*-Cre, *Pitpna^flox/flox^* mice revealed significant reductions in islet expression of *Proprotein convertase-1* (*PC1/3*), *Proprotein convertase-2* (*PC2*), and *Carboxypeptidase E* (*CPE*) (Figure 3E). These collective results report that Pitpna signaling is an essential component for maintaining beta-cell homeostasis. Chronic impairment of granule formation, maturation, and docking consequently triggers a cascade of ER stress and ultimately apoptosis. These defects represent the basis for the hyperglycemia observed in *Ins*-Cre, *Pitpna^flox/flox^* mice and illustrate the critical roles that this lipid transfer protein executes in the beta-cell.

### Pitpna regulates mitochondrial morphology in pancreatic beta-cells

The ER and the mitochondria engage in close physical contacts that are dynamic and are components of an inter-organelle communication system that responds to the metabolic demands of the cell (Tabara et al., 2021). In that regard, we observed that Pitpna-deficiencies impact mitochondrial function as reported by oxygen consumption rates (OCR) in MIN6 cells. *Pitpna* deficiencies attenuated both basal and maximal cellular respiration (Figures S4A, B), and suppressed glycolytic turnover and capacity as reported by the lowered extracellular acidification rates (ECAR) exhibited by cells silenced for *Pitpna* expression (Figures S4C, D). These observations are congruent with previous studies showing that brain and liver lysates from *Pitpna* whole body knockout mice exhibit dramatically reduced total ATP and ATP/ADP ratios (Alb *et al*., 2003). That is, phenotypes also consistent with reduced mitochondrial activity (Haythorne et al., 2019).

The derangements in mitochondrial morphology observed in *Pitpna*-deficient MIN6 cells translated to the animal model. Analyses of mitochondrial morphology in beta-cells of *Ins*-Cre, *Pitpna^flox/flox^* mice established that mitochondria were markedly longer in those Pitpna-deficient beta-cells, and that the frequencies of swollen mitochondria were also significantly increased relative to littermate control animals (Figures S4E-H). Previous studies demonstrate the guanosine triphosphatase (GTPase) Dynamin-related protein 1 (Drp1) is recruited to MERCs where its oligomerization enhances mitochondria constriction and fission (Friedman et al., 2011; Smirnova et al., 2001), and that deletion of Drp1 in beta-cells results in impaired GSIS (Hennings et al., 2018). Consistent with those findings, Drp1 expression was significantly diminished in isolated islets of *Ins*-Cre, *Pitpna^flox/flox^* mice (Figure S4I).

### PITPNA regulates human pancreatic beta-cell function

To experimentally assess whether PITPNA is a physiologically relevant regulator of human beta-cell function, both loss and gain-of-function approaches were taken in isolated islets of non-diabetic (ND) human donors. Implementing lentiviral constructs encoding either an shRNA (which encodes an siRNA targeting the human *PITPNA* mRNA sequence) or the human *PITPNA* full length cDNA, both knockdown (sh-*PITPNA*) and over-expression (OE-*PITPNA)* conditions were validated by immunoblotting methods and qRT-PCR (Figures 4A and S5A). GSIS was subsequently assessed in isolated human islets of ND donors after challenge with either sh-*PITPNA* or *OE-PITPNA* lentiviral vectors or a control lentivirus encoding a non-targeting shRNA vector (sh-*Ctrl*). Consistent with results from isolated islets of *Ins*-Cre, *Pitpna^flox/flox^* mice, lentiviral-mediated *PITPNA* knockdown inhibited insulin secretion upon stimulation with 25mM glucose, while GSIS was significantly elevated in the *PITPNA* over-expression condition relative to mock treated islets (Figure 4B). In addition, intracellular [Ca^2+^]_i_ was diminished in response to a 30mM KCl stimulus in islets of ND donors where *PITPNA* expression was inhibited. Those data support a model where the primary PITPNA execution point lies downstream of K^+^ channel closure – i.e. at the level of granule trafficking, docking and/or exocytosis (Figure 4C).

**Figure 4.**
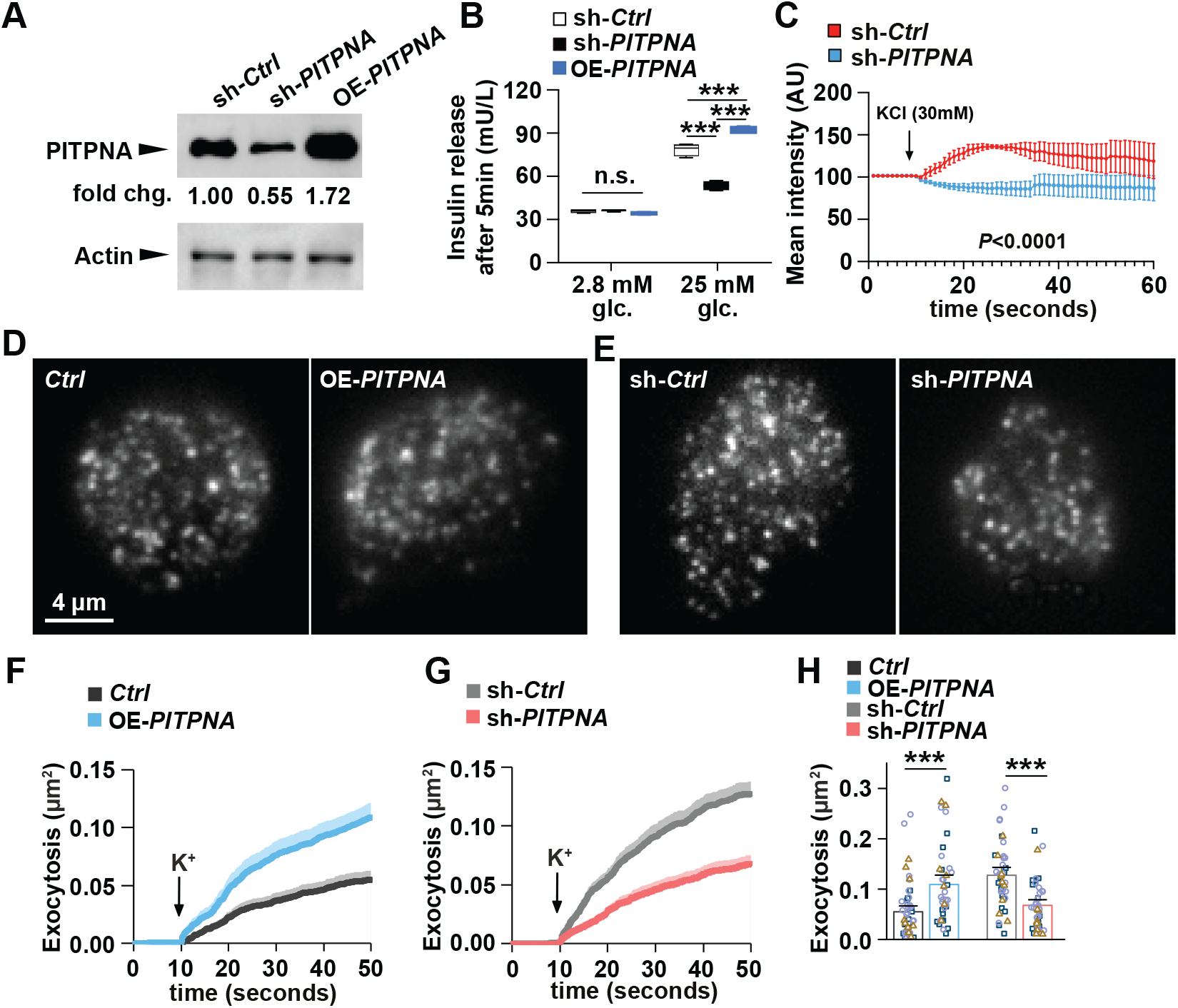
*PITPNA* regulates insulin secretion in human pancreatic beta-cells. **A**, Western blot analysis of PITPNA in isolated islets from non-diabetic human donors after treatment with lentiviruses encoding either an shRNA targeting *PITPNA* (sh-*PITPNA*), cDNA of human *PITPNA (OE-PITPNA)*, or empty control vector (sh-*Ctrl*). **B.** Quantification of insulin release in response to 2.8mM and 25mM glucose concentrations from isolated islets from non-diabetic human subjects after lentiviral-mediated over-expression of *PITPNA* (OE-*PITPNA*) or inhibition of *PITPNA* (sh-*PITPNA*) in comparison to treatment with control lentivirus (sh-*Ctrl*) (n=4). **C**, Quantification of intracellular Ca^2+^ concentration in isolated human islets after lentiviral-mediated inhibition of *PITPNA (sh-PITPNA)* in comparison to control lentivirus (sh-*Ctrl*) (n=5). **D**, **E**, Representative TIRF images of human beta-cells expressing the granule marker NPY-tdmOrange2 after treatment with lentiviruses encoding GFP control (*Ctrl*), cDNA of human *PITPNA* (OE-*PITPNA*) in panel (**D**), as well as an shRNA pool targeting *PITPNA* (sh-*PITPNA*) or shRNA control (sh-*Ctrl*) in panel (**E**); Scale bar, 4 μm. **F**, **G**, Cumulative time course of high K^+^-evoked exocytosis events normalized to cell area, for conditions as in **D**, **E**. Bars at individual time points indicate SEM., K^+^ was elevated to 75mM during t=10-50 seconds. **H**, Total exocytosis measured during TIRF analysis of human beta-cells expressing the granule marker NPY-tdmOrange2 after lentivirus treatments represented in panels (**D**) and (**E**). Data set was generated from 3 unique human donors; dots and their color/symbol indicate individual cells and donor, respectively. Data presented as mean± SEM, unless otherwise indicated. \*\*\**P*<0.001, and n.s. denotes not significant. See also Figure S4.

To interrogate how PITPNA affects stimulus-secretion coupling, total internal reflection (TIRF) microscopy was used to monitor exocytosis and docking of insulin granules in dispersed human islet cells. After plating, islet cells were treated with either sh-*PITPNA* or OE-*PITPNA* lentiviral vectors or their respective control lentivirus (sh-*Ctrl* or an empty vector control, *Ctrl*) (Figures 4D and E). In addition, a genetically encoded NPY-tdmOrange2 marker was used to label granules (Figures 4D, E). Exocytosis was evoked by depolarization with elevated K^+^ (in the presence of diazoxide to prevent spontaneous depolarization) and it followed a biphasic time course (Figures 4F and G). Exocytosis was increased in the face of PITPNA overexpression (by 98% vs *Ctrl* cells; P=0.0004 non-paired t-test; 50 *Ctrl* cells and 40 *OE-PITPNA* cells; 3 donors each; Figures 4F and H), and decreased by *PITPNA* silencing (by 47% vs sh-*Ctrl*; P=1E-05 non-paired t-test; 40 sh-*Ctrl* cells and 43 sh-*PITPNA* cells; 3 donors each; Figures 4G and 4H). These data indicate a positive correlation between PITPNA expression and exocytosis in human beta cells and, from these observations, we conclude that PITPNA-dependent changes in exocytosis reflect changes in the secretory machinery of individual insulin granules. Electron microscopy imaging reported that insulin granule core density and numbers of docked vesicles were significantly reduced after *PITPNA* knockdown in isolated human ND islets relative to control lentivirus-treated islets (Figures 5A-C). Moreover, shRNA-mediated silencing of *PITPNA* impaired the formation of mature secretory granules (MSG) with a reciprocal increase in the numbers of immature secretory granules (ISG) (Figure 5D). Conversely, *PITPNA* over-expression in isolated ND islets increased MSG numbers with associated reductions in ISG numbers (Figure 5D). These results demonstrate that PITPNA is a potent regulator of granule maturation and docking in human beta-cells.

**Figure 5.**
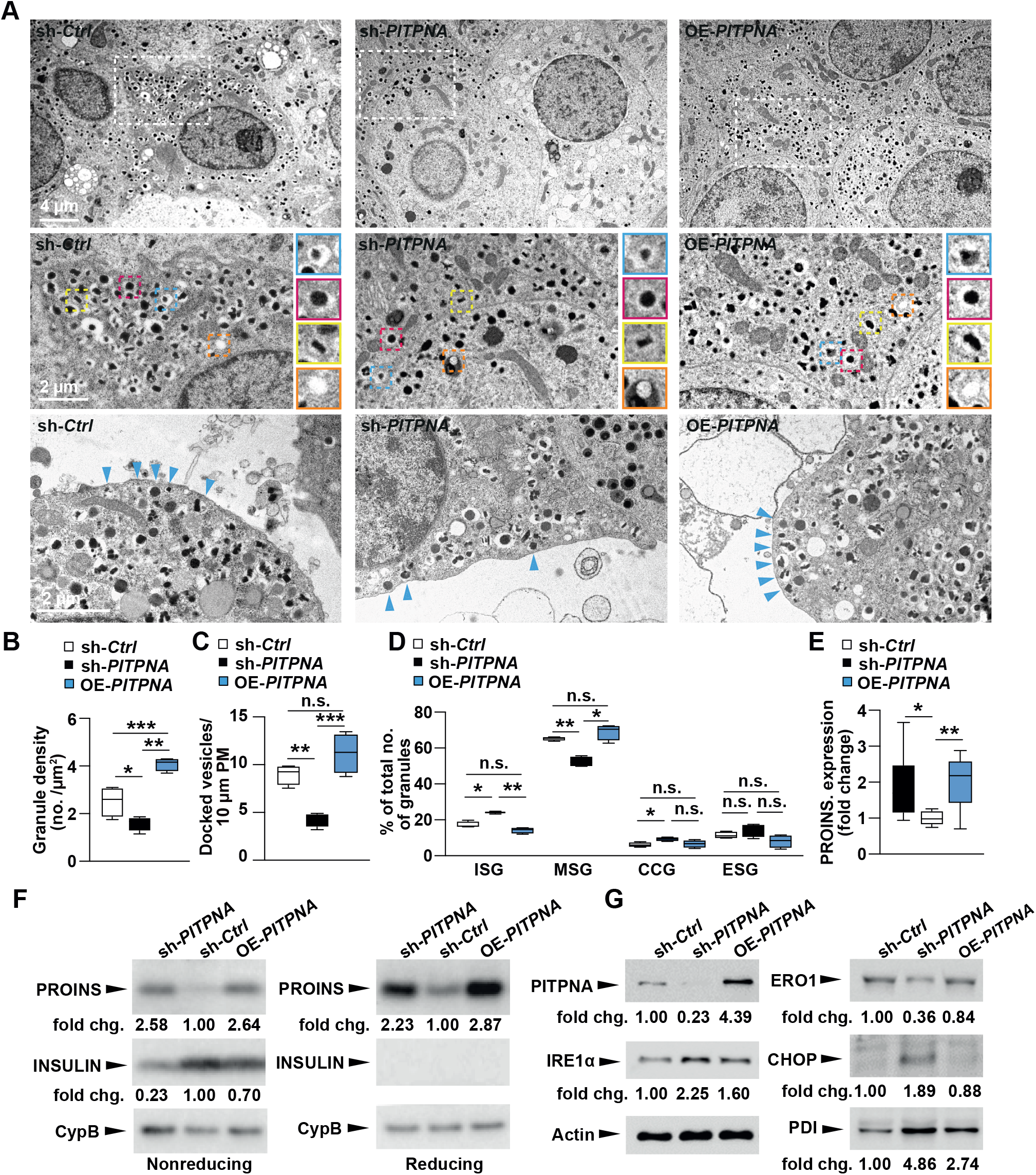
*PITPNA* regulates insulin granule maturation and proinsulin processing in human pancreatic beta-cells. **A**, Representative transmission electron micrographs of pancreatic beta-cells from non-diabetic human donors after lentiviral-mediated over-expression of *PITPNA* (OE-*PITPNA*) or inhibition of *PITPNA* (sh-*PITPNA*) in comparison to control lentivirus (*sh-Ctrl*); granule profile: immature secretory granule (blue box), mature secretory granules (red box), crystal-containing granules (yellow box), and empty secretory granules (orange box). **B**, **C**, Quantification of granule density and docked vesicles in beta-cells of lentiviral*-*treated human islets shown in panel (**A**) (n=4). **D**, Quantification of immature secretory granule (ISG), mature secretory granules (MSG), crystal-containing granules (CCG), and empty secretory granules (ESG) in beta-cells of isolated human islets after lentiviral treatments shown in panel (**A**) (n=8-9). **E**, **F**, Quantification of proinsulin in isolated human islets after densitometric analysis of western blots shown in panel (**F**). **G,** Western blot analysis of PITPNA, and ER stress/unfolded protein response (UPR) proteins IRE1α, ERO1, PDI, and CHOP in human islets after lentiviral-mediated over-expression of *PITPNA* (OE-*PITPNA*), knockdown of *PITPNA* (sh-*PITPNA*) or control lentivirus (sh-*Ctrl*). Results presented as mean ± SEM. \**P*< 0.05; \*\**P*– 0.01; \*\*\**p*<0.001, and n.s. denotes not significant. See also Figure S4.

The collective insulin granule data collected in both human and mouse loss-of-function studies suggested PITPNA insufficiencies in human beta-cells ultimately disrupt proinsulin packaging into insulin granules. Indeed, in a manner consistent with the results from *Ins*-Cre, *Pitpna^flox/flox^* mice, proinsulin levels were elevated upon *PITPNA* silencing (sh-*PITPNA*) in isolated human ND islets (Figures 5E, F). Reciprocally, islet insulin levels were reduced relative to control lentivirus-treated human ND islets (sh-*Ctrl*) -- further reporting proinsulin processing is impaired upon loss of PITPNA activity. By contrast, increasing PITPNA expression in islets (OE-*PITPNA*) elevated both proinsulin and insulin levels compared to control-treated human ND islets (sh-*Ctrl*). These results demonstrate that the enhanced GSIS supported by increased PITPNA activity is supported by increased granule maturation and proinsulin processing (Figures 5E, F). Previous studies highlighted an association of proinsulin accumulation with perturbed expression of UPR/ER stress proteins (Arunagiri et al., 2018). Our findings with *Ins*-Cre, *Pitpna^flox/flox^* mice (Figure 3D) prompted examination of whether proinsulin accumulation in isolated ND human islets induces ER stress. Immunoblot analyses confirmed *PITPNA* silencing in ND human islets (sh-*PITPNA*) resulted in increased expression of CHOP as well as other components of the ER stress pathway -- including inositol-requiring enzyme 1 alpha (IRE1a) and protein disulfide isomerase-a1 (PDI) (Figure 5G). By contrast, protein disulfide oxidase ER-Oxidoreducin 1 alpha (ERO1) steady-state levels were decreased when *PITPNA* was silenced in ND human islets. These collective data report that expression levels of multiple components of the ER stress pathway are perturbed under conditions of PITPNA insufficiency (Figure 5G). Moreover, these collective data demonstrate that loss of *PITPNA* results in similar derangements in both human and mouse beta-cell systems. These include impaired granule biogenesis and maturation that is accompanied by increased islet expression of multiple ER stress markers such as CHOP -- a member of the C/EBP family of transcription factors linked to programmed cell death (Marciniak et al., 2004; Song et al., 2008; Zinszner et al., 1998).

The translation of phenotypes associated with PITPNA deficiencies from murine to human beta-cells extended to mitochondrial dysmorphologies. *PITPNA* silencing in ND human beta-cells resulted in lengthening of mitochondrial ribbons while *PITPNA* over-expression markedly shifted the morphological distribution to shorter mitochondria (Figures S5B, C). *PITPNA* silencing in human beta-cells also reduced the number of morphologically ‘orthodox’ mitochondria with proportional increases in the frequencies of ‘swollen’ mitochondria (Figure S5D). These results demonstrate that PITPNA in human beta-cells potently regulates insulin exocytosis, intracellular Ca^2+^ concentrations, granule maturation and docking, and mitochondrial morphology. The data further project that diminished PITPNA expression during the course of T2D is a plausible contributor to beta-cell failure.

### PITPNA mediates PtdIns-4-P synthesis in human pancreatic islets

The available data indicate PITPNA stimulates PtdIns 4-OH kinases by using its lipid-exchange activity to render PtdIns a better substrate for the enzyme. It is in this way that PITPNA promotes PtdIns-4-P synthesis (Bankaitis et al., 2012; Grabon et al., 2019; Xie *et al*., 2018). To test whether reduction of *PITPNA* expression in human islets attenuates formation of PtdIns-4-P, we again performed shRNA-mediated silencing of *PITPNA* in isolated islets of non-diabetic donors. After 48 hours of incubation, PtdIns-4-P status was assessed using immunohistochemical methods (Figure 6A). The PtdIns-4-P signal in insulin^+^ beta-cells was significantly diminished -- indicating lentiviral-mediated knockdown of *PITPNA* significantly reduced cellular levels of this phosphoinositide (Figure 6B). These results were supported by an independent assay monitoring GOLPH3 localization as this protein is recruited to TGN membranes by virtue of its ability to bind PtdIns-4-P (Kuna and Field, 2019) (Xie *et al*., 2018). *PITPNA* silencing in isolated human islets evoked release of GOLPH3 from TGN membranes as evidenced by the significant reductions in GOLPH3 co-localization with the TGN marker GOLGIN97 (Figures 6C, D).

**Figure 6.**
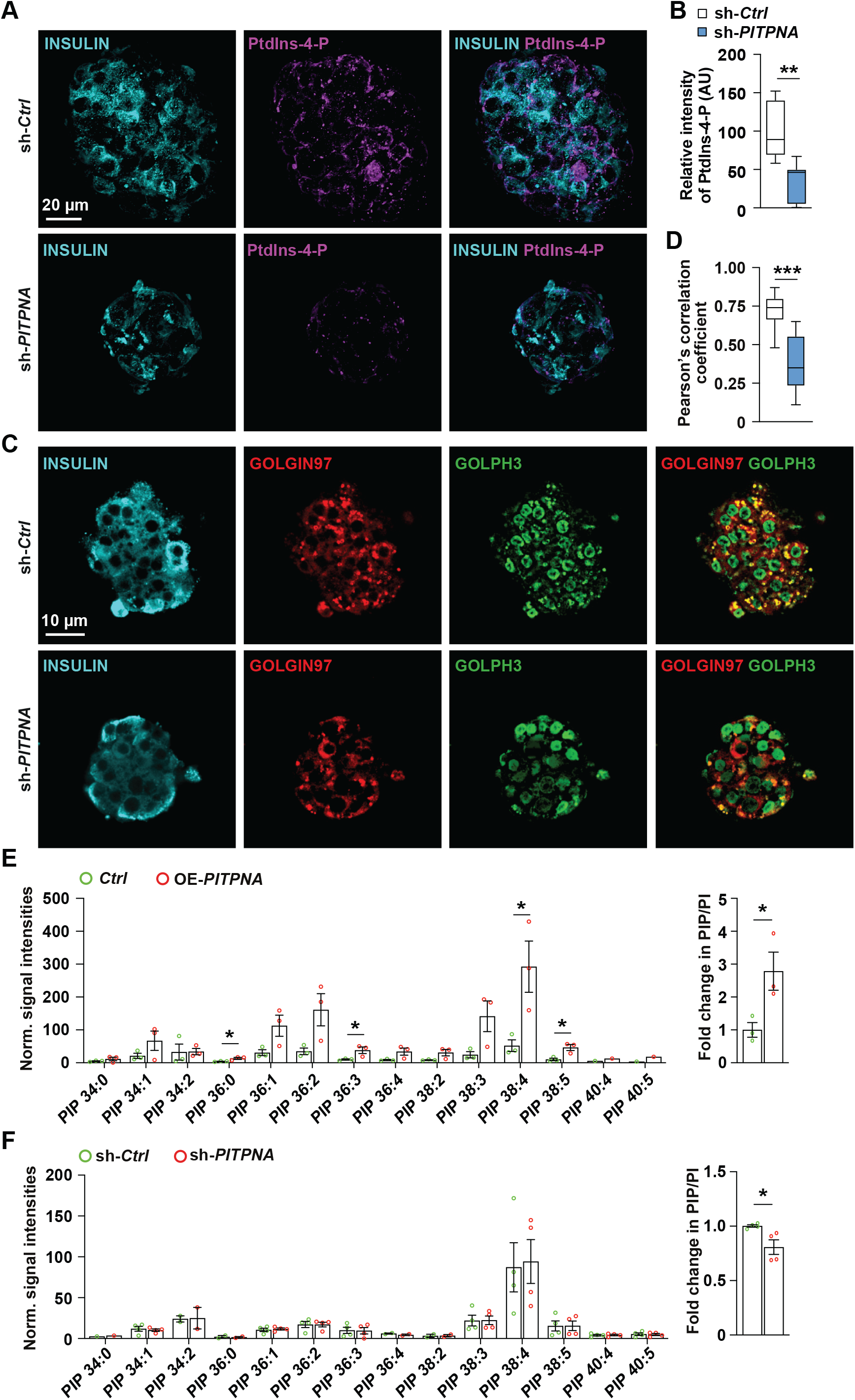
Inhibition of *PITPNA* in isolated human islets disrupts subcellular localization of PtdIns-4-P to the TGN. **A**, **B** Immunostaining for INSULIN and Ptdlns-4-P, and quantification of the intensity of Ptdlns-4-P in isolated human islets after lentiviral-mediated inhibition of *PITPNA* (sh-*PITPNA*) or treatment with control lentivirus (sh-*Ctrl*) (n=5-7). Scale bar = 20μm. **C**, **D** Quantification of GOLGIN97 and GOLPH3 colocalization after immunostaining of isolated human islets after lentiviral-mediated inhibition of *PITPNA* (sh-*PITPNA*) (n=10) or treatment with control lentivirus (sh-*Ctrl*) (n=23). **E**, Quantification of phosphatidylinositol-phosphate (PtdIns or PIP) species in isolated human islets after lentiviral-mediated over-expression of *PITPNA (OE-PITPNA)* or treatment with a control lentivirus (*Ctrl*) (n=3). Results normalized to total cellular PtdIns (PI). **F**, Quantification of phosphatidylinositol-phosphate (PtdIns or PIP) species in isolated human islets after lentiviral-mediated inhibition of *PITPNA* (sh-*PITPNA*) or treatment with a lentivirus expressing an shRNA control (sh-*Ctrl*) (n=4) and normalized to total cellular PtdIns (PI). Results presented as mean ± SEM. \**P*< 0.05, \*\**P*< 0.01, \*\*\**P*<0.001. See also Figure S5.

To further test whether reduced *PITPNA* expression in human islets attenuates PtdIns-4-P synthesis, *PITPNA* was either silenced or over-expressed in isolated islets of ND human donors. Mass spectrometry-based quantitative lipidomics were then performed to measure bulk cellular PtdIns and PtdIns-P levels. Although mass spectrometry cannot distinguish regio-isomers; PtdIns-4-P is the most abundant isomer in mammalian cells and PtdIns −4-P is estimated to constitute >90% of total cellular PtdIns-P (Hammond et al., 2012; Stephens et al., 1993). After normalization of PtdIns-P levels to total cellular PtdIns, the data demonstrate that *PITPNA* over-expression (*PITPNA-OE*) in isolated islets of ND human donors increased PtdIns-P levels compared to mock controls (*Ctrl*) (Figures 6E and S5E). Conversely, *PITPNA* silencing in isolated human islets decreased cellular PtdIns-P levels relative to control islets (Figures 6F and S5F). These observations demonstrate that modulation of *PITPNA* in isolated human islets impacts PtdIns-4-P homeostasis, and are consistent with studies in mammalian neural stem cells (Xie *et al*., 2018).

### Restoration of PITPNA rescues beta-cell function in T2D islets

The weight of the collective data gleaned from both murine animal models and human islets, including the demonstration that *PITPNA* expression was diminished in pancreatic beta-cells of T2D human subjects, implicates PITPNA as a major factor in beta-cell failure during T2D. These aggregate results raised the provocative question of whether recovery of PITPNA expression in T2D islets restores function to the diseased tissue. Indeed, lentiviral-mediated induction of *PITPNA* in isolated islets from T2D human donors significantly elevated PITPNA expression levels in islets of the T2D donor (T2D-*PITPNA* OE), and these levels were comparable to the endogenous expression levels recorded for islets of the ND human donor (Non) (Figure 7A). Strikingly, the rescue of *PITPNA* expression in T2D islets significantly improved GSIS in response to 15mM glucose compared to control treated T2D islets (Figures 7B and S6A). Moreover, recovery of PITPNA expression in T2D islets rescued PtdIns-4-P synthesis as evidenced by PITPNA inducing redistribution of GOLPH3 from a dispersed cytoplasmic localization to TGN membranes marked by GOLGIN97 (Figures 7C, D).

**Figure 7.**
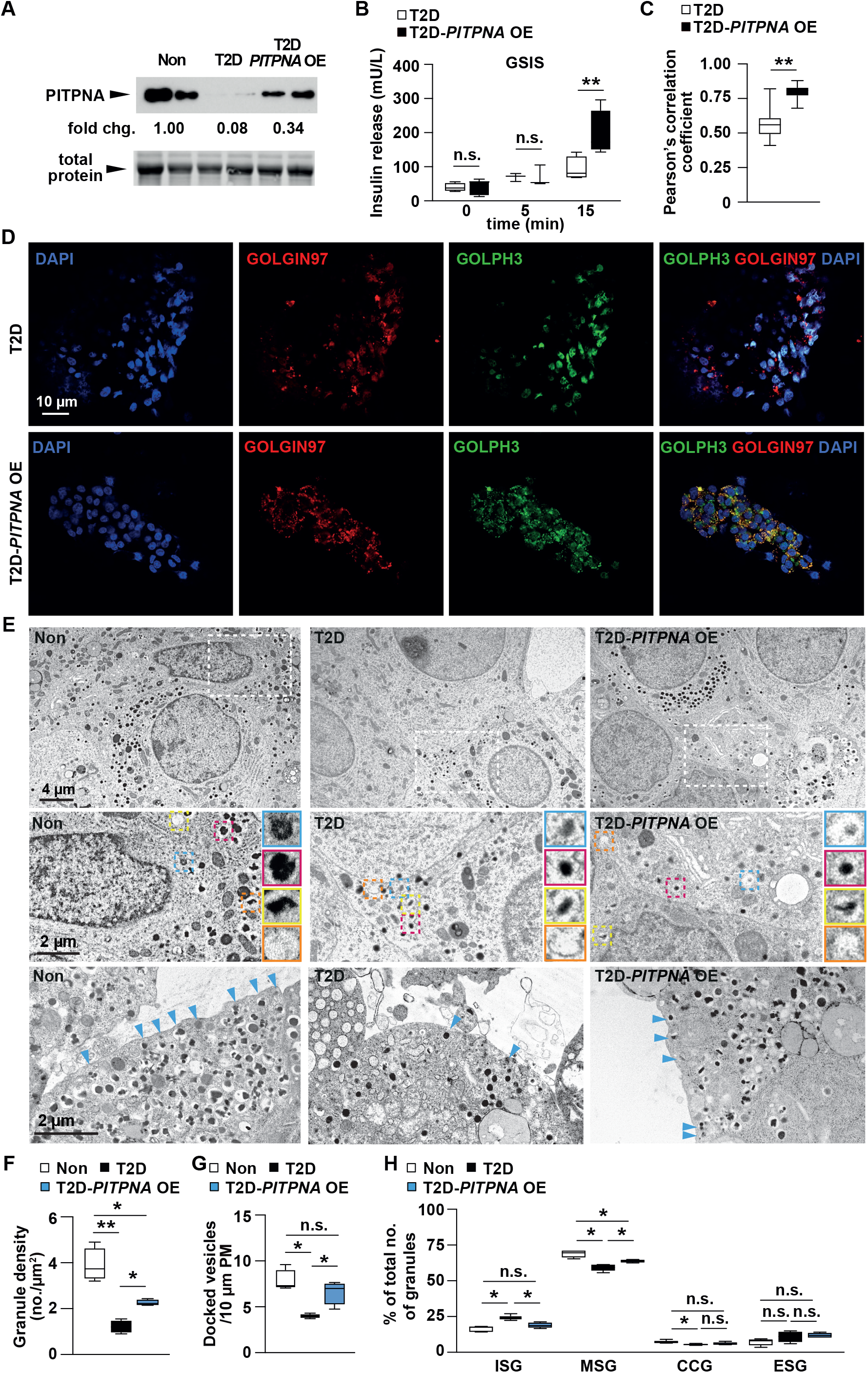
Restoration of*PITPNA* in isolated islets of T2D human subjects rescues pancreatic beta-cell function. **A**, Western blot analysis of PITPNA in human non-diabetic (Non), and T2D islets after either lentiviral-mediated over-expression of *PITPNA (T2D-PITPNA* OE) or treatment with a control lentivirus (T2D) (n=2). **B,** Quantification of insulin release from isolated islets from T2D donors after either lentiviral-mediated over-expression of *PITPNA* (T2D-*PITPNA* OE) or treatment with a control lentivirus (T2D) (n=4). **C, D** Quantification of GOLGIN97 and GOLPH3 colocalization after immunostaining of isolated human islets from T2D donors after either lentiviral-mediated over-expression of *PITPNA* (T2D-*PITPNA* OE) or treatment with a control lentivirus (T2D (n=15). Scale bar = 10μm. **E**, Representative transmission electron micrographs of pancreatic beta-cells of non-diabetic (ND) or T2D human donors after treatment with a control lentivirus (T2D) or lentiviral-mediated over-expression of *PITPNA* (T2D-*PITPNA* OE); immature secretory granule (blue box), mature secretory granules (red box), crystal-containing granules (yellow box), and empty secretory granules (orange box). Quantification of **F**, granule density, **G**, docked vesicles, and **H**, granule profile: immature secretory granule (ISG), mature secretory granules (MSG), crystal-containing granules (CCG), and empty secretory granules (ESG) in beta-cells of non-diabetic (ND) or T2D human donors after treatment with a control lentivirus (T2D) or lentiviral-mediated over-expression of *PITPNA* (T2D-*PITPNA* OE) (n=4). Results presented as mean ± SEM. \**P*< 0.05; \*\**P*< 0.01; \*\*\**p*< 0.001, and n.s. denotes not significant. See also Figure S6.

Restoration of PITPNA expression to T2D islets exhibited other profound effects. Electron microscopy analyses indicated insulin granule number, docking and maturation were rescued upon induction of *PITPNA* expression in T2D islets (Figure 7E). Notably, insulin granule number per μm^2^ was significantly lower in T2D islets compared to granule density in non-diabetic islets. Recovery of *PITPNA* expression in T2D beta-cells markedly rescued the reduction in granule number (Figure 7F), fully rescued the granule docking defects in T2D beta-cells (Figure 7G), and effected a partial rescue of mature granule numbers (Figure 7H). Additionally, restoration of PITPNA expression in islets of four individual T2D human donors (T2D-OE) resulted in the downregulation of steady-state levels of CHOP, PDI, and BiP/GRP78 (Figure S6B) as well as restoration of proinsulin expression (Figure S6C). Taken together, these results demonstrate that restoration of *PITPNA* expression in T2D beta-cells substantially reverses the GSIS defects, the impaired insulin granule biogenesis and maturation, and the chronic ER stress associated with human T2D.

## DISCUSSION

Critical to the development of therapeutics for diabetes are strategies for promoting insulin release while preserving pancreatic beta-cell mass. Recent studies focus on defects in insulin processing and granule maturation as causes for reduced insulin secretion that are linked to all major forms of diabetes (Campbell and Newgard, 2021; Liu *et al*., 2021) – a focus that rests on demonstrations that: 1) glucose-dependent granule docking is a limiting factor for insulin secretion and 2) reduced granule docking characterizes beta-cell dysfunction during human T2D (Gandasi et al., 2018). In this study, we demonstrate that reduced PITPNA-dependent PtdIns-4-P signaling in the beta cell TGN results in beta-cell failure. We show that PITPNA deficiencies impair insulin granule maturation and exocytosis, and that these trafficking defects induce proinsulin accumulation, promote chronic ER stress, and derange mitochondrial dynamics and performance. The data outline a high degree of functional dependence between the TGN, ER and mitochondria, and identify PITPNA as a central regulator of this intra-organelle crosstalk. Finally, we report the remarkable demonstration that restoring PITPNA expression to T2D human islets is sufficient to reverse beta-cell failure by rescuing GSIS, insulin granule maturation, proinsulin processing, and by alleviating the chronic ER stress that accompanies these defects in T2D beta-cells. These results: (i) highlight PITPNA-dependent PtdIns-4-P synthesis on TGN membranes as critical for sustaining insulin granule biogenesis and maturation, (ii) indicate compromise of this activity is a powerful marker of beta-cell failure during T2D, and (iii) identify new prospects for T2D therapy.

All available in vivo evidence, collected from single cell yeast to mammalian models, indicates that soluble PITPs potentiate constitutive membrane trafficking from late compartments of the secretory pathway – specifically TGN/endosomes. Analyses of headgroup-specific PITP mutants and localization of PtdIns-4-P biosensors indicate the biochemical basis for PITP function is to stimulate PtdIns-4-P synthesis on TGN/endosomal membranes with the result that PtdIns-4-P binding proteins (i.e. effectors of PtdIns-4-P signaling) are recruited to these compartments (Alb et al., 1995; Alb et al., 2007; Bankaitis *et al*., 1990; Bankaitis *et al*., 2010; Hay and Martin, 1993; Lete *et al*., 2020; Ohashi *et al*., 1995; Schaaf *et al*., 2008; Xie and Bankaitis, 2022; Xie *et al*., 2018). It is in this fashion that PtdIns-4-P is proposed to act as a transient tag to convey spatial information that helps organize membrane trafficking (Balla, 2013; Behnia and Munro, 2005). The current demonstration that PITPNA is required for insulin granule formation, maturation and exocytosis now extends this concept to regulated membrane trafficking pathways in human pancreatic beta-cells. This conclusion is further supported by: (i) the demonstration that modulation of *PITPNA* in human beta-cells regulates PtdIns-P, (ii) the Sac2 phosphatase is a PtdIns-4-P binding protein that localizes to the insulin granule surface where it mediates granule docking to the plasma membrane and exocytosis (Nguyen et al., 2019), and (iii) measurements reporting that PtdIns (the direct metabolic precursor of PtdIns-4-P) constitutes ~21% of insulin granule lipid in the INS-1 832/13 beta-cell line (MacDonald et al., 2015). While the precise role(s) of PtdIns-4-P in granule docking and exocytosis remains to be fully clarified, the demonstration that dephosphorylation of PtdIns-4-P by the phosphatase Sac2 disrupts insulin granule docking and GSIS, and that Sac2 expression is decreased in T2D islets alludes to its functional significance (Nguyen *et al*., 2019; Omar-Hmeadi and Idevall-Hagren, 2021). We suggest that Sac2-mediated dephosphorylation of PtdIns-4-P ‘signals’ the end of the insulin granule biogenesis/maturation phase, and ‘identifies’ the mature granule as competent for mobilization to the plasma membrane for docking and exocytosis.

A striking consequence associated with PITPNA inhibition in human beta-cells is the potent increase in proinsulin levels. Initial accumulation of proinsulin correlates with a stressed ER in islets of *Lepr^db/db^* mice as blood glucose levels rise (~237 mg/dL), and is maintained until proinsulin levels dramatically fall upon onset of severe hyperglycemia (~523 mg/dL) (Arunagiri et al., 2019). Our demonstration that PITPNA levels are significantly reduced in T2D islets compared to expression in islets of non-diabetic controls, and that restoring PITPNA expression to the beta-cell helps to recover proinsulin expression, agree with those previous findings. The accumulation of proinsulin detected after acute inhibition of *PITPNA* in human islets may reflect the impaired granule formation and/or maturation at an early stage of dysfunction, that persists until chronic insulin demand and ER stress cause the beta-cell to cease proinsulin production leading to hyperglycemia. Moreover, the accumulation of proinsulin after acute inhibition of *PITPNA* shows downstream defects in granule maturation and docking and GSIS are ultimately linked to induce ER stress (Sun et al., 2015). Activation of the ER stress pathway might be directly related to adverse changes in mitochondrial or ER dynamics (Fonseca *et al*., 2011; Harding and Ron, 2002), or in activation of an inter-organellar response that negatively feeds back on proinsulin export from the ER (Mousley et al., 2008).

The perturbations in mitochondrial performance and health of the endoplasmic reticulum in PITPNA-deficient beta-cells are notable. PtdIns-4-P is present on the surface of TGN-derived vesicles recruited to MERCs, and this PtdIns-4-P pool is reported to aid in potentiation of mitochondrial fission and ER dynamics (Liesa and Shirihai, 2013; Mishra and Chan, 2014; Nagashima et al., 2020; Tabara *et al*., 2021; Youle and van der Bliek, 2012). As PITPNA promotes PtdIns-4-P synthesis in the mammalian TGN by facilitating presentation of PtdIns to PtdIns 4-OH kinases (Xie *et al*., 2018), we suggest that the PITPNA-regulated PtdIns-4-P pool in beta-cells the coordinates actions of the TGN in ER/mitochondrial dynamics in addition to facilitating insulin granule biogenesis. It is presently thought that mitochondrial fission is essential for sustaining a healthy pool of mitochondria by allowing for the clearance of damaged mitochondria through mitophagy and de novo biogenesis (Bock and Tait, 2020; Youle and van der Bliek, 2012). Moreover, mitophagy protects human pancreatic beta-cells from inflammatory damage during diabetes (Sidarala et al., 2020) -- indicating the removal of dysfunctional mitochondria is essential for preventing inflammatory stress and cell death. Our results showing mitochondrial lengthening as a consequence of functional ablation of *PITPNA* in both murine and human beta-cells suggests diminished PITPNA-dependent PtdIns-4-P synthesis impacts mitochondrial dynamics in the beta-cell. That *PITPNA* insufficiencies in human beta-cells induce accumulation of swollen mitochondria further emphasizes this point.

Our demonstration that *Pitpna* is a direct target of miR-375 shows that the complex relationship between the TGN, ER, and mitochondria is subject to regulation by the miRNA pathway. MiR-375 is the most abundant microRNA in the pancreatic beta-cell and is a potent regulator of insulin secretion and adaptive proliferation (Poy *et al*., 2004; Poy *et al*., 2009; Tattikota *et al*., 2014; Tattikota *et al*., 2013). Establishing an association between *Pitpna* and miR-375 suggests a framework for how the beta-cell exerts regulatory control over its critical functions such as granule maturation, exocytosis, and mitochondrial dynamics. We previously demonstrated how miR-375 targets (e.g. *Cadm1, Gphn, Elavl4* and *Mtpn)* regulate beta-cell secretion (Poy *et al*., 2009; Tattikota *et al*., 2013). Inclusion of *Pitpna* in this regulon amply illustrates the functional diversity of microRNA-targeted genes that mediate exocytosis. We posit that suppression of these genes by miR-375 provides broad regulatory control over the beta-cell secretory machinery and ‘secretome’ under normal steady state conditions and this circuit may prevent excess insulin release and safeguards the central nervous system from hypoglycemia (Poy, 2016). These findings reinforce the notion that the microRNA pathway is a critical component for how cells adapt to changes in their metabolic environment as well as demonstrate how disruption of this pathway renders the beta cell incapable of maintaining a proper homeostatic balance with the ultimate result of diabetic disease (LaPierre and Stoffel, 2017; Poy, 2016).

In summary, this study describes several important conceptual advances. These include: (i) establishment of PITPNA as a major regulator of PtdIns-4-P signaling in the TGN of human pancreatic beta-cells, (ii) demonstration that PITPNA is required for efficient insulin granule maturation, docking, secretion, and proinsulin processing in mammalian (including human) pancreatic beta-cells, and (iii) demonstration that restoration of PITPNA expression in human T2D beta-cells rescues insulin secretion, granule maturation and alleviates ER stress. These data not only highlight PITPNA deficiency as a major contributing factor to reduced insulin output and beta-cell failure, but also report a functional crosstalk between the miRNA pathway and lipid signaling control of membrane trafficking factors that are relevant to human diabetes. This study raises the intriguing prospect that enhancing PITPNA expression or activity in islets of T2D human subjects may rescue the multiple defects that contribute to beta-cell degeneration to the extent that physiologically significant activity is revived in the T2D pancreas.

## MATERIALS AND METHODS

### Human Islets

Human islets from non-diabetic (ND) and type 2 diabetic (T2D) subjects isolated from cadaveric pancreas were obtained from the Integrated Islet Distribution Program (IIDP), the University of Alberta IsletCore, Prodo Laboratories, and the Nordic Network for Clinical Islet Transplantation (Uppsala, Sweden) with permission from the Johns Hopkins Institutional Review Board (IRB00244487). Human islet cells were obtained from de-identified donors and all organ donors provided informed consent for use of human islets for research. Relevant donor information including age, gender, ethnicity, diabetes status and body mass index (BMI) are listed in Table S1. Diabetes status was determined from patient records and available hemoglobin A1c (HbA1c) data.

### Animals

Mice were maintained on a 12-hour light/dark cycle with ad libitum access to regular chow food (2016 Teklad global 16% diet, Envigo) and the Johns Hopkins Animal Care and Use Committee approved all experimental procedures under protocol MO18C281. Results were consistent in both genders; however, data from female mice are not shown. *Pitpna* whole-body knockout mice were previously described (Alb *et al*., 2003; Xie *et al*., 2018). *Pitpna*-floxed mice (VAB line) were generated using a *Pitpna*-floxed allele generated by TALEN-based methods and transplaced into C57BL/6 embryonic stem cells by homologous recombination. Details are available upon request. The successfully targeted *Pitpna* allele had a LoxP sequence inserted upstream of exon 8 and a neomycin cassette (flanked by Frt sequences)-LoxP sequence inserted downstream of exon 10. The neomycin cassette flanked by Frt sequences was removed by crossing to an FLP deleter strain. In the resulting strain (i.e. *Pitpna*-floxed strain), exons 8-10 of *Pitpna* were flanked by LoxP sequences. Deletion of exons 8-10 generates a *Pitpna* null allele. The primers for genotyping the *Pitpna*-floxed allele were: TAMU002_LoxP_F: 5’-AGTGAGTTCCAAAA TGGCCAGGTT-3’; and TAMU002_LoxP_R: 5’-GCCAGTTCTTTTTGTCGCTGTGAA-3’. The size of the PCR product was 242bp for the wild-type *Pitpna* allele, and 312bp for the floxed *Pitpna* allele. Floxed *Pitpna* mice were crossed with *Ins1-Cre* mice purchased from Jackson Labs (Thorens *et al*., 2015). Floxed *Ago2* mice were generated and crossed with *Ins-Cre* mice from P. Herrera as described (Tattikota *et al*., 2014). *Lep^ob/ob^* mice (cat. no: 000632) were purchased from Jackson Laboratories, Maine, USA. Numbers of animals are reported in each figure legends, and experiments were conducted in a blinded manner where the genotype is unknown during actual testing.

### Gene expression analysis in mouse and human islets

Total RNA was extracted using the TRIzol reagent (Invitrogen). Quantitative real time PCR (qRT-PCR) for *miR-375* was quantified by TaqMan Assays using the TaqMan MicroRNA Reverse Transcription Kit and hsa-miR-375 primer sets (Thermo Fisher Scientific, 000564). *MiR-375* levels were normalized to *miR-U6* expression. For the expression of gene mRNAs, cDNA was synthesized using RevertAid First Strand cDNA synthesis kit (Fermentas), and qRT-PCR was measured using gene-specific primers with FastStart SYBR Green PCR Master Mix (Roche) on a StepOne Real-Time PCR System (ThermoFisher). Gene expression analysis from cell lines and mouse and human islets was performed as described, primers used with FastStart SYBR Green PCR Master Mix (Roche) are described in Table S4. Human islet expression data and accompanying donor information were previously published (Fadista *et al*., 2014) and are publicly accessible at Gene Expression Omnibus (GEO accession number GSE50398). Briefly, RNA-seq data sets were downloaded, trimmed (TrimGalore) and mapped to GRCh38 (HISAT2 mapper) (Kim et al., 2015). Read counts for each sample were generated in SeqMonk software and normalised. The expression levels for *PITPNA* and *INSULIN* were correlated to the published clinical data included with the GEO submission. The single cell RNA-seq data (GEO accession number GSE85241) (Muraro *et al*., 2016) was downloaded from https://hemberg-lab.github.io/scRNA.seq.datasets/human/pancreas/ as a log normalized single cell experiment R object and processed using the R package Seurat v3.2.3 (Stuart et al., 2019).

### Analytic Procedures

Insulin measurements from plasma and pancreatic extracts were measured by ELISA (Crystal Chem), blood glucose and luciferase assays were measured as described (Poy *et al*., 2009). Islet morphometric analysis was performed on 8μm sections of paraffin-embedded pancreas approximately 150-200 μm apart. Sections were dewaxed, washed, and stained for insulin (Dako A0564), glucagon (Millipore MABN238), Ki-67 (NovaCastra), or TUNEL (Roche cat. no. 11684795910). Cell numbers from all islets in 3-7 sections were counted with ImageJ software from 20X images obtained using a Nikon A1RSI Spectral Confocal Microscope. In vivo insulin release and glucose (GTT) tolerance tests were performed following a 6-hour fast and intraperitoneally injection of glucose (2g/kg BW). Insulin secretion from isolated islets was performed as described (Poy *et al*., 2009).

### Cell Culture, immunoprecipitation, and western blotting antibodies

MIN6 cells were cultured in DMEM (Invitrogen) containing 4.5g/L glucose supplemented with 15% v/v heat-inactivated FCS, 50 μMβ-mercaptoethanol, and 50 mg/mL penicillin and 100 mg/ml streptomycin and insulin release was performed as described (Poy *et al*., 2004). The following primary antibodies were used for Western blots at 1:1000 dilution: PITPNA (Abcam, ab180234), Cadm1 (MBL, CM004-3), Gephyrin (BD Biosciences, 610585), CHOP (Cell Signaling, 2895S), BiP/GRP78 (Cell Signaling, 3177S), DRP1 (Proteintech, 12957-1-AP), β-Actin (Cell Signaling, 3700S), and γ-Tubulin (Sigma, T6557). The following primary antibodies were used for immunofluorescence: PITPNA (1:200, Sigma, SAB1400211). Antibodies were used on paraffin-embedded pancreata fixed in 4% paraformaldehyde for 3 hours. Image densitometry of 16-bit TIF images for all Western blots was performed using ImageJ. MicroRNA mimics and siRNA pools were purchased from Qiagen GmbH (Germany) and scrambled pool controls are defined as an equimolar stock solution of either 48 random siRNA sequences, or 12 unique mimics of miRNAs not expressed in the beta-cell (i.e. miR-122, miR-1) and not predicted by the TargetScan algorithm to bind the 3’UTR of *Pitpna*. For biochemical fractionation, an eight-step sucrose gradient was performed on MIN6 cells as described previously (Tattikota *et al*., 2013). Briefly, MIN6 cells were washed, pelleted, and resuspended in buffer containing 5 mM HEPES, 0.5 mM EGTA, and 1X Complete Protease inhibitors (Roche Applied Science) at pH 7.4 and homogenized. Homogenate was spun at 3000 X g for 10 min at 4 °C, and the post-nuclear supernatant was loaded onto an 8-step discontinuous sucrose density gradient (HEPES-buffered 0.2–2 M sucrose) and centrifuged at 55,000 rpm for 2h at 4 °C using an MLS50 rotor (Beckman Coulter). Extracellular acidification rate (ECAR) and oxygen consumption rate (OCR) were measured in MIN6 cells using an XF24 Analyzer (Seahorse Bioscience, MA, USA).

### Lentiviral-mediated over-expression and knockdown in isolated human islets

Lentiviruses were generated after subcloning the *PITPNA* cDNA sequence into the expression vector pCCL-cPPT-PGK-IRES-WPRE (Addgene). The resulting construct was transfected along with packaging plasmids pMD2.G and pSPAX2 (Addgene) into HEK293T cells. Cell culture media containing the virus was collected 48 and 72 hours after transfection, concentrated and stored at −80°C. Knockdown of *PITPNA* by MISSION shRNA vectors (Sigma-Aldrich) was confirmed in human pancreatic 1.1B4 cells and isolated islets. Human islets were treated with non-overlapping shRNAs against the human *PITPNA* mRNA (accession number NM_006224), and TRCN00000299703 (SHCLNV 06302009MN) was used for all studies. TRC2 pLKO.5 Lentiviral Transduction Particles (pLKO.5-puro non-Mammalian shRNA Control Plasmid DNA; SHC00204V) were used to treat control human islets. Polybrene (Santa Cruz Biotechnology, Cat# sc-134220, Texas, USA) was added to the media with the final concentration of 10 μg/ml before infection. In brief, 250 islet equivalents (IEQ) seeded in each 12-well plate were infected with each lentivirus at an M.O.I of 20 for 48-72 hours to ensure complete infection.

### Total internal reflection fluorescence (TIRF) microscopy

For TIRF microscopy experiments, human islets were obtained from the Nordic Network for Clinical Islet Transplantation, Uppsala Sweden, with ethical clearance (Uppsala Regional Ethics Board 2006/348) and the donor families’ written informed consent. Islets (donor IDs R442, 2583, 2585) were dispersed into single cells in cell dissociation buffer (Thermo Fisher Scientific) supplemented with trypsin (0.005%, Life Technologies), washed and plated in serum-containing medium on 22-mm polylysine-coated coverslips, allowed to settle overnight, and then transduced with adenovirus coding for the granule marker NPY-tdmOrange2. Cells were imaged as described previously (Gandasi *et al*., 2018) using a lens-type total internal reflection (TIRF) microscope, based on an AxioObserver Z1 with a 100×/1.46 objective (Carl Zeiss). TIRF illumination with a decay constant of ~100 nm (calculated based on exit angle) was created using two DPSS lasers at 491 and 561 nm (Cobolt, Stockholm, Sweden) that passed through a cleanup filter (zet405/488/561/640×, Chroma) and was controlled with an acousto-optical tunable filter (AA-Opto, France). Excitation and emission light were separated using a beamsplitter (ZT405/488/561/640rpc, Chroma) and the emission light chromatically separated (QuadView, Roper) onto separate areas of an EMCCD camera (QuantEM 512SC, Roper) with a cutoff at 565 nm (565dcxr, Chroma) and emission filters (ET525/50m and 600/50m, Chroma). Scaling was 160 nm per pixel. Cells were imaged in (mM) 138 NaCl, 5.6 KCl, 1.2 MgCl2, 2.6 CaCl2, 0.2 diazoxide (to prevent spontaneous depolarizations), 10 D-glucose, 5 HEPES (pH 7.4 with NaOH) at ~35°C. Exocytosis was evoked with high 75 mM K^+^ (equimolarly replacing Na^+^), applied by computer-timed local pressure ejection through a pulled glass capillary. Exocytosis events were identified manually based on the characteristic rapid loss of the granule marker fluorescence (1-2 frames).

### Phosphoinositide determinations and quantitation

Phospholipids were extracted and analyzed by the standard procedure as described (de la Cruz et al., 2020; Traynor-Kaplan et al., 2017). Briefly, adherent human islets were washed with PBS and collected from 6 well plates then transfer into Lo-Bind polypropylene tubes followed by centrifuged at 30,000 g for 1 min at 4°C. After removing the PBS, 0.5 M TCA was added to the pellet, vortexed, and incubated on ice for 10 min. The cooled mixture (TCA and islets) was centrifuged at 30,000 g for 3 min at 4°C and discarded the supernatant. Finally, added 5% (w/v) TCA containing 10 mM EDTA to the pellet and vortexed, and then stored at −80°C. Internal standards of PtdIns(4,5)P_2_, PtdIns(4)P, and PtdIns were added to the precipitates. The lipid analytical internal standards were ammonium salts from Avanti Polar Lipids (LIPID MAPS MS Standards). For lipid extraction, added samples with ice-cold chloroform-methanol-12.1 N HCl (40:80:1). The organic layer was then separated and evaporated. The dried extracts were derivatized (methylated) with TMS-DM and quantified by targeted analysis as described (Traynor-Kaplan *et al*., 2017). Summary of mass spectrometry data presented in Table S3.

### Analysis of intracellular calcium

Intracellular calcium [_i_Ca^+2^] assay was performed (Fluo-4NW Invitrogen, F36206, excitation 494 nm, emission 516 nm) according to the manufacturer’s instructions. Briefly, [_i_Ca^+2^] was recorded for 60-90 s after addition of the KCl. Human islets were fixed in black 96-well optical bottom plates with poly-D-lysine coating. After the dye loading for an hour, the recording was done under confocal microscope (40× objective) at room temp using an excitation filter of 488 nm. Fold change [_i_Ca^+2^] was calculated from the baseline fluorescence recorded during the first 5 s before the addition of KCl. Images were captured at 1 s intervals for up to 60 s and the intracellular free calcium concentration is represented by mean fluorescence intensity.

### Immunostaining and confocal microscopy

The following primary antibodies were used for immunofluorescence: anti-GOLPH3 (1:1000, Abcam, ab98023), anti-Golgin97 (1:100, Invitrogen, A-21270), anti-PtdIns-4-P (1:500, Echelon Biosciences cat. no. Z-P004), and anti-insulin (1:1000, Dako cat.no A0564). For immunostaining, both primary and secondary antibodies were diluted in 1× PBS containing 2.5% bovine serum albumin and 0.2% Triton-X-100. Antibody incubation steps (primary antibody: 3-4 hrs; secondary antibody: 1hr) were performed in a humidified chamber protected from direct light. Alexa Fluor 488, 594, 647 anti-rabbit, anti-mouse or anti-guinea pig secondary antibodies used in this study are listed in Table S2. Cell nuclei were stained with DAPI and mounted with Fluorsave reagent (MilliporeSigma, 345789) for fluorescence microscopy. Confocal images were acquired on a Nikon TiE confocal microscope using the NIS-Elements software with 60× oil immersion objective. Images were imported into the *Fiji* version (http://fiji.sc) of the *ImageJ* software and the colocalization analyses were performed using the Coloc2 plugin (https://imagej.net/plugins/coloc-2) -- an automated system that evaluates the fluorescent intensities of every pixel within an area of interest. Quantification of colocalization was performed using Pearson’s correlation coefficient. The Pearson’s correlation coefficient reflects the degree of linear relationship between two variables; in this case, the fluorescence intensities of two fluorescently tagged proteins GOLPH3 and Golgin97.

### Transmission electron microscopy (TEM)

Isolated mouse islets and MIN6 cells were fixed in 2% paraformaldehyde/2.5% glutaraldehyde in 0.1M Sodium Cacodylate buffer (cat. no. 15960-01 Electron Microscopy Sciences) for 2 hours at 4°C and then stained in 1.0% osmium tetroxide (cat. no. 19100 Electron Microscopy Sciences) for 1 hour. After dehydrated in ethanol, cells were embedded with Spurr’s Low Viscosity Embedding Kit (cat. no. EM0300-1KT, Electron Microscopy Sciences), sectioned (70-90 nm thick), placed on Formvar (200 mesh) copper grids and contrasted with uranyl acetate (cat. no. 22409 Electron Microscopy Sciences) and lead citrate (cat. no. 22410 Electron Microscopy Sciences). Imaging was performed on a Philips Morgagni transmission electron microscope and acquired images were analyzed with respect to insulin granule and mitochondrial morphology.

### Lipid extraction and mass spectrometric analysis

Cells were harvested by trypsinization and washed twice with ice-cold PBS. A modified protocol of Bligh & Dyer was used to extract lipids from cells (Bligh and Dyer, 1959). Briefly, 900μL of chloroform:methanol (1:2, v:v) (Thermo Fisher) was added to 2× 10^6^ cells. After vortexing for 1 minute and incubating for 15 minutes on ice, 300μL of chloroform was added to the mixture, followed by mild vortexing and addition of 300μL distilled water. The mixture was vortexed for 2 minutes and centrifuged at 14,4000 rpm for 2 minutes at 4°C. The lipids were isolated from the lower organic phase. The sample was vacuum dried (Thermo Savant SPD SpeedVac) and the dried extract resuspended in 200μL of chloroform:methanol (1:2, v/v) containing standards: PC 28:0, PE 28:0, PI 25:0, PG 28:0, PA 28:0, PS 28:0, LPC 17:0, LPE 14:0, d_6_-CE 18:0 and d_5_-TAG 48:0 (Avanti Polar Lipids). Phospholipids and neutral lipids were analyzed on an Agilent 1290 HPLC system coupled with an Agilent Triple Quadrupole mass spectrometer 6460, using Zorbax Eclipse Plus C18 column, 2.1×50mm, 1.8μm. The mobile phases were: A (acetonitrile:10mM ammonium formate, 40:60) and B (acetonitrile:10mM ammonium formate, 90:10). For phospholipids separation the gradient was as follows: start at 20% B, to 60% B in 2min, to 100% B in 5min, hold at 100% B for 2 min, back to 20% be in 0.01 min, hold 20% B 1.79 min (total runtime 10.8mins), the flow rate was 0.4 mL/min and the column temperature 30°C. For neutral lipids separation the gradient was as follows: start at 20% B, to 75% B in 2min, to 100% B in 4min, hold at 100% B for 3 min, back to 20% be in 0.01 min, hold 20% B 1.79 min (total runtime 10.8mins), the flow rate was 0.4 mL/min and the column temperature 40°C. Positive and negative electrospray ionization (ESI) was undertaken using the following parameters: gas temperature, 300°C; gas flow, 5 l/min; nebulizer, 45 psi; sheath gas temperature, 250°C; and sheath gas flow, 11 l/min; capillary voltage, 3.5 kV. Phospholipids and neutral lipids were measured using multiple reaction monitoring (MRM), details can be found in Table S3. Each biological replicate was measured twice and the average measurement used for analysis. Identification of peaks were based on retention time (RT) and specific MRM transitions for each lipid. Raw peak areas were integrated using Agilent MassHunter Quantitative Analysis software. Individual lipid species were quantified by comparison with spiked internal standards. The molar fractions of individual lipid species and each lipid class were normalized to total lipids as follows: individual lipid intensities were divided by the relevant internal standard’s intensity and multiplied by the standard’s concentration; the obtained concentration value was divided by the sum of all lipids concentrations to yield molar fractions (mol%).

### Statistical analysis

All results are expressed as mean ± standard error (SEM) and statistical analysis is summarized in Table S2. A P-value of less than or equal to 0.05 was considered statistically significant. \**P*<0.05, \*\**P*<0.01, and \*\*\**P*<0.001. All graphical and statistical analyses were performed using the Prism8 software (Graphpad Software, USA) and Microsoft Excel. Comparisons between data sets with two groups were evaluated using an unpaired Student’s t test. ANOVA analysis was performed for comparisons of three or more groups.

## ACKNOWLEDGEMENTS

The authors thank David Castle, T. Osborne, L. Nagy, and M. Elena-Arango for assistance in the conduct of this work. This work was funded by Johns Hopkins All Children’s Hospital (M.N.P.), NIH grant R35 GM131804 (V.A.B.), the Helmholtz Gemeinschaft (M.N.P.), two European Foundation for the Study of Diabetes EFSD/Lilly Programme Grants (M.N.P. and S.B.), Swedish Science Council (S.B.), NovoNordisk Foundation (S.B.), and the Deutsche Forschungsgemeinschaft (YA 721/3-1 to X.Y.).

## AUTHOR CONTRIBUTIONS

Y.T.Y. and C.S. performed the primary expression analysis, animal husbandry, electron microscopy image analysis, immunohistochemical, and morphometric analysis and edited the manuscript. X.Y., Y.W., J.S., S.K., S.N., and A.A, performed expression analysis. L.L. and S.B. performed TIRF microscopy and edited the manuscript. A.P. and M.M. performed immunohistochemical analysis. A.G. and F.v.M. reanalyzed public expression datasets. Y.W., A.C-G. and M.W. performed and analyzed the lipidomic analysis. A.T.K. quantified phosphoinositides. P.A. edited the manuscript. Z.X and V.A.B. developed and provided the *Pitpna* mutant animal lines and edited the manuscript. M.N.P. conceived and designed the study, wrote the manuscript, and is the guarantor of this work and takes responsibility for the integrity of the data and the accuracy of the data analysis. All authors contributed to interpretation of the data and approved the final version of this manuscript.

## DECLARATIONS OF INTERSTS

The authors declare no competing interests.

---

**Figure S1. *Pitpna* is a direct target of miR-375 in the pancreatic beta-cell**. **A**, Reporter activity in MIN6 cells transfected with a Renilla luciferase reporter construct containing the 3’UTR of the *Pitpna* gene in addition to either a miR-375-mimic or scrambled control mimic pool. *Pitpna* WT, construct contains the wild-type sequence of *Pitpna* 3’UTR; *Pitpna* MUT, construct contains a 3’UTR sequence where 4 nucleotides of the putative sequence complementary to the miR-375 seed sequence of *Pitpna* 3 ‘UTR were mutated (n=4). **B**, qRT-PCR analysis for miR-375 and *Pitpna* expression (n=4) in MIN6 cells after transfection of either an inhibitory antisense RNA oligonucleotide complimentary to miR-375 (Antg-375) or scrambled RNA oligonucleotide control pool (Antg-ctrl). **C**, Western blot analysis of Pitpna, Cadm1 and Gphn in MIN6 cells, transfected with the Antg-375 or Antg-ctrl. **D**, Western blot analysis of Pitpna, Cadm1, and Gphn in MIN6 cells transfected with the miR-375-mimic or scrambled mimic control pool. **E**, qRT-PCR analysis of *Pitpna, Cadm1, Gephyrin*, and *Ago2* mRNA expression in islets of WT and *Ins*-Cre, *Ago2^flox/flox^* mice at 10 weeks of age (n=4). **F**, Targeted analysis of PtdIns (PI) species in MIN6 cells after transfection of either an inhibitory antisense RNA oligonucleotide complementary to miR-375 (antg-375) or scrambled RNA oligonucleotide control pool (antg-ctrl); Lipid species were expressed as mean molar fractions (n=6). Results are presented as mean ± SEM. \**P*< 0.05, \*\**P*< 0.01, \*\*\**P*<0.001, and n.s. denotes not significant.

**Figure S2. Whole-body *Pitpna* knockout mice exhibit decreased pancreatic beta-cell mass. A**, Immunostaining of insulin and glucagon (Gcg) in paraffin-embedded pancreata from whole-body *Pitpna* knockout (*Pitpna* KO) and littermate wild-type (WT) control mice at age P0. Scale bar = 50 μm. In far-right panel, scale bar = 20 μm. **B**, Quantification of insulin^+^ cells and islet number per area pancreas (mm^2^) in *Pitpna* KO and littermate control mice at age P0 (n=5). **C**, Quantification of total pancreatic insulin and proinsulin content per pancreatic weight (mg) in *Pitpna* KO and littermate control mice at age P0 (n=7-13). **D**, Transmission electron micrographs of pancreas from WT and total *Pitpna* knockout mice (*Pitpna* KO) at age P0. Scale bar= 2μm (left and center panel) and 500 nm (right panel). Dashed black box identifies image in center panel. Yellow arrows identify immature insulin granules (center panel). Solid red and blue lines in center panel identifies plasma membrane. Blue arrows in right panel identify docked vesicles. **E**, Quantification of docked vesicles, and immature secretory granule (ISG), mature secretory granules (MSG), and empty secretory granules (ESG) in beta-cells of WT and *Pitpna* KO mice (n=5-15). **F**, Quantification of granule size in WT and *Pitpna* KO mice (n=4). **G**, Immunostaining of insulin, glucagon (Gcg) in addition to apoptotic marker TUNEL in paraffin-embedded pancreata from total *Pitpna* knockout (*Pitpna* KO) and littermate control (WT) mice at age P0. Scale bar= 30μm. **H**, Quantification of TUNEL-positive beta cells in pancreata from total *Pitpna* knockout (*Pitpna* KO) and littermate control (WT) mice at age P0 (n=5). **I**, Quantification of TUNEL-positive alpha cells in pancreata from total *Pitpna* knockout (*Pitpna* KO) and littermate control (WT) mice at age P0 (n=5). **J**, Immunostaining of insulin and Ki67 in paraffin-embedded pancreata from total *Pitpna* knockout (*Pitpna* KO) and littermate control (WT) mice at age P0. Scale bar = 50 μm. In far-right panel, scale bar =10 μm. **K**, Quantification of Ki67-positive beta cells in pancreata from total *Pitpna* knockout (*Pitpna* KO) and littermate control (WT) mice at age P0 (n=5). Results presented as mean ± SEM. \**P*<0.05, \*\**P*<0.01 and \*\*\**P*<0.001, and n.s. denotes not significant.

**Figure S3 Pitpna regulates pancreatic beta-cell function. A**, Quantification of glucose-stimulated insulin secretion from MIN6 cells after siRNA-mediated knockdown of *Pitpna* or control transected cells (n=4). **B**, **C**, quantitative RT-PCR (n=4) and western blot analysis after siRNA-mediated knockdown of *Pitpna* and scrambled control in MIN6 cells. **D**, Quantification of cellular insulin content after siRNA-mediated knockdown of *Pitpna* and scrambled control in MIN6 cells (n=4). **E**, Measurement of glucose-stimulated insulin release from isolated mouse islets after overexpression of *Pitpna* (n=3). **F**, **G**, quantitative RT-PCR (n=4) and western blot analysis after overexpression of *Pitpna* or scrambled control transfection in MIN6 cells (n=3). **H**, Measurement of cellular insulin content after overexpression of *Pitpna* or scrambled control transfection in MIN6 cells (n=3). Results resented as mean ± S.E.M. \**P*< 0.05; \*\**P*< 0.01; \*\*\**P*< 0.001, and n.s. denotes not significant. AU is arbitrary units.

**Figure S4. Conditional deletion of *Pitpna* in the beta-cell induces alterations in mitochondrial configuration and morphology. A**, Representative Seahorse flux analysis of oxygen consumption rate (OCR) in MIN6 cells after siRNA-mediated knockdown of *Pitpna* (n=7). During experiment, cells were exposed to oligomycin (O), FCCP (F), and the combination of rotenone and antimycin A (R/A) at the time points indicated. **B**, Basal and maximal respiration were measured after either siRNA-mediated knockdown of *Pitpna* or scrambled control transfection in MIN6 cells. **C**, Representative Seahorse flux analysis of extracellular acidification rate (ECAR) in MIN6 cells after siRNA-mediated knockdown of *Pitpna* (n=7). and **D**, Glycolysis and glycolytic capacity were measured after either siRNA-mediated knockdown of *Pitpna* or scrambled control transfection in MIN6 cells. **E**, Representative transmission electron micrographs of mitochondria within pancreatic beta-cells of *Ins*-Cre, *Pitpna^flox/flox^* and littermate control (WT) mice at age 8 weeks (n=4-5). Scale bar = 1μm. **F,** Quantification of mitochondrial length distribution in *Ins*-Cre, *Pitpna^flox/flox^* and littermate control (WT) mice at age 8 weeks (n=4-5). **G**, Representative transmission electron micrographs identify unique mitochondrial configurations. Scale bar = 1μm. **H**, Quantification of distribution of mitochondrial configurations in pancreatic beta-cells of *Ins*-Cre, *Pitpna^flox/flox^* and littermate control (WT) mice at age 8 weeks (n=7). **I**, Western blot analysis of Pitpna, and Dynamin related protein 1 (Drp1) in isolated islets of 10-week-old *Ins*-Cre, *Pitpna^flox/flox^* and littermate control (WT) mice. Results presented as mean ± SEM. \**P*< 0.05, \*\*\**P*< 0.001, and n.s. denotes not significant.

**Figure S5. *PITPNA* regulates mitochondrial morphology in human pancreatic beta-cells. A**, Quantification of knockdown of *PITPNA* in human pancreatic 1.1B4 cells by individual shRNA clones by qRT-PCR. **B**, Representative transmission electron micrographs reveal mitochondrial morphology in pancreatic beta-cells from isolated human islets after lentiviral-mediated over-expression of *PITPNA* (OE-*PITPNA*) or inhibition of *PITPNA* (sh-*PITPNA*) or treatment with control lentivirus (sh-*Ctrl*). **C**, Quantification of mitochondrial length distribution in pancreatic beta-cells from isolated human islets after lentiviral-mediated over-expression of *PITPNA* (OE*-PITPNA*) or inhibition of *PITPNA* (sh-*PITPNA*) or treatment with control lentivirus (sh-*Ctrl*). **D**, Mitochondrial morphology in beta-cells of isolated human islets after lentiviral-mediated over-expression of *PITPNA* (OE-*PITPNA*), inhibition of *PITPNA* (sh-*PITPNA*), or treatment with control lentivirus (sh-*Ctrl*) (n=7). **E**, Quantification of phosphatidylinositol (PI) after lentiviral-mediated over-expression of *PITPNA* (OE-*PITPNA*) or treatment with control lentivirus (*Ctrl*) (n=3). **F**, Quantification of phosphatidylinositol (PI) in isolated human islets after lentiviral-mediated inhibition of *PITPNA* (sh-*PITPNA*) or treatment with a lentivirus expressing an shRNA control (*sh-Ctrl*) (n=4). Results are presented as mean ± SEM. \**P*< 0.05; \*\**P*< 0.01; \*\*\**p*< 0.001, and n.s. denotes not significant.

**Figure S6. Restoration of *PITPNA* in isolated islets of T2D human subjects improves insulin secretion and reverses expression of ER stress proteins. A**, Quantification of insulin release from isolated human islets from individual T2D donors after lentiviral-mediated over-expression of *PITPNA (T2D-PITPNA* OE) or treatment with a control lentivirus (T2D-Ctrl) (n=4). **B,** Western blot analysis of PITPNA and ER stress/unfolded protein response (UPR) proteins IRE1α, CHOP, ERO1, PDI, and BiP/Grp78 in T2D human islets after lentiviral-mediated over-expression of *PITPNA* (T2D-OE), or treatment with a control lentivirus (T2D-Ctrl) (n=4). Summary of mean fold change values displayed at right. **C**, Western blot analysis of proinsulin in T2D human islets after lentiviral-mediated over-expression of *PITPNA* (T2D-OE), or treatment with a control lentivirus (T2D-Ctrl) (n=4). Summary of mean fold change values displayed at right. Results presented as mean ± SEM. \**P*< 0.05; \*\**P*< 0.01, and n.s. denotes not significant.

## Notes

### Competing Interest Statement

The authors have declared no competing interest.

## REFERENCES

Agarwal, V., Subtelny, A.O., Thiru, P., Ulitsky, I., and Bartel, D.P. (2018). Predicting microRNA targeting efficacy in Drosophila. Genome Biol 19, 152. 10.1186/s13059-018-1504-3.

Alb, J.G., Jr., Cortese, J.D., Phillips, S.E., Albin, R.L., Nagy, T.R., Hamilton, B.A., and Bankaitis, V.A. (2003). Mice lacking phosphatidylinositol transfer protein-alpha exhibit spinocerebellar degeneration, intestinal and hepatic steatosis, and hypoglycemia. J Biol Chem 278, 33501–33518. 10.1074/jbc.M303591200.

Alb, J.G., Jr., Gedvilaite, A., Cartee, R.T., Skinner, H.B., and Bankaitis, V.A. (1995). Mutant rat phosphatidylinositol/phosphatidylcholine transfer proteins specifically defective in phosphatidylinositol transfer: implications for the regulation of phospholipid transfer activity. Proc Natl Acad Sci U S A 92, 8826–8830. 10.1073/pnas.92.19.8826.

Alb, J.G., Jr., Phillips, S.E., Wilfley, L.R., Philpot, B.D., and Bankaitis, V.A. (2007). The pathologies associated with functional titration of phosphatidylinositol transfer protein alpha activity in mice. J Lipid Res 48, 1857–1872. 10.1194/jlr.M700145-JLR200.

Alejandro, E.U., Gregg, B., Blandino-Rosano, M., Cras-Meneur, C., and Bernal-Mizrachi, E. (2015). Natural history of beta-cell adaptation and failure in type 2 diabetes. Mol Aspects Med 42, 19–41. 10.1016/j.mam.2014.12.002.

Arunagiri, A., Haataja, L., Cunningham, C.N., Shrestha, N., Tsai, B., Qi, L., Liu, M., and Arvan, P. (2018). Misfolded proinsulin in the endoplasmic reticulum during development of beta cell failure in diabetes. Ann N Y Acad Sci 1418, 5–19. 10.1111/nyas.13531.

Arunagiri, A., Haataja, L., Pottekat, A., Pamenan, F., Kim, S., Zeltser, L.M., Paton, A.W., Paton, J.C., Tsai, B., Itkin-Ansari, P., et al. (2019). Proinsulin misfolding is an early event in the progression to type 2 diabetes. Elife 8. 10.7554/eLife.44532.

Ashlin, T.G., Blunsom, N.J., and Cockcroft, S. (2021). Courier service for phosphatidylinositol: PITPs deliver on demand. Biochim Biophys Acta Mol Cell Biol Lipids 1866, 158985. 10.1016/j.bbalip.2021.158985.

Back, S.H., and Kaufman, R.J. (2012). Endoplasmic reticulum stress and type 2 diabetes. Annu Rev Biochem 81, 767–793. 10.1146/annurev-biochem-072909-095555.

Balla, T. (2013). Phosphoinositides: tiny lipids with giant impact on cell regulation. Physiol Rev 93, 1019–1137. 10.1152/physrev.00028.2012.

Bankaitis, V.A., Aitken, J.R., Cleves, A.E., and Dowhan, W. (1990). An essential role for a phospholipid transfer protein in yeast Golgi function. Nature 347, 561–562. 10.1038/347561a0.

Bankaitis, V.A., Garcia-Mata, R., and Mousley, C.J. (2012). Golgi membrane dynamics and lipid metabolism. Curr Biol 22, R414–424. 10.1016/j.cub.2012.03.004.

Bankaitis, V.A., Mousley, C.J., and Schaaf, G. (2010). The Sec14 superfamily and mechanisms for crosstalk between lipid metabolism and lipid signaling. Trends Biochem Sci 35, 150–160. 10.1016/j.tibs.2009.10.008.

Behnia, R., and Munro, S. (2005). Organelle identity and the signposts for membrane traffic. Nature 438, 597–604. 10.1038/nature04397.

Bligh, E.G., and Dyer, W.J. (1959). A rapid method of total lipid extraction and purification. Can J Biochem Physiol 37, 911–917. 10.1139/o59-099.

Bock, F.J., and Tait, S.W.G. (2020). Mitochondria as multifaceted regulators of cell death. Nat Rev Mol Cell Biol 21, 85–100. 10.1038/s41580-019-0173-8.

Bridges, D., and Saltiel, A.R. (2012). Phosphoinositides in insulin action and diabetes. Curr Top Microbiol Immunol 362, 61–85. 10.1007/978-94-007-5025-8_3.

Campbell, J.E., and Newgard, C.B. (2021). Mechanisms controlling pancreatic islet cell function in insulin secretion. Nat Rev Mol Cell Biol 22, 142–158. 10.1038/s41580-020-00317-7.

Chen, C., Cohrs, C.M., Stertmann, J., Bozsak, R., and Speier, S. (2017). Human beta cell mass and function in diabetes: Recent advances in knowledge and technologies to understand disease pathogenesis. Mol Metab 6, 943–957. 10.1016/j.molmet.2017.06.019.

Cleves, A.E., McGee, T.P., Whitters, E.A., Champion, K.M., Aitken, J.R., Dowhan, W., Goebl, M., and Bankaitis, V.A. (1991). Mutations in the CDP-choline pathway for phospholipid biosynthesis bypass the requirement for an essential phospholipid transfer protein. Cell 64, 789–800. 10.1016/0092-8674(91)90508-v.

Cruz-Garcia, D., Ortega-Bellido, M., Scarpa, M., Villeneuve, J., Jovic, M., Porzner, M., Balla, T., Seufferlein, T., and Malhotra, V. (2013). Recruitment of arfaptins to the trans-Golgi network by PI(4)P and their involvement in cargo export. EMBO J 32, 1717–1729. 10.1038/emboj.2013.116.

De Camilli, P., Emr, S.D., McPherson, P.S., and Novick, P. (1996). Phosphoinositides as regulators in membrane traffic. Science 271, 1533–1539. 10.1126/science.271.5255.1533.

de la Cruz, L., Traynor-Kaplan, A., Vivas, O., Hille, B., and Jensen, J.B. (2020). Plasma membrane processes are differentially regulated by type I phosphatidylinositol phosphate 5-kinases and RASSF4. J Cell Sci 133. 10.1242/jcs.233254.

Di Paolo, G., and De Camilli, P. (2006). Phosphoinositides in cell regulation and membrane dynamics. Nature 443, 651–657. 10.1038/nature05185.

Dickeson, S.K., Lim, C.N., Schuyler, G.T., Dalton, T.P., Helmkamp, G.M., Jr., and Yarbrough, L.R. (1989). Isolation and sequence of cDNA clones encoding rat phosphatidylinositol transfer protein. J Biol Chem 264, 16557–16564.

Eizirik, D.L., Pasquali, L., and Cnop, M. (2020). Pancreatic beta-cells in type 1 and type 2 diabetes mellitus: different pathways to failure. Nat Rev Endocrinol 16, 349–362. 10.1038/s41574-020-0355-7.

Fadista, J., Vikman, P., Laakso, E.O., Mollet, I.G., Esguerra, J.L., Taneera, J., Storm, P., Osmark, P., Ladenvall, C., Prasad, R.B., et al. (2014). Global genomic and transcriptomic analysis of human pancreatic islets reveals novel genes influencing glucose metabolism. Proc Natl Acad Sci U S A 111, 13924–13929. 10.1073/pnas.1402665111.

Ferrannini, E. (2010). The stunned beta cell: a brief history. Cell Metab 11, 349–352. 10.1016/j.cmet.2010.04.009.

Fonseca, S.G., Gromada, J., and Urano, F. (2011). Endoplasmic reticulum stress and pancreatic beta-cell death. Trends Endocrinol Metab 22, 266–274. 10.1016/j.tem.2011.02.008.

Friedman, J.R., Lackner, L.L., West, M., DiBenedetto, J.R., Nunnari, J., and Voeltz, G.K. (2011). ER tubules mark sites of mitochondrial division. Science 334, 358–362. 10.1126/science.1207385.

Fullwood, Y., dos Santos, M., and Hsuan, J.J. (1999). Cloning and characterization of a novel human phosphatidylinositol transfer protein, rdgBbeta. J Biol Chem 274, 31553–31558. 10.1074/jbc.274.44.31553.

Gandasi, N.R., Yin, P., Omar-Hmeadi, M., Ottosson Laakso, E., Vikman, P., and Barg, S. (2018). Glucose-Dependent Granule Docking Limits Insulin Secretion and Is Decreased in Human Type 2 Diabetes. Cell Metab 27, 470–478 e474. 10.1016/j.cmet.2017.12.017.

Grabon, A., Bankaitis, V.A., and McDermott, M.I. (2019). The interface between phosphatidylinositol transfer protein function and phosphoinositide signaling in higher eukaryotes. J Lipid Res 60, 242–268. 10.1194/jlr.R089730.

Hammond, G.R., Fischer, M.J., Anderson, K.E., Holdich, J., Koteci, A., Balla, T., and Irvine, R.F. (2012). PI4P and PI(4,5)P2 are essential but independent lipid determinants of membrane identity. Science 337, 727–730. 10.1126/science.1222483.

Harding, H.P., and Ron, D. (2002). Endoplasmic reticulum stress and the development of diabetes: a review. Diabetes 51 Suppl 3, S455–461. 10.2337/diabetes.51.2007.s455.

Hay, J.C., and Martin, T.F. (1993). Phosphatidylinositol transfer protein required for ATP-dependent priming of Ca(2+)-activated secretion. Nature 366, 572–575. 10.1038/366572a0.

Haythorne, E., Rohm, M., van de Bunt, M., Brereton, M.F., Tarasov, A.I., Blacker, T.S., Sachse, G., Silva Dos Santos, M., Terron Exposito, R., Davis, S., et al. (2019). Diabetes causes marked inhibition of mitochondrial metabolism in pancreatic beta-cells. Nat Commun 10, 2474. 10.1038/s41467-019-10189-x.

Helmkamp, G.M., Jr., Harvey, M.S., Wirtz, K.W., and Van Deenen, L.L. (1974). Phospholipid exchange between membranes. Purification of bovine brain proteins that preferentially catalyze the transfer of phosphatidylinositol. J Biol Chem 249, 6382–6389.

Hennings, T.G., Chopra, D.G., DeLeon, E.R., VanDeusen, H.R., Sesaki, H., Merrins, M.J., and Ku, G.M. (2018). In Vivo Deletion of beta-Cell Drp1 Impairs Insulin Secretion Without Affecting Islet Oxygen Consumption. Endocrinology 159, 3245–3256. 10.1210/en.2018-00445.

Hokin, M.R., and Hokin, L.E. (1953). Enzyme secretion and the incorporation of P32 into phospholipides of pancreas slices. J Biol Chem 203, 967–977.

Kim, D., Langmead, B., and Salzberg, S.L. (2015). HISAT: a fast spliced aligner with low memory requirements. Nat Methods 12, 357–360. 10.1038/nmeth.3317.

Kuna, R.S., and Field, S.J. (2019). GOLPH3: a Golgi phosphatidylinositol(4)phosphate effector that directs vesicle trafficking and drives cancer. J Lipid Res 60, 269–275. 10.1194/jlr.R088328.

LaPierre, M.P., and Stoffel, M. (2017). MicroRNAs as stress regulators in pancreatic beta cells and diabetes. Mol Metab 6, 1010–1023. 10.1016/j.molmet.2017.06.020.

Lete, M.G., Tripathi, A., Chandran, V., Bankaitis, V.A., and McDermott, M.I. (2020). Lipid transfer proteins and instructive regulation of lipid kinase activities: Implications for inositol lipid signaling and disease. Adv Biol Regul 78, 100740. 10.1016/j.jbior.2020.100740.

Liesa, M., and Shirihai, O.S. (2013). Mitochondrial dynamics in the regulation of nutrient utilization and energy expenditure. Cell Metab 17, 491–506. 10.1016/j.cmet.2013.03.002.

Liu, M., Huang, Y., Xu, X., Li, X., Alam, M., Arunagiri, A., Haataja, L., Ding, L., Wang, S., Itkin-Ansari, P., et al. (2021). Normal and defective pathways in biogenesis and maintenance of the insulin storage pool. J Clin Invest 131. 10.1172/JCI142240.

MacDonald, M.J., Ade, L., Ntambi, J.M., Ansari, I.U., and Stoker, S.W. (2015). Characterization of phospholipids in insulin secretory granules and mitochondria in pancreatic beta cells and their changes with glucose stimulation. J Biol Chem 290, 11075–11092. 10.1074/jbc.M114.628420.

Marciniak, S.J., Yun, C.Y., Oyadomari, S., Novoa, I., Zhang, Y., Jungreis, R., Nagata, K., Harding, H.P., and Ron, D. (2004). CHOP induces death by promoting protein synthesis and oxidation in the stressed endoplasmic reticulum. Genes Dev 18, 3066–3077. 10.1101/gad.1250704.

Martin, T.F. (1998). Phosphoinositide lipids as signaling molecules: common themes for signal transduction, cytoskeletal regulation, and membrane trafficking. Annu Rev Cell Dev Biol 14, 231–264. 10.1146/annurev.cellbio.14.1.231.

Mishra, P., and Chan, D.C. (2014). Mitochondrial dynamics and inheritance during cell division, development and disease. Nat Rev Mol Cell Biol 15, 634–646. 10.1038/nrm3877.

Mousley, C.J., Tyeryar, K., Ile, K.E., Schaaf, G., Brost, R.L., Boone, C., Guan, X., Wenk, M.R., and Bankaitis, V.A. (2008). Trans-Golgi network and endosome dynamics connect ceramide homeostasis with regulation of the unfolded protein response and TOR signaling in yeast. Mol Biol Cell 19, 4785–4803. 10.1091/mbc.E08-04-0426.

Muoio, D.M., and Newgard, C.B. (2008). Mechanisms of disease:Molecular and metabolic mechanisms of insulin resistance and beta-cell failure in type 2 diabetes. Nat Rev Mol Cell Biol 9, 193–205. 10.1038/nrm2327.

Muraro, M.J., Dharmadhikari, G., Grun, D., Groen, N., Dielen, T., Jansen, E., van Gurp, L., Engelse, M.A., Carlotti, F., de Koning, E.J., and van Oudenaarden, A. (2016). A Single-Cell Transcriptome Atlas of the Human Pancreas. Cell Syst 3, 385–394 e383. 10.1016/j.cels.2016.09.002.

Nagashima, S., Tabara, L.C., Tilokani, L., Paupe, V., Anand, H., Pogson, J.H., Zunino, R., McBride, H.M., and Prudent, J. (2020). Golgi-derived PI(4)P-containing vesicles drive late steps of mitochondrial division. Science 367, 1366–1371. 10.1126/science.aax6089.

Nguyen, P.M., Gandasi, N.R., Xie, B., Sugahara, S., Xu, Y., and Idevall-Hagren, O. (2019). The PI(4)P phosphatase Sac2 controls insulin granule docking and release. J Cell Biol 218, 3714–3729. 10.1083/jcb.201903121.

Nolan, C.J., and Prentki, M. (2019). Insulin resistance and insulin hypersecretion in the metabolic syndrome and type 2 diabetes: Time for a conceptual framework shift. Diab Vasc Dis Res 16, 118–127. 10.1177/1479164119827611.

Ohashi, M., Jan de Vries, K., Frank, R., Snoek, G., Bankaitis, V., Wirtz, K., and Huttner, W.B. (1995). A role for phosphatidylinositol transfer protein in secretory vesicle formation. Nature 377, 544–547. 10.1038/377544a0.

Omar-Hmeadi, M., and Idevall-Hagren, O. (2021). Insulin granule biogenesis and exocytosis. Cell Mol Life Sci 78, 1957–1970. 10.1007/s00018-020-03688-4.

Porksen, N., Hollingdal, M., Juhl, C., Butler, P., Veldhuis, J.D., and Schmitz, O. (2002). Pulsatile insulin secretion: detection, regulation, and role in diabetes. Diabetes 51 Suppl 1, S245–254. 10.2337/diabetes.51.2007.s245.

Poy, M.N. (2016). MicroRNAs: An adaptive mechanism in the pancreatic beta-cell…and beyond? Best Pract Res Clin Endocrinol Metab 30, 621–628. 10.1016/j.beem.2016.07.003.

Poy, M.N., Eliasson, L., Krutzfeldt, J., Kuwajima, S., Ma, X., Macdonald, P.E., Pfeffer, S., Tuschl, T., Rajewsky, N., Rorsman, P., and Stoffel, M. (2004). A pancreatic islet-specific microRNA regulates insulin secretion. Nature 432, 226–230. 10.1038/nature03076.

Poy, M.N., Hausser, J., Trajkovski, M., Braun, M., Collins, S., Rorsman, P., Zavolan, M., and Stoffel, M. (2009). miR-375 maintains normal pancreatic alpha-and beta-cell mass. Proc Natl Acad Sci U S A 106, 5813–5818. 10.1073/pnas.0810550106.

Rameh, L.E., and Deeney, J.T. (2016). Phosphoinositide signalling in type 2 diabetes: a beta-cell perspective. Biochem Soc Trans 44, 293–298. 10.1042/BST20150229.

Rhodes, C.J. (2005). Type 2 diabetes-a matter of beta-cell life and death? Science 307, 380–384. 10.1126/science.1104345.

Rohm, T.V., Meier, D.T., Olefsky, J.M., and Donath, M.Y. (2022). Inflammation in obesity, diabetes, and related disorders. Immunity 55, 31–55. 10.1016/j.immuni.2021.12.013.

Schaaf, G., Ortlund, E.A., Tyeryar, K.R., Mousley, C.J., Ile, K.E., Garrett, T.A., Ren, J., Woolls, M.J., Raetz, C.R., Redinbo, M.R., and Bankaitis, V.A. (2008). Functional anatomy of phospholipid binding and regulation of phosphoinositide homeostasis by proteins of the sec14 superfamily. Mol Cell 29, 191–206. 10.1016/j.molcel.2007.11.026.

Shrestha, N., De Franco, E., Arvan, P., and Cnop, M. (2021). Pathological beta-Cell Endoplasmic Reticulum Stress in Type 2 Diabetes: Current Evidence. Front Endocrinol (Lausanne) 12, 650158. 10.3389/fendo.2021.650158.

Sidarala, V., Pearson, G.L., Parekh, V.S., Thompson, B., Christen, L., Gingerich, M.A., Zhu, J., Stromer, T., Ren, J., Reck, E.C., et al. (2020). Mitophagy protects beta cells from inflammatory damage in diabetes. JCI Insight 5. 10.1172/jci.insight.141138.

Smirnova, E., Griparic, L., Shurland, D.L., and van der Bliek, A.M. (2001). Dynamin-related protein Drp1 is required for mitochondrial division in mammalian cells. Mol Biol Cell 12, 2245–2256. 10.1091/mbc.12.8.2245.

Song, B., Scheuner, D., Ron, D., Pennathur, S., and Kaufman, R.J. (2008). Chop deletion reduces oxidative stress, improves beta cell function, and promotes cell survival in multiple mouse models of diabetes. J Clin Invest 118, 3378–3389. 10.1172/JCI34587.

Stephens, L.R., Jackson, T.R., and Hawkins, P.T. (1993). Agonist-stimulated synthesis of phosphatidylinositol(3,4,5)-trisphosphate: a new intracellular signalling system? Biochim Biophys Acta 1179, 27–75. 10.1016/0167-4889(93)90072-w.

Stuart, T., Butler, A., Hoffman, P., Hafemeister, C., Papalexi, E., Mauck, W.M., 3rd, Hao, Y., Stoeckius, M., Smibert, P., and Satija, R. (2019). Comprehensive Integration of Single-Cell Data. Cell 177, 1888–1902 e1821. 10.1016/j.cell.2019.05.031.

Sun, J., Cui, J., He, Q., Chen, Z., Arvan, P., and Liu, M. (2015). Proinsulin misfolding and endoplasmic reticulum stress during the development and progression of diabetes. Mol Aspects Med 42, 105–118. 10.1016/j.mam.2015.01.001.

Tabara, L.C., Morris, J.L., and Prudent, J. (2021). The Complex Dance of Organelles during Mitochondrial Division. Trends Cell Biol 31, 241–253. 10.1016/j.tcb.2020.12.005.

Talchai, C., Lin, H.V., Kitamura, T., and Accili, D. (2009). Genetic and biochemical pathways of beta-cell failure in type 2 diabetes. Diabetes Obes Metab 11 Suppl 4, 38–45. 10.1111/j.1463-1326.2009.01115.x.

Tanaka, S., and Hosaka, K. (1994). Cloning of a cDNA encoding a second phosphatidylinositol transfer protein of rat brain by complementation of the yeast sec14 mutation. J Biochem 115, 981–984. 10.1093/oxfordjournals.jbchem.a124448.

Tattikota, S.G., Rathjen, T., McAnulty, S.J., Wessels, H.H., Akerman, I., van de Bunt, M., Hausser, J., Esguerra, J.L., Musahl, A., Pandey, A.K., et al. (2014). Argonaute2 mediates compensatory expansion of the pancreatic beta cell. Cell Metab 19, 122–134. 10.1016/j.cmet.2013.11.015.

Tattikota, S.G., Sury, M.D., Rathjen, T., Wessels, H.H., Pandey, A.K., You, X., Becker, C., Chen, W., Selbach, M., and Poy, M.N. (2013). Argonaute2 regulates the pancreatic beta-cell secretome. Mol Cell Proteomics 12, 1214–1225. 10.1074/mcp.M112.024786.

Thorens, B., Tarussio, D., Maestro, M.A., Rovira, M., Heikkila, E., and Ferrer, J. (2015). Ins1(Cre) knock-in mice for beta cell-specific gene recombination. Diabetologia 58, 558–565. 10.1007/s00125-014-3468-5.

Traynor-Kaplan, A., Kruse, M., Dickson, E.J., Dai, G., Vivas, O., Yu, H., Whittington, D., and Hille, B. (2017). Fatty-acyl chain profiles of cellular phosphoinositides. Biochim Biophys Acta Mol Cell Biol Lipids 1862, 513–522. 10.1016/j.bbalip.2017.02.002.

van Raalte, D.H., and Verchere, C.B. (2017). Improving glycaemic control in type 2 diabetes: Stimulate insulin secretion or provide beta-cell rest? Diabetes Obes Metab 19, 1205–1213. 10.1111/dom.12935.

Wirtz, K.W. (1991). Phospholipid transfer proteins. Annu Rev Biochem 60, 73–99. 10.1146/annurev.bi.60.070191.000445.

Wright, J., Birk, J., Haataja, L., Liu, M., Ramming, T., Weiss, M.A., Appenzeller-Herzog, C., and Arvan, P. (2013). Endoplasmic reticulum oxidoreductin-1alpha (Ero1alpha) improves folding and secretion of mutant proinsulin and limits mutant proinsulin-induced endoplasmic reticulum stress. J Biol Chem 288, 31010–31018. 10.1074/jbc.M113.510065.

Wuttke, A. (2015). Lipid signalling dynamics at the beta-cell plasma membrane. Basic Clin Pharmacol Toxicol 116, 281–290. 10.1111/bcpt.12369.

Xie, Z., and Bankaitis, V.A. (2022). Phosphatidylinositol transfer protein/planar cell polarity axis regulates neocortical morphogenesis by supporting interkinetic nuclear migration. Cell Rep 39, 110869. 10.1016/j.celrep.2022.110869.

Xie, Z., Hur, S.K., Zhao, L., Abrams, C.S., and Bankaitis, V.A. (2018). A Golgi Lipid Signaling Pathway Controls Apical Golgi Distribution and Cell Polarity during Neurogenesis. Dev Cell 44, 725–740 e724. 10.1016/j.devcel.2018.02.025.

Xin, Y., Kim, J., Okamoto, H., Ni, M., Wei, Y., Adler, C., Murphy, A.J., Yancopoulos, G.D., Lin, C., and Gromada, J. (2016). RNA Sequencing of Single Human Islet Cells Reveals Type 2 Diabetes Genes. Cell Metab 24, 608–615. 10.1016/j.cmet.2016.08.018.

Yong, J., Johnson, J.D., Arvan, P., Han, J., and Kaufman, R.J. (2021). Therapeutic opportunities for pancreatic beta-cell ER stress in diabetes mellitus. Nat Rev Endocrinol 17, 455–467. 10.1038/s41574-021-00510-4.

Youle, R.J., and van der Bliek, A.M. (2012). Mitochondrial fission, fusion, and stress. Science 337, 1062–1065. 10.1126/science.1219855.

Zinszner, H., Kuroda, M., Wang, X., Batchvarova, N., Lightfoot, R.T., Remotti, H., Stevens, J.L., and Ron, D. (1998). CHOP is implicated in programmed cell death in response to impaired function of the endoplasmic reticulum. Genes Dev 12, 982–995. 10.1101/gad.12.7.982.

