## Supplementary figures and images for "Restoration of PITPNA in Type 2 diabetic human islets reverses pancreatic beta-cell dysfunction"

### FIGURE S1

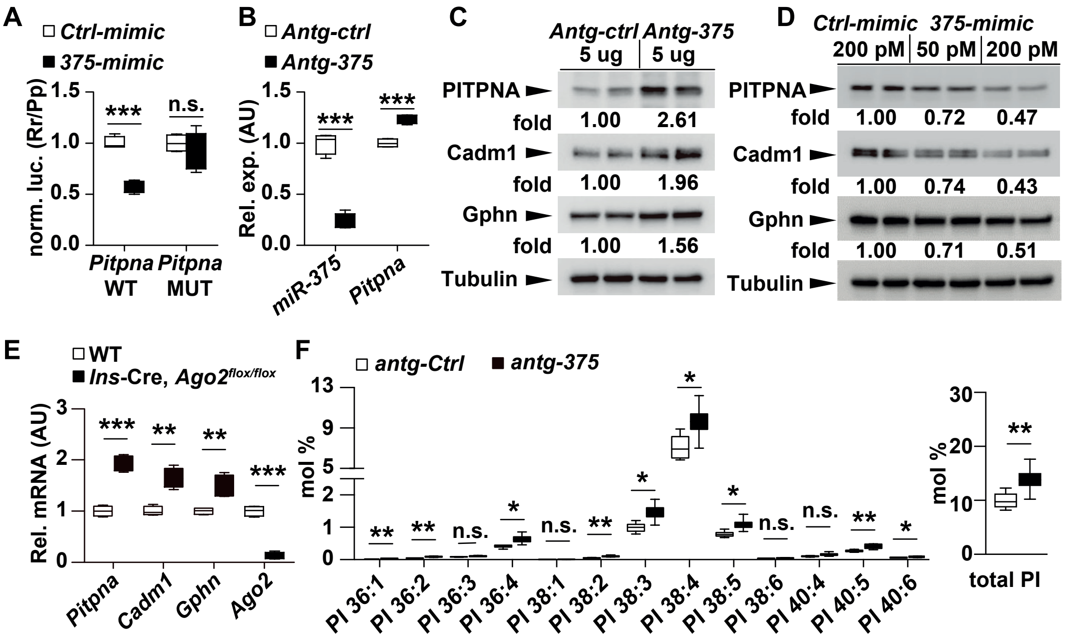


Yeh et al., 2022
Figure S1

### FIGURE S2

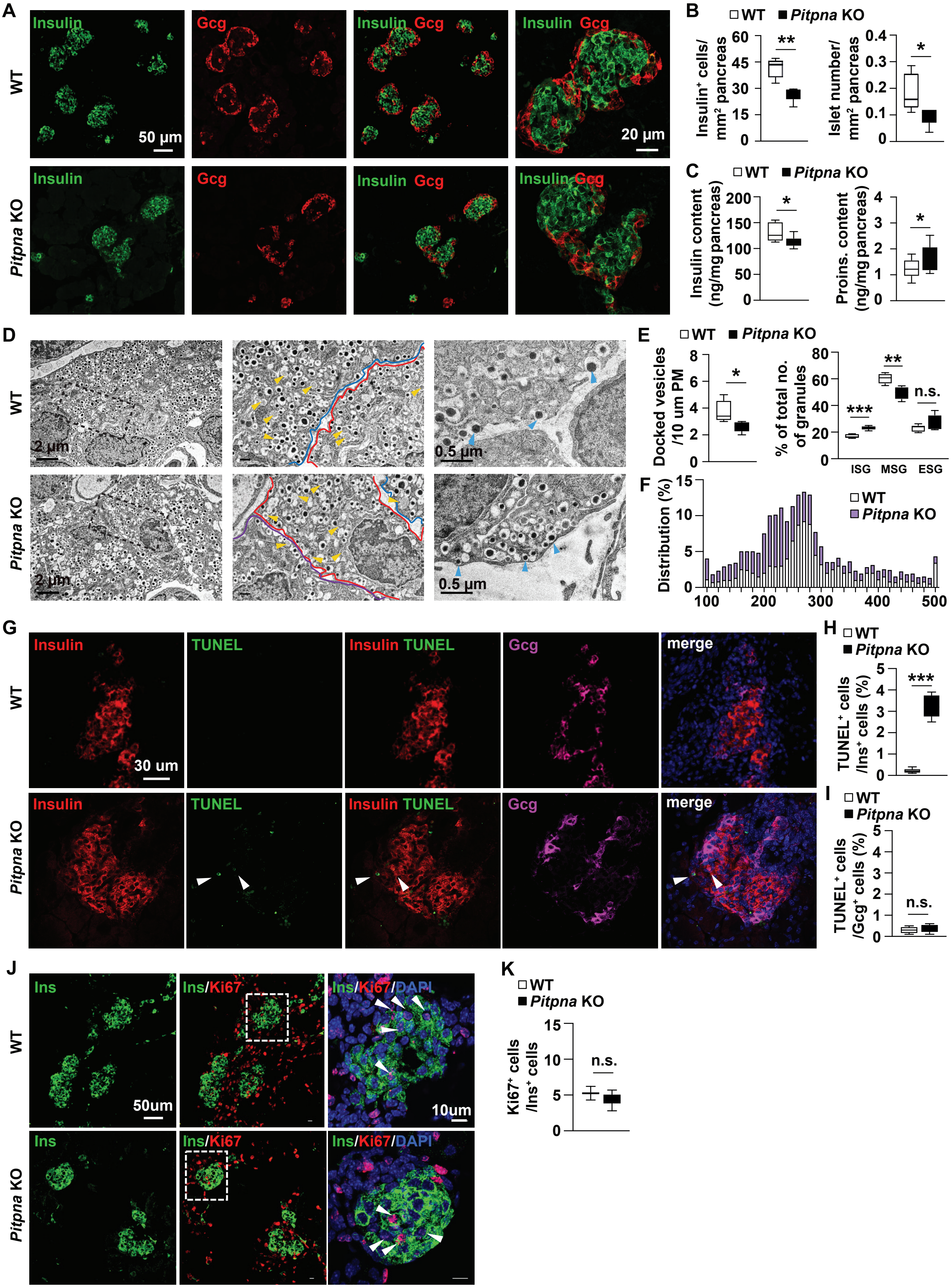


Yeh et al., 2022
Figure S2

### FIGURE S3

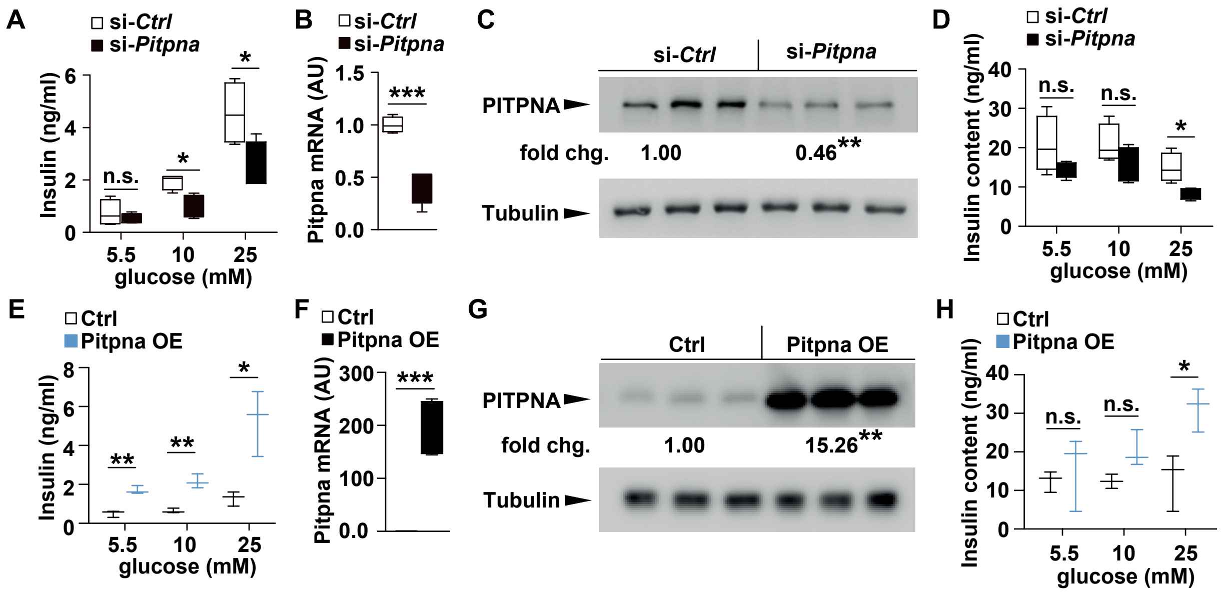


Yeh et al., 2022
Figure S3

### FIGURE S4

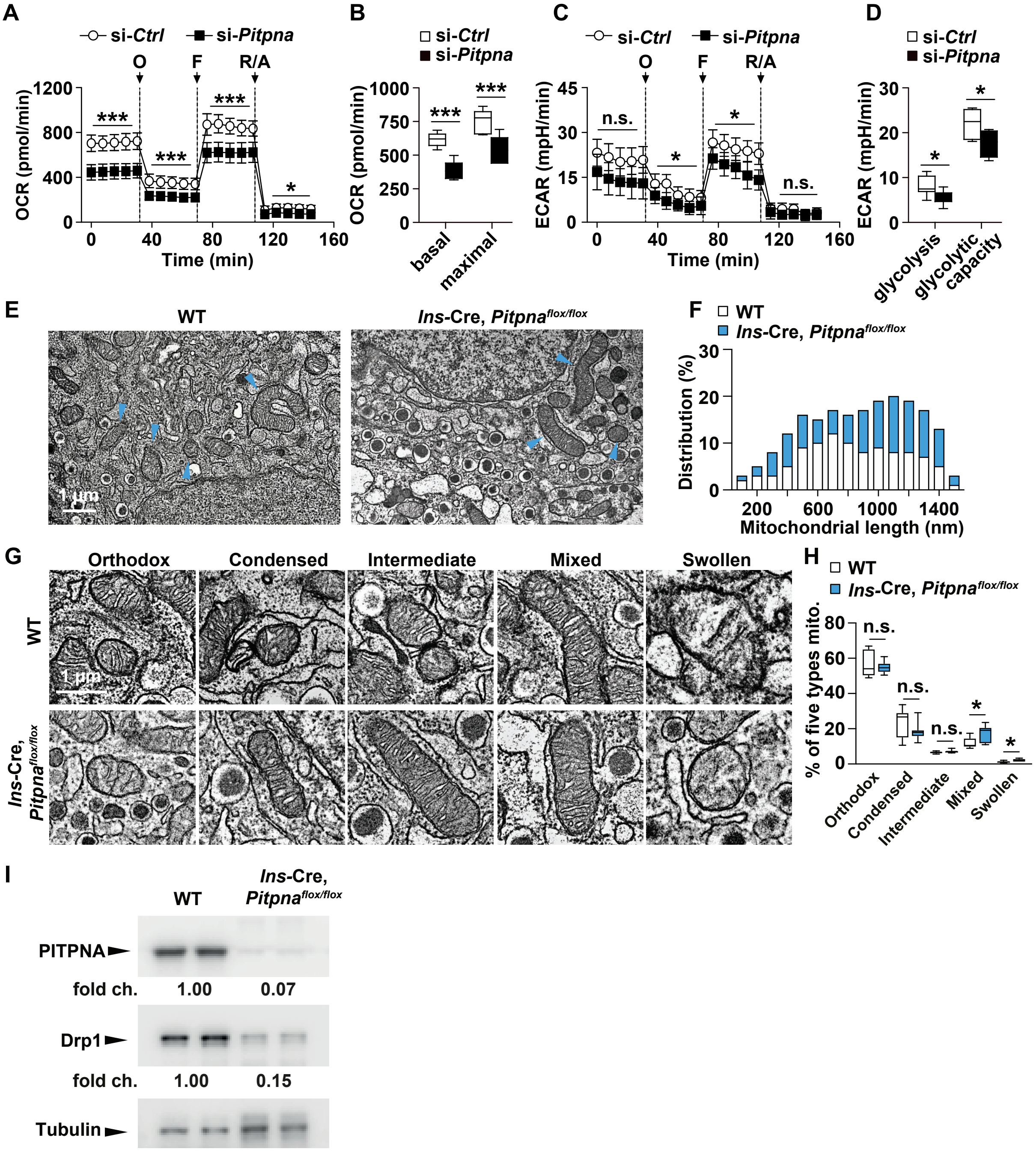


Yeh et al., 2022
Figure S4

### FIGURE S5

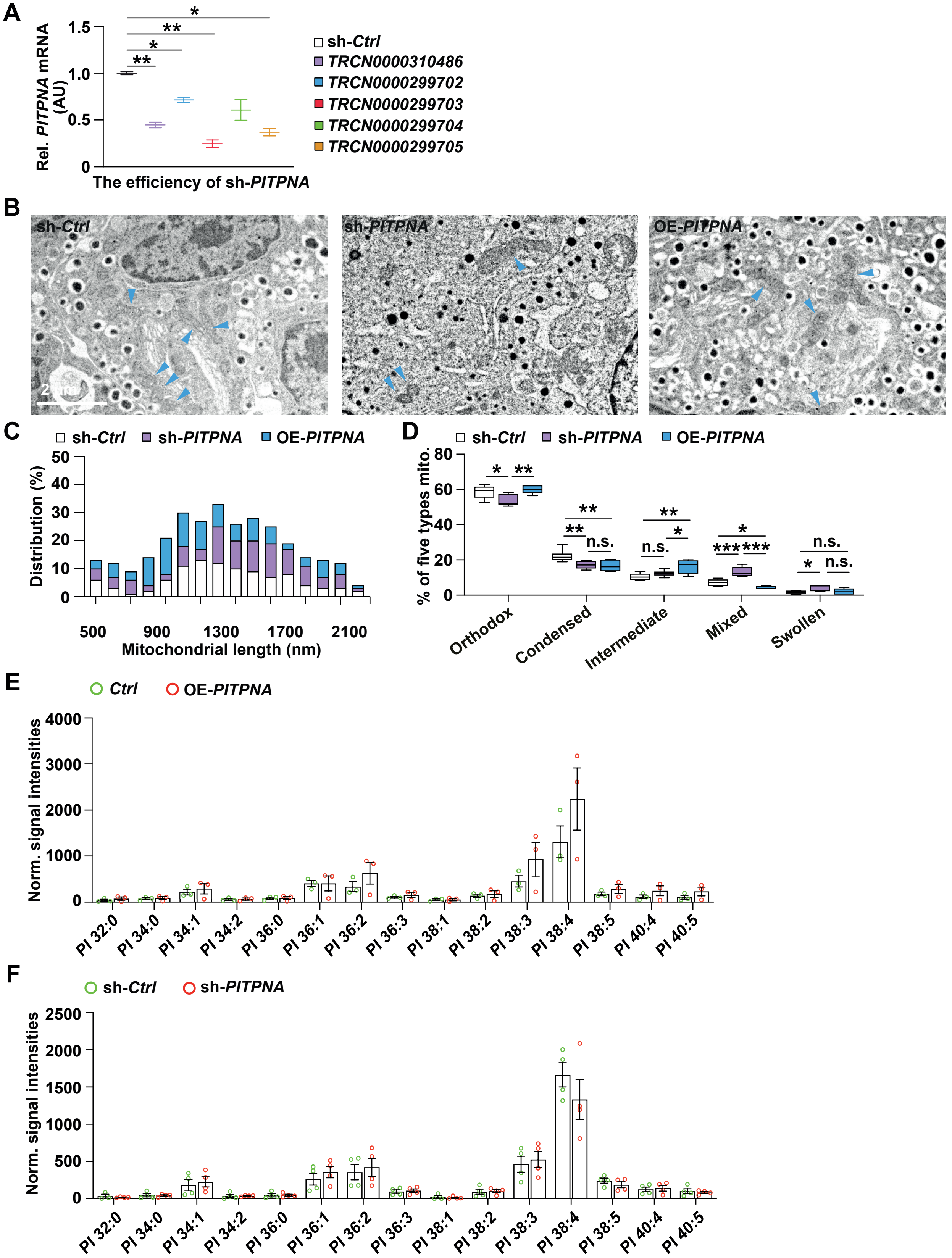


Yeh et al., 2022
Figure S5

### FIGURE S6

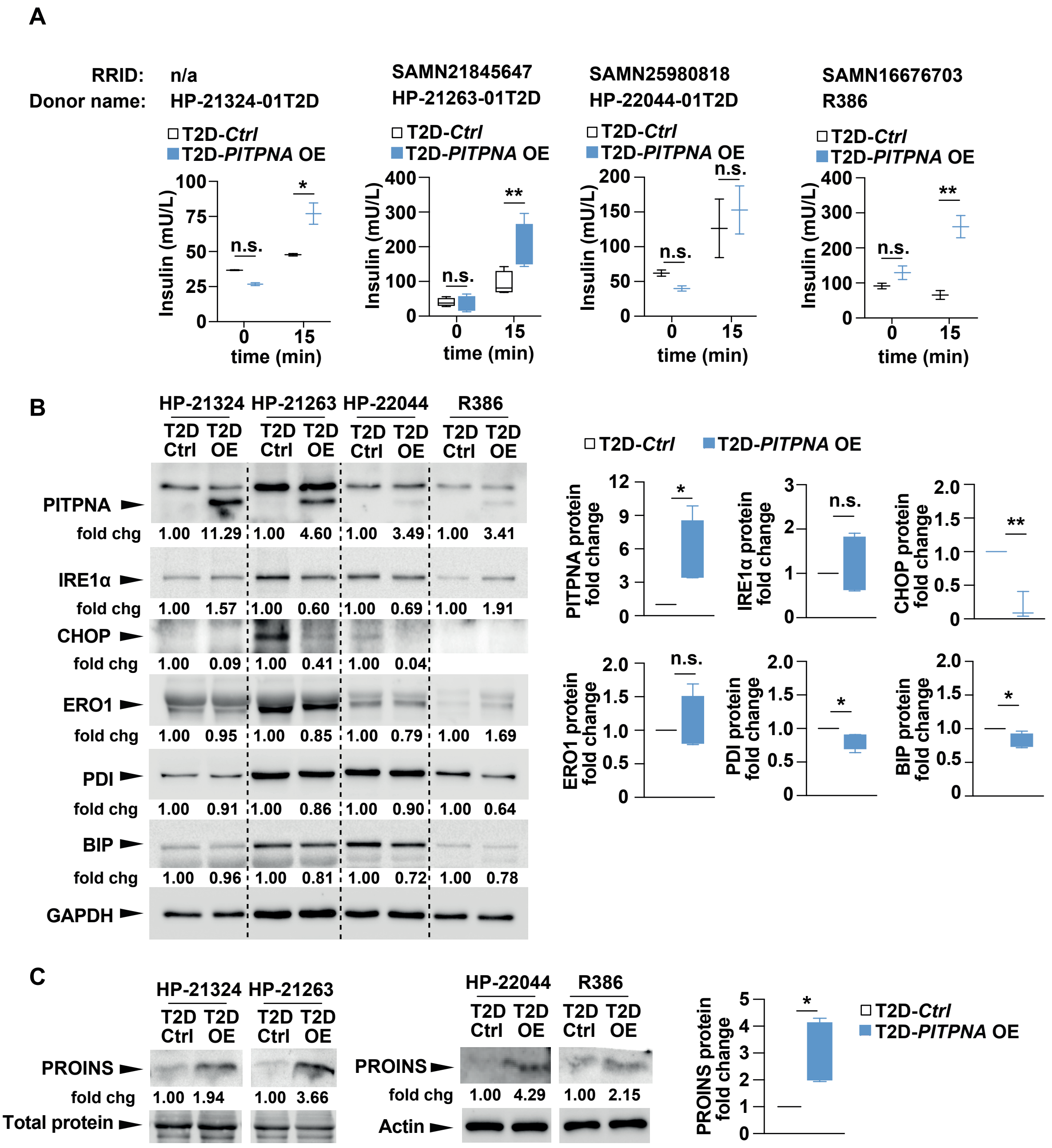


Yeh et al., 2022
Figure S6

### FIGURE S8

Fig.4A

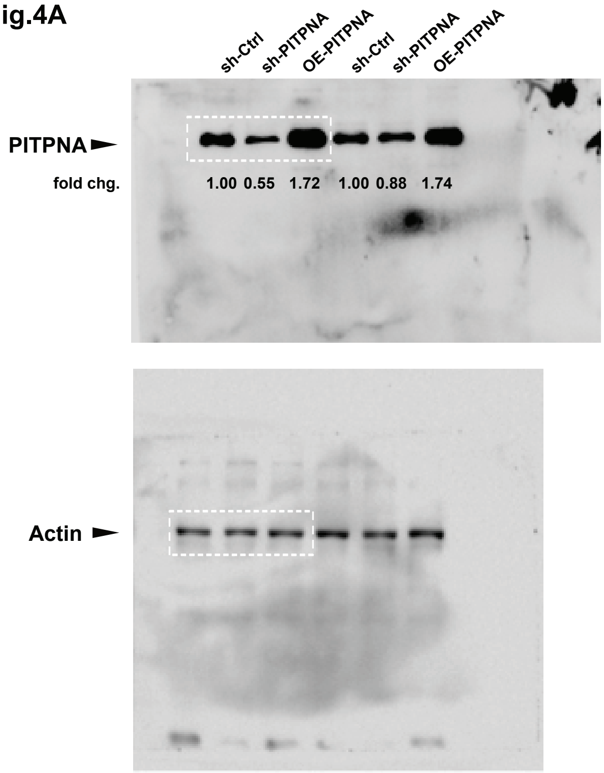


Fig.5F

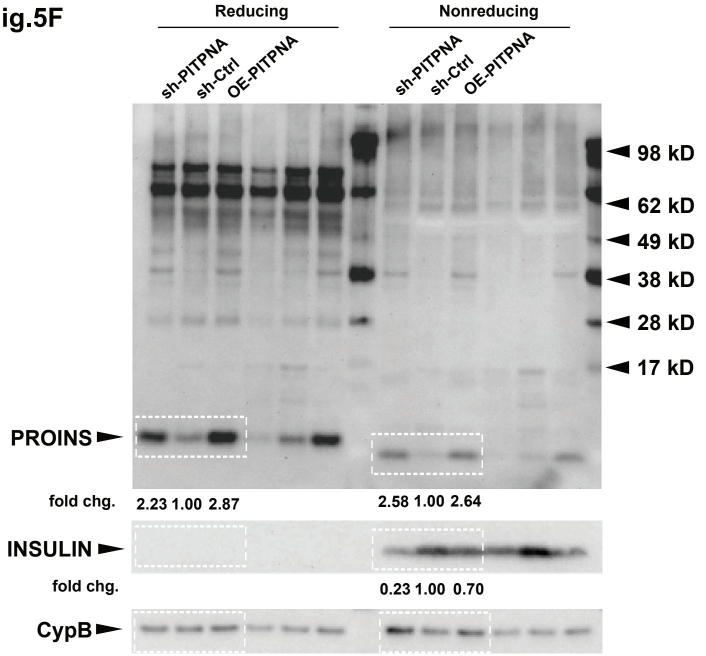


Fig.5G

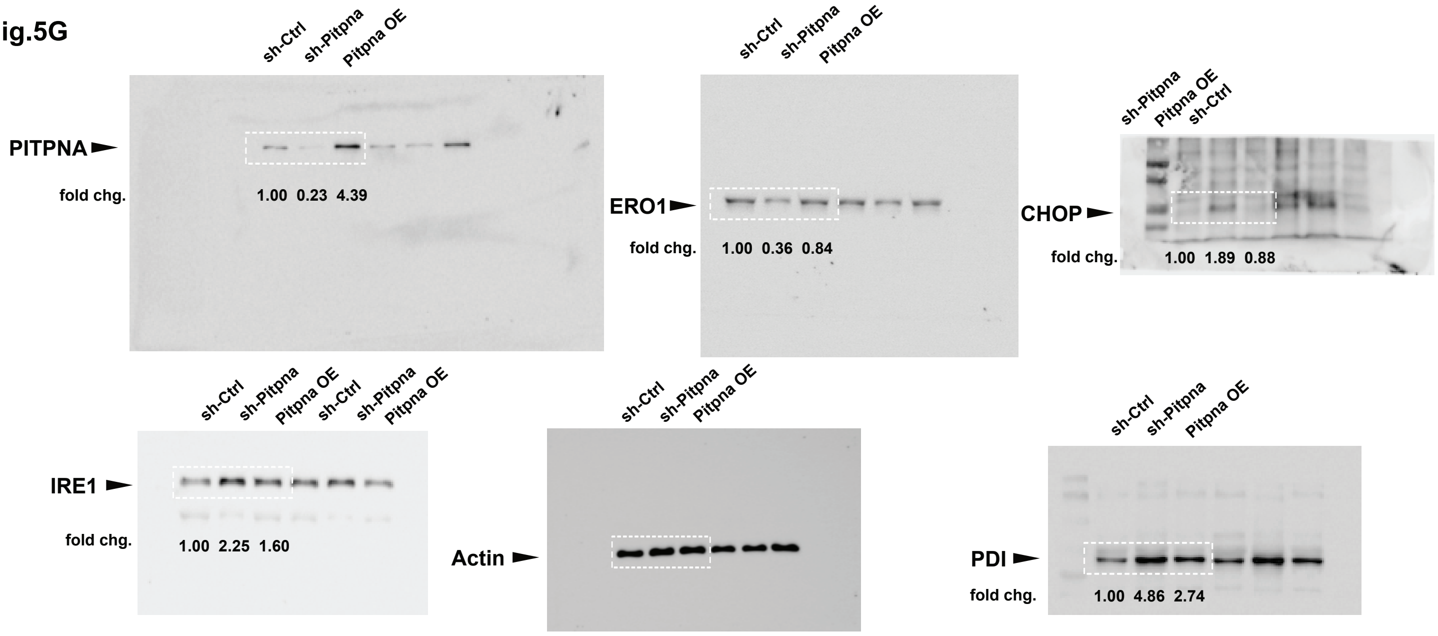


Fig.7A

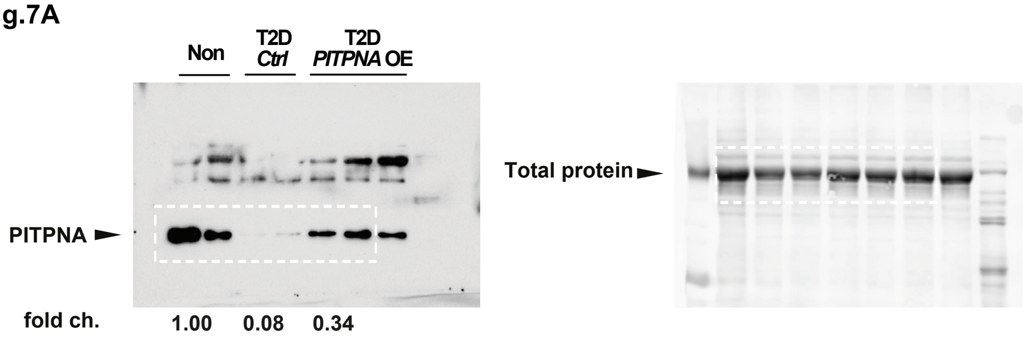


Note: Dashed white lines identify cropped areas

Yeh et al., 2022
Figure S8

### FIGURE S11

Fig.S6B

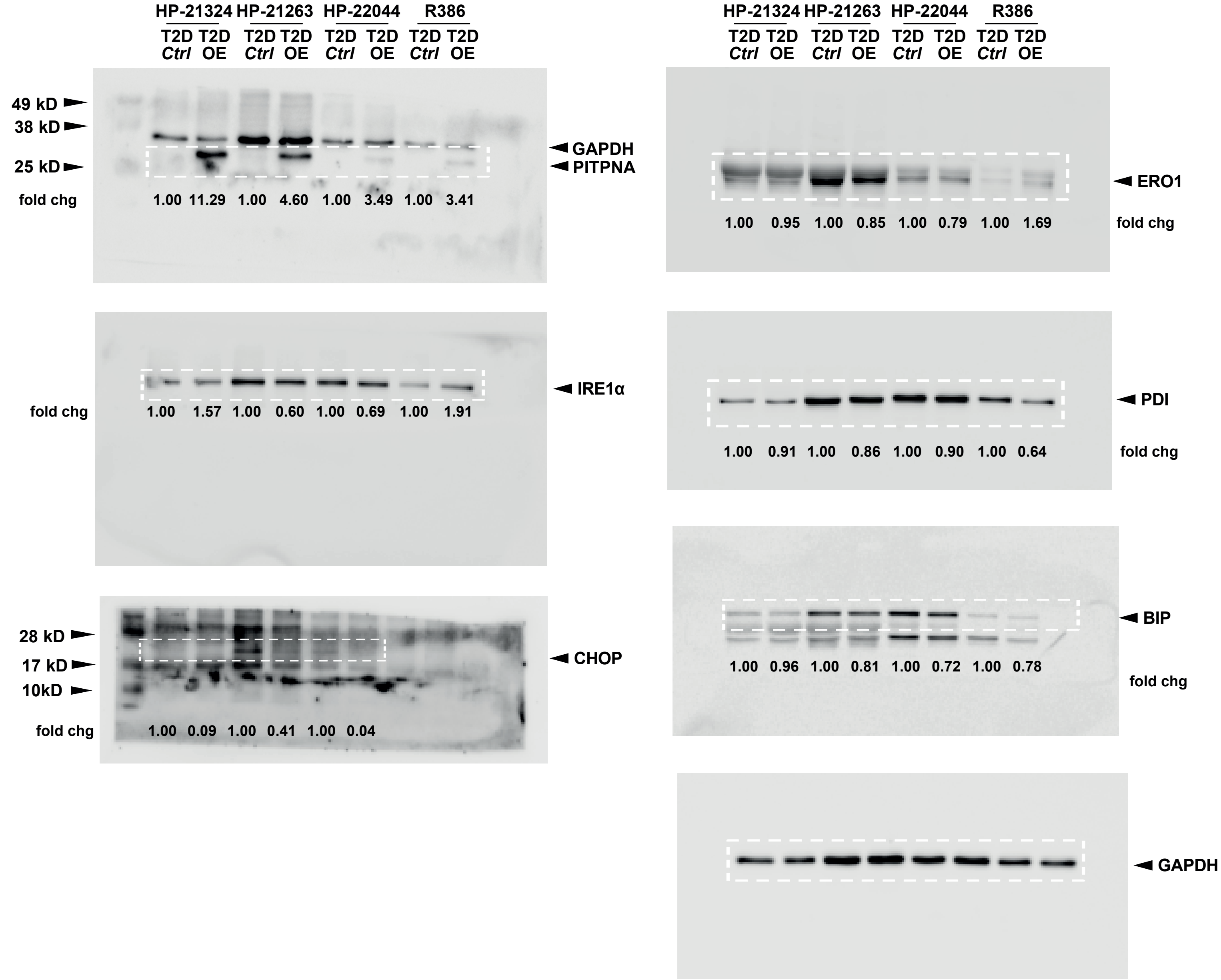


Note: Dashed white lines identify cropped areas

Yeh et al., 2022
Figure S11

### FIGURE S12

Fig.S6C

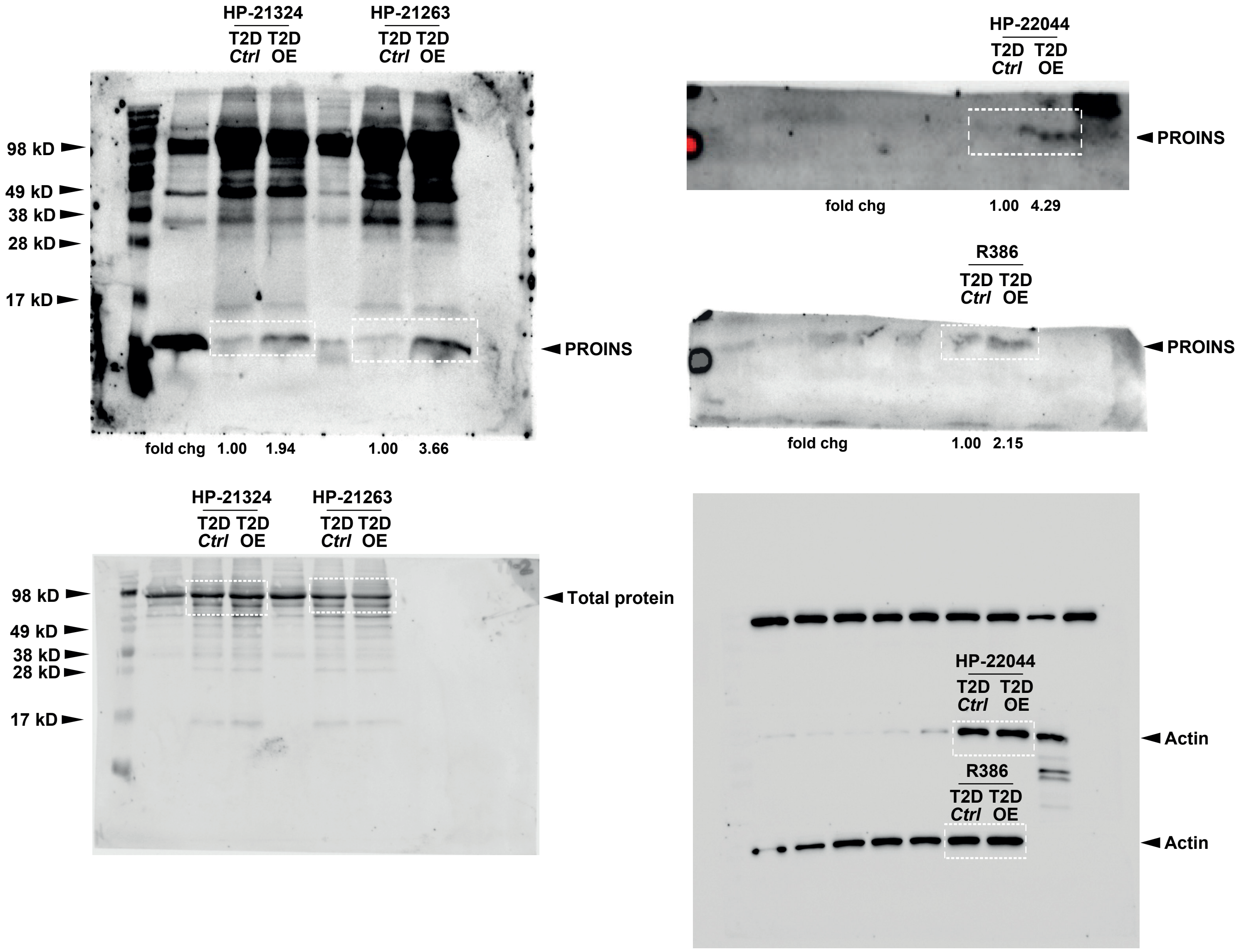


Note: Dashed white lines identify cropped areas

Yeh et al., 2022
Figure S12
