## Supplementary material for "Restoration of PITPNA in Type 2 diabetic human islets reverses pancreatic beta-cell dysfunction": FIGURE S7

Fig.1I

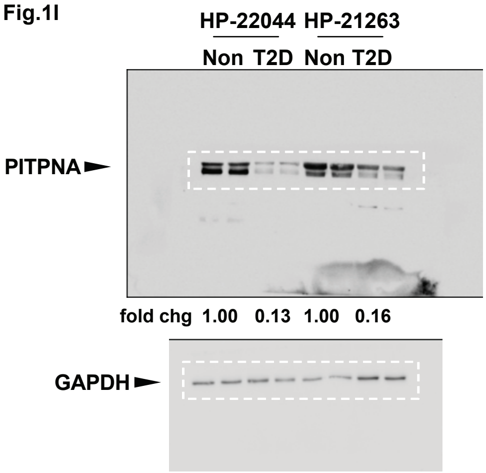


Fig.2A

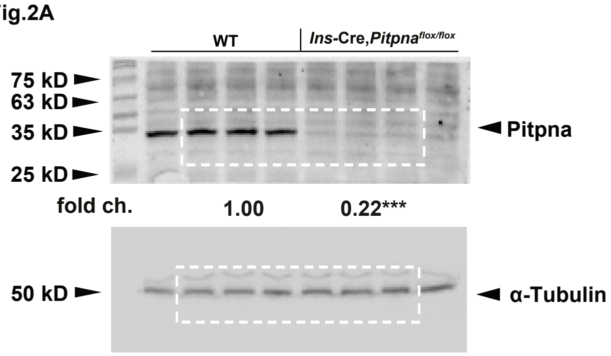


Fig.3D

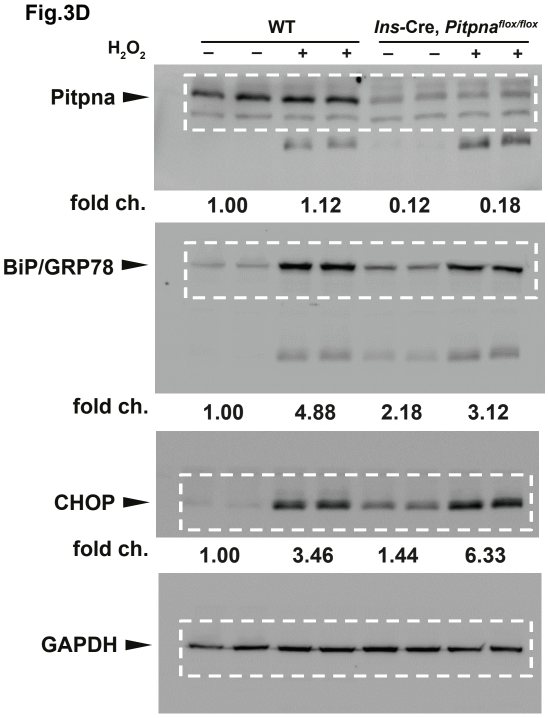


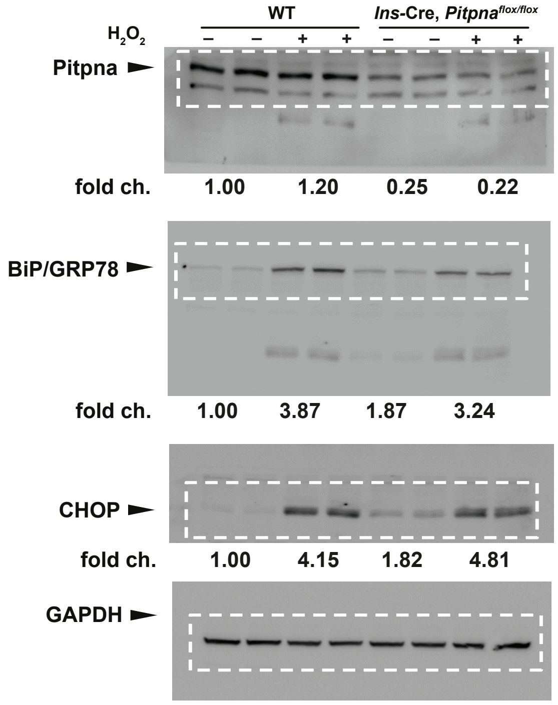


| | | WT | | Ins-Cre, *Pitpna*$^{flox/flox}$ | |
|---|---|---|---|---|---|
| | $H_2O_2$ | – – | + + | – – | + + |
| Pitpna mean (n=4) | fold ch. | 1.00 | 1.15 | 0.19 | 0.17 |
| | s.e.m. | 0.07 | 0.11 | 0.02 | 0.03 |
| | *P*-value | | 0.43 | <0.0001 | <0.0001 |
| BiP/GRP78 mean (n=4) | fold ch. | 1.00 | 4.78 | 2.21 | 3.30 |
| | s.e.m. | 0.04 | 0.63 | 0.32 | 0.21 |
| | *P*-value | | <0.0001 | 0.148 | 0.004 |
| CHOP mean (n=4) | fold ch. | 1.00 | 3.65 | 1.52 | 6.57 |
| | s.e.m. | 0.06 | 0.22 | 0.11 | 0.70 |
| | *P*-value | | 0.001 | 0.7589 | <0.0001 |

* additional panels shown above not in manuscript

Note: Dashed white lines identify cropped areas

Yeh et al., 2022
Figure S7
