## Supplementary material for "Restoration of PITPNA in Type 2 diabetic human islets reverses pancreatic beta-cell dysfunction": FIGURE S9

Fig.S1C

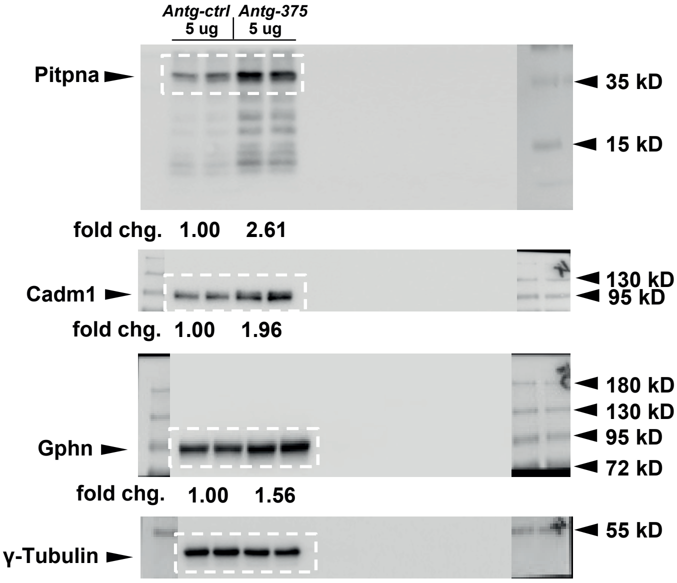


| | | *Antg-ctrl* 5 ug | *Antg-375* 5 ug |
|---|---|---|---|
| Pitpna mean (n=5) | fold ch. | 1.00 | 2.13 |
| | s.e.m. | 0.05 | 0.23 |
| | *P*-value | | 0.0012 |
| Cadm1 mean (n=5) | fold ch. | 1.00 | 1.84 |
| | s.e.m. | 0.02 | 0.09 |
| | *P*-value | | < 0.0001 |
| Gphn mean (n=4) | fold ch. | 1.00 | 1.38 |
| | s.e.m. | 0.06 | 0.10 |
| | *P*-value | | 0.0086 |

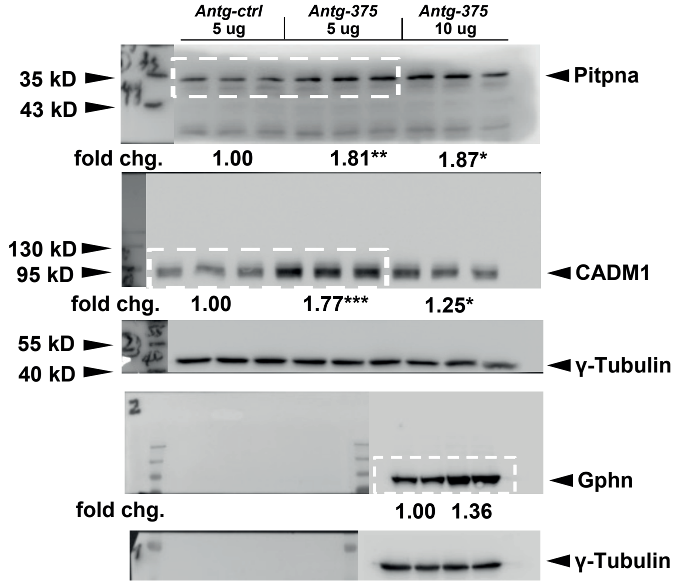


* additional panels shown above not in manuscript

Fig.S1D

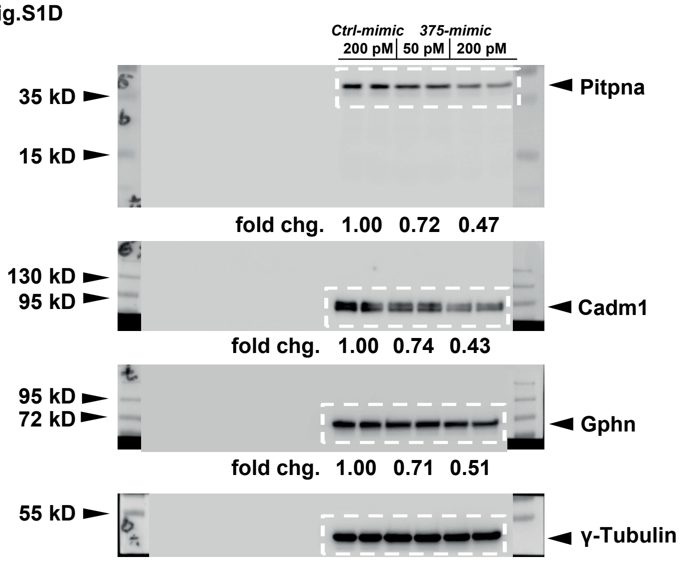


| | | *Ctrl-mimic* 200 pM | *375-mimic* 200 pM |
|---|---|---|---|
| Pitpna mean (n=5) | fold ch. | 1.00 | 0.61 |
| | s.e.m. | 0.04 | 0.08 |
| | *P*-value | | 0.0024 |
| Cadm1 mean (n=5) | fold ch. | 1.00 | 0.50 |
| | s.e.m. | 0.05 | 0.04 |
| | *P*-value | | < 0.0001 |

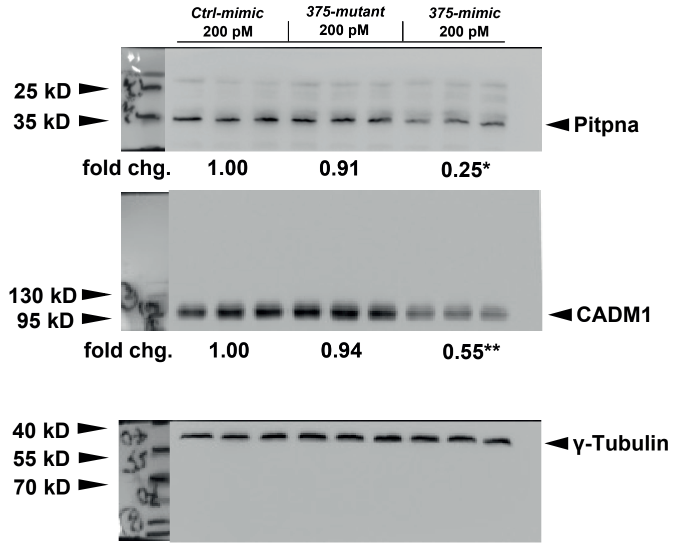


* additional panels shown above not in manuscript

Note: Dashed white lines identify cropped areas

Yeh et al., 2022
Figure S9
