## Supplementary material for "Restoration of PITPNA in Type 2 diabetic human islets reverses pancreatic beta-cell dysfunction": FIGURE S10

Fig.S3C

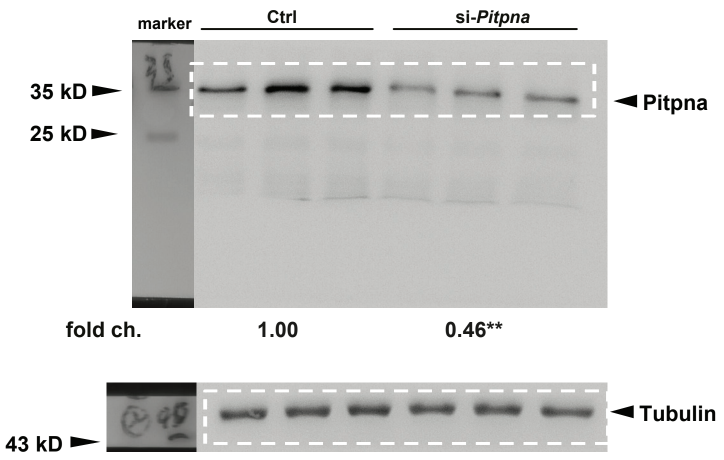


Fig.S3G

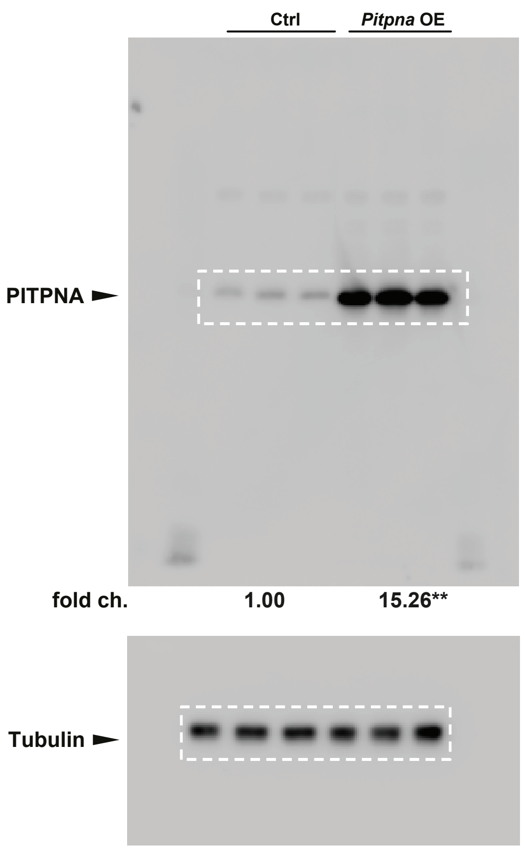


Fig.S4E

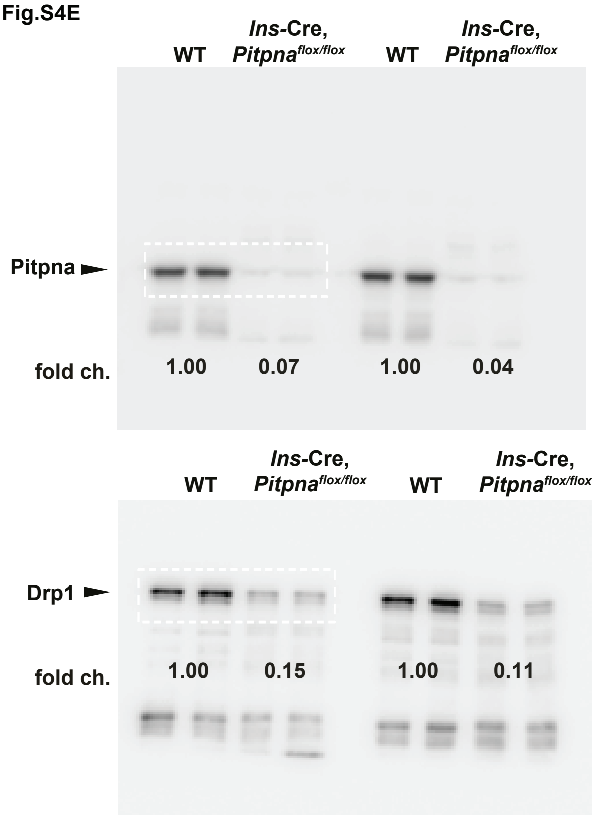


Fig.S4E

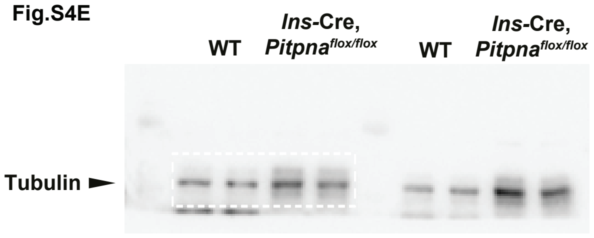


Fig.S4E

| | | WT | *Ins*-Cre, *Pitpna*$^{flox/flox}$ |
|---|---|---|---|
| Pitpna mean (n=4) | fold ch. | 1.00 | 0.05 |
| | s.e.m. | 0.056 | 0.002 |
| | *P*-value | | < 0.0001 |
| Drp1 mean (n=4) | fold ch. | 1.00 | 0.12 |
| | s.e.m. | 0.057 | 0.0005 |
| | *P*-value | | = 0.0068 |

Note: Dashed white lines identify cropped areas

Yeh et al., 2022
Figure S10
