## Supplementary material for "Restoration of PITPNA in Type 2 diabetic human islets reverses pancreatic beta-cell dysfunction": TABLE S1

| **TABLE S1. Demographic information for human pancreatic islet donors.** | | | | | | | | | | | | | | | | | | |
| --- | --- | --- | --- | --- | --- | --- | --- | --- | --- | --- | --- | --- | --- | --- | --- | --- | --- | --- |
| **Donor RRID** | **Name** | **Donor Type** | **Islet Procurement Center** | | **Age (yr)** | | **Gender** | | **Race** | **BMI (kg/m^2^)** | | **HbA1c** | **Islet Viability** | | **Islet Purity** | | **Cause of death** | **Notes** |
| SAMN25980818 | HP-22044-01T2D | T2D | Prodo | | 55 | | Male | | Asian | 30.7 | | 6.5 | 95 | | 90 | | Cerebrovascular/Stroke |  |
|  | HP-21342-01T2D | T2D | Prodo | | 48 | | Male | | Hispanic/Latino | 39.2 | | 7.1 | 95 | | 50 | | Anoxia | T2D for 0-5yrs. |
| SAMN21845647 | HP-21263-01T2D | T2D | IIDP/Scharp-Lacy | | 62 | | Male | | Caucasian | 34.9 | | 6.8 | 95 | | 95 | | Cerebrovascular/Stroke | T2D for 0-5yrs. Diet and oral medication treated. |
| SAMN25936113 | R432 | T2D | University of Alberta | | 66 | | Female | | n/a | 36.5 | | 8.6 | n/a | | 75 | | n/a | T2D for 25yrs. Diet and metformin treated. No insulin treatment. |
| SAMN16676703 | R386 | T2D | University of Alberta | | 43 | | Female | | n/a | 35.8 | | n/a | n/a | | 50 | | n/a | T2D for 17yrs. |
| SAMN21845649 | HP-21263-01T2D | T2D | Prodo | | 62 | | Male | | Caucasian | 34.6 | | 6.8 | 95 | | 95 | | Cerebrovascular/Stroke | T2D for 5yrs. Diet treated. |
| SAMN27279243 | HP-22089-01 | Non-diabetic | Prodo | | 48 | | Male | | Hispanic/Latino | 18.6 | | 5.3 | 95 | | 87 | | Cerebrovascular/Stroke |  |
|  | HP-22076-01 | Non-diabetic | Prodo | | 36 | | Male | | Caucasian | 23.2 | | 5.3 | 95 | | 90 | | Anoxia |  |
|  | HP-22084-01 | Non-diabetic | Prodo | | 44 | | Female | | Caucasian | 30.4 | | 5.2 | 95 | | 90 | | Cerebrovascular/Stroke |  |
| SAMN25690319 | HP-22034-01 | Non-diabetic | Prodo | | 49 | | Male | | Caucasian | 32.5 | | 5.5 | 95 | | 90 | | Anoxia |  |
|  | HP-22030-01 | Non-diabetic | Prodo | | 36 | | Male | | Asian | 25.6 | | 5.4 | 95 | | 85 | | Anoxia |  |
|  | HP-22021-01 | Non-diabetic | Prodo | | 52 | | Female | | African American | 23.2 | | 5.5 | 95 | | 90 | | Cerebrovascular/Stroke |  |
|  | HP-21337-01 | Non-diabetic | Prodo | | 26 | | Male | | Hispanic/Latino | 30.3 | | 5.9 | 95 | | 95 | | Head Trauma |  |
|  | HP-20021-01 | Non-diabetic | Prodo | | 52 | | Male | | Caucasian | 29.6 | | 5.4 | 95 | | 90 | | Cerebrovascular/Stroke |  |
| SAMN14255441 | HP-20060-01 | Non-diabetic | Prodo | | 48 | | Male | | Caucasian | 25.6 | | 5.9 | 95 | | 95 | | Anoxia |  |
|  | HP-21280-01 | Non-diabetic | Prodo | | 53 | | Male | | Caucasian | 24.54 | | 5.8 | 95 | | 90 | | Head Trauma |  |
| SAMN21032331 | HP-21239-01 | Non-diabetic | Prodo | | 40 | | Male | | African American | 27.2 | | 5.7 | 95 | | 90 | | Head Trauma |  |
| SAMN26177826 | HP-22052-01 | Non-diabetic | Prodo | | 52 | | Male | | Caucasian | 25.4 | | 4.2 | 95 | | 95 | | Anoxia |  |
| SAMN22021186 | HP-21272-01 | Non-diabetic | Prodo | | 39 | | Male | | Caucasian | 29 | | 5.4 | 95 | | 93 | | Anoxia |  |
|  | HP-21287-01 | Non-diabetic | Prodo | | 44 | | Male | | Caucasian | 27.6 | | 5.2 | 95 | | 85 | | Head Trauma |  |
|  | HP-21286-01 | Non-diabetic | Prodo | | 30 | | Male | | Hispanic/Latino | 27.6 | | 5.3 | 95 | | 85 | | Cerebrovascular/Stroke |  |
| SAMN21244110 | HP-21247-01 | Non-diabetic | IIDP/Scharp-Lacy | | 53 | | Male | | Caucasian | 31.2 | | 5.9 | 95 | | 95 | | Anoxia |  |
| SAMN20926064 | 1224 | Non-diabetic | IIDP/So.Calif.Islet Res.Ctr. | | 62 | | Female | | African American | 26.8 | | n/a | 96 | | 90 | | Cerebrovascular/Stroke |  |
| SAMN20923891 | HIL003 | Non-diabetic | IIDP/Loyola Medical Ctr. | | 45 | | Male | | Caucasian | 50.6 | | 5.4 | 95 | | 93 | | Anoxia |  |
| SAMN20064638 | HU1222 | Non-diabetic | IIDP/So.Calif.Islet Res.Ctr. | | 66 | | Male | | Asian | 29.9 | | n/a | 96 | | 85 | | Cerebrovascular/Stroke |  |
| **Human islet material related to Figure 1H** | | | |  | |  | |  |  | |  |  | |  | |  |  | |
|  | HP-11256-01T2D | T2D | Prodo | | 42 | | Male | | Asian | 30.2 | | 10 | 95 | | 70 | | Cerebrovascular/Stroke |  |
|  | HP-13346-01T2D | T2D | Prodo | | 29 | | Female | | Caucasian | 38.7 | | 9.9 | 95 | | 90 | | Cerebrovascular/Stroke | Metformin treated. |
|  | HP-14034-01T2D | T2D | Prodo | | 55 | | Male | | Hispanic | 22.4 | | 6.8 | 95 | | 90 | | Head Trauma |  |
|  | HP-14009-01T2D | T2D | Prodo | | 52 | | Female | | Caucasian | 32.5 | | 8.2 | 95 | | 85 | | Anoxia |  |
| SAMN08776519 | HP-13324-01T2D | T2D | Prodo | | 44 | | Female | | Hispanic | 30.5 | | 7.1 | 95 | | 90 | | Anoxia |  |
|  | HP-14005-01 | Non-diabetic | Prodo | | 53 | | Female | | Caucasian | 20.3 | | n/a | 95 | | 90 | | Cerebrovascular/Stroke |  |
| SAMN08776518 | HP-13326-01 | Non-diabetic | Prodo | | 38 | | Female | | Caucasian | 33.1 | | 5.5 | 95 | | 95 | | Cerebrovascular/Stroke |  |
| SAMN08784599 | HP-12303-01 | Non-diabetic | Prodo | | 41 | | Female | | Caucasian | 35.5 | | 5.3 | 95 | | 90 | | Head Trauma |  |
| SAMN08784445 | HP-13041-01 | Non-diabetic | Prodo | | 59 | | Female | | African American | 22.6 | | n/a | 95 | | 95 | | Brain Tumor |  |
|  | HP-13049-01 | Non-diabetic | Prodo | | 27 | | Female | | Caucasian | 21.7 | | 5.1 | 95 | | 80 | | Anoxia |  |
|  | HP-13076-01 | Non-diabetic | Prodo | | 64 | | Male | | Caucasian | 24.7 | | n/a | 95 | | 95 | | Cerebrovascular/Stroke |  |
| SAMN08784305 | HP-13159-01 | Non-diabetic | Prodo | | 52 | | Male | | Caucasian | 34.3 | | 5.7 | 95 | | 90 | | Anoxia |  |
|  | HP-13088-01 | Non-diabetic | Prodo | | 19 | | Male | | Hispanic | 32.5 | | 5.4 | 95 | | 85 | | Gunshot Wound |  |
|  | HP-14025-01 | Non-diabetic | Prodo | | 39 | | Female | | Caucasian | 28.2 | | 5.3 | 95 | | 85 | | Cerebrovascular/Stroke |  |
|  | HP-14019-01 | Non-diabetic | Prodo | | 34 | | Male | | Hispanic | 28.5 | | n/a | 95 | | 85 | | Head Trauma |  |
|  | HP-14073-01 | Non-diabetic | Prodo | | 43 | | Male | | Caucasian | 25.8 | | n/a | 95 | | 85 | | Head Trauma |  |
| SAMN08783903 | HP-13297-01 | Non-diabetic | Prodo | | 44 | | Female | | Asian/Filipino | 23.4 | | 5.1 | 95 | | 80-85 | | Cerebrovascular/Stroke |  |
|  | HP-14059-01 | Non-diabetic | Prodo | | 28 | | Male | | Caucasian | 25.3 | | 5.6 | 95 | | 80-85 | | Anoxia |  |
|  | HP-14053-01 | Non-diabetic | Prodo | | 43 | | Male | | Hispanic | 30.8 | | 5 | 95 | | 85 | | Head Trauma |  |
|  | HP-14039-01 | Non-diabetic | Prodo | | 37 | | Female | | Hispanic | 38 | | 5.5 | 95 | | 85-90 | | Cerebrovascular/Stroke |  |
