## Supplementary material for "Restoration of PITPNA in Type 2 diabetic human islets reverses pancreatic beta-cell dysfunction": TABLE S2

| Table S2. Summary of statistical analyses | | | |
| --- | --- | --- | --- |
| Figure | Sample size (n) | Statistical Test | Values |
| 1E | Non (n=51)  Pre-diabetes (n=27)  Disbetes (n=11) | Ordinary one-way ANOVA  Turkey’s multiple comparisons test | Summary: F=3.543, P =0.0332  Multiple comparison:  Non vs Pre-diabetes: P=0.7248  NOD vs Diabetes: P=0.025  Diabetes vs Pre-diabetes: P=0.1278 |
| 1F | N=77 | Linear regression | F=2.871, R sequared =0.03687 |
| 1G | N=89 | Linear regression | F=1.239, R sequared =0.01404 |
| 1H | Non (n=15)  T2D (n=5) | Two-tailed unpaired Student’s t-test | t=4.125, P=0.0006 |
| 2A | Wild-type (n=3)  *Ins-*Cre, *Pitpna^flox/flox^* (n=3) | Two-tailed unpaired Student’s t-test | t=16.2, P<0.0001 |
| 2B | WT (n=6)  Pitpna KO (n=6)  WT (n=5)  Pitpna KO (n=6) | Two-tailed unpaired Student’s t-test | Fed: t=3.687, P=0.0042  Fast: t=0.2001, P=0.8454  Plasma ins.: t=2.569, P=0.0303 |
| 2C | Wild-type (n=6)  *Ins-*Cre, *Pitpna^flox/flox^* (n=6) | Two-way repeated-measure ANOVA  Post-hoc multiple comparisons test (Sidak's) | Interaction: F=3.387, P=0.0272  Time: F=52.20, P<0.0001  Genotype: F=101.3, P<0.0001 |
| 2D | Wild-type (n=6)  *Ins-*Cre, *Pitpna^flox/flox^* (n=6) | Two-way repeated-measure ANOVA  Post-hoc multiple comparisons test (Sidak's) | Interaction: F=2.343, P=0.0675  Time: F=46.58, P<0.0001  Genotype: F=15.14, P=0.0003 |
| 2E | Wild-type (n=5)  *Ins-*Cre, *Pitpna^flox/flox^* (n=5) | Two-tailed unpaired Student’s t-test | 2.8: t=0.9584, P=0.3695  16.7: t=3.506, P=0.008  KCl: t=3.169, P=0.0132 |
| 2G | Wild-type (n=8)  *Ins-*Cre, *Pitpna^flox/flox^* (n=8) | Two-tailed unpaired Student’s t-test | t=2.979, P=0.01 |
| 2H | Wild-type (n=8)  *Ins-*Cre, *Pitpna^flox/flox^* (n=8) | Two-tailed unpaired Student’s t-test | ISG: t=7.551, P<0.0001  MSG: t=5.951, P<0.0001  CCG: t=0.3304, P=0.746  ESG: t=2.462, P=0.0274 |
| 2J | Wild-type (n=5)  *Ins-*Cre, *Pitpna^flox/flox^* (n=5) | Two-tailed unpaired Student’s t-test | Islets number: t=3.151, P=0084  Ins^+^ cells: t=8.775, P<0.0001  Beta-cells: t=3.191, P=0.0128 |
| 3B | Wild-type (n=6)  *Ins-*Cre, *Pitpna^flox/flox^* (n=6) | Two-tailed unpaired Student’s t-test | t=4.867, P=0.0007 |
| 3C | Wild-type (n=6)  *Ins-*Cre, *Pitpna^flox/flox^* (n=6) | Two-tailed unpaired Student’s t-test | t=0.2388, P=0.8161 |
| 3E | Wild-type (n=5)  *Ins-*Cre, *Pitpna^flox/flox^* (n=5) | Two-tailed unpaired Student’s t-test | Pitpna: t=12.35, P<0.0001  PC1/3: t=8.768, P<0.0001  PC2: t=3.021, P=0.0165  CPE: t=2.509, P=0.0364  CGA: t=2.115, P=0.0673  CGB: t=1.184, P=0.2704 |
| 4B | sh-Ctrl (n=4)  sh-PITPNA (n=4)  OE-PITPNA (n=4) | Ordinary one-way ANOVA  Turkey’s multiple comparisons test | 2.8 mM glc.Summary: F=3.937, P =0.0591  Multiple comparison:  sh-Ctrl vs sh-PITPNA: P=0.7839  sh-Ctrl vs OE-PITPNA: P=0.1625  sh-PITPNA vs OE-PITPNA: P=0.0581  25 mM glc.Summary: F=137, P<0.0001  Multiple comparison:  sh-Ctrl vs sh-PITPNA: P<0.0001  sh-Ctrl vs OE-PITPNA: P=0.0008  sh-PITPNA vs OE-PITPNA: P<0.0001 |
| 4C | sh-Ctrl (n=5)  sh-PITPNA (n=5) | Two-way repeated-measure ANOVA  Post-hoc multiple comparisons test (Sidak's) | Interaction: F=2.494, P<0.0001  Time: F=0.4961, P=0.5381  Genotype: F=8.047, P=0.0176 |
| 4H | sh-Ctrl (n=40)  sh-PITPNA (n=43)  Ctrl (n=50)  OE-PITPNA (n=40) | Two-way ANOVA  Tukey’s multiple comparisons test | Summary: F=11.67, P<0.0001  Multiple comparison:  sh-Ctrl vs sh-PITPNA: P=0.0003  Ctrl vs OE-PITPNA: P=0.001 |
| 5B | sh-Ctrl (n=4)  sh-PITPNA (n=4)  OE-PITPNA (n=4) | Ordinary one-way ANOVA  Turkey’s multiple comparisons test | Summary: F=32.18, P<0.0001  Multiple comparison:  sh-Ctrl vs sh-PITPNA: P=0.0399  sh-Ctrl vs OE-PITPNA: P=0.0019  sh-PITPNA vs OE-PITPNA: P<0.0001 |
| 5C | sh-Ctrl (n=4)  sh-PITPNA (n=4)  OE-PITPNA (n=4) | Ordinary one-way ANOVA  Turkey’s multiple comparisons test | Summary: F=24.99, P=0.0002  Multiple comparison:  sh-Ctrl vs sh-PITPNA: P=0.0029  sh-Ctrl vs OE-PITPNA: P=0.1217  sh-PITPNA vs OE-PITPNA: P=0.0002 |
| 5D | sh-Ctrl (n=4)  sh-PITPNA (n=4)  OE-PITPNA (n=4) | Ordinary one-way ANOVA  Turkey’s multiple comparisons test | ISG Summary: F=54.34, P=0.0006  Multiple comparison:  sh-Ctrl vs sh-PITPNA: P=0.0195  sh-Ctrl vs OE-PITPNA: P=0.0512  sh-PITPNA vs OE-PITPNA: P=0.0043  MSG Summary: F=25.97, P=0.0131  Multiple comparison:  sh-Ctrl vs sh-PITPNA: P=0.0019  sh-Ctrl vs OE-PITPNA: P=0.3926  sh-PITPNA vs OE-PITPNA: P=0.0295  CCG Summary: F=5.43, P=0.0995  Multiple comparison:  sh-Ctrl vs sh-PITPNA: P=0.0365  sh-Ctrl vs OE-PITPNA: P=0.9449  sh-PITPNA vs OE-PITPNA: P=0.0894  ESG Summary: F=2.6, P=0.0817  Multiple comparison:  sh-Ctrl vs sh-PITPNA: P=0.1434  sh-Ctrl vs OE-PITPNA: P=0.2095  sh-PITPNA vs OE-PITPNA: P=0.1599 |
| 5E | sh-Ctrl (n=7)  sh-PITPNA (n=6)  OE-PITPNA (n=5) | Two-tailed unpaired Student’s t-test | sh-Ctrl vs sh-PITPNA: t=2.373, P=0.037  sh-Ctrl vs OE-PITPNA: t=3.33, P=0.0076  sh-PITPNA vs OE-PITPNA: t=0.2873, P=0.7804 |
| 6B | sh-Ctrl (n=5)  sh-PITPNA (n=7) | Two-tailed unpaired Student’s t-test | sh-Ctrl vs sh-PITPNA: t=3.702, P=0.0041 |
| 6D | sh-Ctrl (n=23)  sh-PITPNA (n=10) | Two-tailed unpaired Student’s t-test | sh-Ctrl vs sh-PITPNA: t=7.16, P<0.0001 |
| 6E | Control (n=3)  OE-PITPNA (n=3) | Two-tailed unpaired Student’s t-test | PIP 34:0: t=1.166, P=0.3085  PIP 34:1: t=1.505, P=0.2069  PIP 34:2: t=0.0527, P=0.9604  PIP 36:0: t=2.921, P=0.0432  PIP 36:1: t=2.452, P=0.0703  PIP 36:2: t=2.53, P=0.0646  PIP 36:3: t=2.777, P=0.05  PIP 36:4: t=2.346, P=0.0788  PIP 38:2: t=2.316, P=0.0815  PIP 38:3: t=2.469, P=0.069  PIP 38:4: t=3.013, P=0.0394  PIP 38:5: t=3.711, P=0.3604 |
| 6E | Control (n=3)  PITPNA OE (n=3) | Two-tailed unpaired Student’s t-test | t=2.88, P=0.045 |
| 6F | sh-Control (n=4)  sh-PITPNA (n=4) | Two-tailed unpaired Student’s t-test | PIP 34:1: t=0.4875, P=0.6432  PIP 34:2: t=0.0595, P=0.958  PIP 36:0: t=0.2991, P=0.793  PIP 36:1: t=0.42, P=0.6891  PIP 36:2: t=0.0192, P=0.9853  PIP 36:3: t=0.0937, P=0.9284  PIP 36:4: t=1.384, P=0.3006  PIP 38:2: t=0.0728, P=0.9485  PIP 38:3: t=0.0521, P=0.9601  PIP 38:4: t=0.1757, P=0.8663  PIP 38:5: t=0.0241, P=0.9815  PIP 40:4: t=0.137, P=0.8955  PIP 40:5: t=0.2352, P=0.8219 |
| 6F | sh-Control (n=4)  sh-PITPNA (n=4) | Two-tailed unpaired Student’s t-test | t=2.872, P=0.0284 |
| 7B | T2D (n=4)  T2D-PITPNA OE (n=4) | Two-tailed unpaired Student’s t-test | 0 min: t=0.386, P=0.7128  5 min: t=5.27, P>1  15 min: t=2.685, P=0.363 |
| 7C | T2D (n=15)  T2D-PITPNA OE (n=15) | Two-tailed unpaired Student’s t-test | t=6.851, P<0.0001 |
| 7F | Non (n=4)  T2D (n=4)  T2D-PITPNA OE (n=4) | Ordinary one-way ANOVA  Turkey’s multiple comparisons test | Summary: F=40.85, P=0.0019  Multiple comparison:  Non vs T2D: P=0.0068  Non vs T2D-PITPNA OE: P=0.0418  T2D vs T2D-PITPNA OE: P=0.0247 |
| 7G | Non (n=4)  T2D (n=4)  T2D-PITPNA OE (n=4) | Ordinary one-way ANOVA  Turkey’s multiple comparisons test | Summary: F=10.2, P=0.0495  Multiple comparison:  Non vs T2D: P=0.0235  Non vs T2D-PITPNA OE: P=0.6311  T2D vs T2D-PITPNA OE: P=0.0324 |
| 7H | Non (n=4)  T2D (n=4)  T2D-PITPNA OE (n=4) | Ordinary one-way ANOVA  Turkey’s multiple comparisons test | ISG Summary: F=12.59, P=0.0038  Multiple comparison:  Non vs T2D: P=0.0035  Non vs T2D-PITPNA OE: P=0.3567  T2D vs T2D-PITPNA OE: P=0.0269  MSG Summary: F=27.71, P=0.0047  Multiple comparison:  Non vs T2D: P=0.0124  Non vs T2D-PITPNA OE: P=0.0213  T2D vs T2D-PITPNA OE: P=0.0104  CCG Summary: F=0.6778, P=0.6262  Multiple comparison:  Non vs T2D: P=0.0119  Non vs T2D-PITPNA OE: P=0.4167  T2D vs T2D-PITPNA OE: P=0.5409  ESG Summary: F=3.797, P=0.0938  Multiple comparison:  Non vs T2D: P=0.1124  Non vs T2D-PITPNA OE: P=0.0971  T2D vs T2D-PITPNA OE: P=0.8798 |
| S1A | Ctrl-mimic (n=4)  375-mimic (n=4) | Two-tailed unpaired Student’s t-test | Wild-type 3’UTR:  t=3.216, P=0.0182  MUT 3’UTR:  t=0.0000, P=1.0000 |
| S1B | Antg-ctrl (n=4)  Antg-375 (n=4) | Two-tailed unpaired Student’s t-test | t=11.88, P<0.0001 |
| S1E | Wild type (n=4)  *Ins-*Cre, *Ago2^flox/flox^* (n=4) | Two-tailed unpaired Student’s t-test | Pitpna: t=10.49, P<0.0001  Cadm1: t=5.999, P=0.0010  Gphn: t=4.216, P=0.0056  Ago2: t=14.87, P<0.0001 |
| S1F | Antg-Ctrl (n=6)  antg-375 (n=6) | Two-tailed unpaired Student’s t-test | PI 36:1: t=8.052, P=0.0017  PI 36:2: t=5.832, P=0.003  PI 36:3: t=2.811, P=0.059  PI 36:4: t=4.013, P=0.019  PI 38:1: t=1.873, P=0.16  PI 38:2: t=4.933, P=0.0087  PI 38:3: t=3.898, P=0.016  PI 38:4: t=3.088, P=0.034  PI 38:5: t=3.742, P=0.019  PI 38:6: t=1.885, P=0.18  PI 40:4: t=2.692, P=0.078  PI 40:5: t=4.664, P=0.0088  PI 40:6: t=2.874, P=0.048 |
| S1F | antg-Ctrl (n=6)  antg-375 (n=6) | Two-tailed unpaired Student’s t-test | PI total: t=3.444, P=0.0063 |
| S2B | WT (n=5)  Pitpna KO (n=5)  WT (n=5)  Pitpna KO (n=5) | Two-tailed unpaired Student’s t-test  Two-tailed unpaired Student’s t-test | Insulin^+^ cells: t=4.673, P=0.0016  Islet number: t=2.53, P=0.0353 |
| S2C | WT (n=7)  Pitpna KO (n=7)  WT (n=13)  Pitpna KO (n=8) | Two-tailed unpaired Student’s t-test  Two-tailed unpaired Student’s t-test | Insulin content: t=2.289, P=0.041  Proinsulin content: t=2.067, P=0.0526 |
| S2E | WT (n=5)  Pitpna KO (n=15)  WT (n=5)  Pitpna KO (n=5) | Two-tailed unpaired Student’s t-test  Two-tailed unpaired Student’s t-test | Docked vesicles: t=2.81, P=0.0228  ISG: t=6.45, P=0.0002  MSG: t=3.829, P=0.005  ESG: t=1.552, P=0.1593 |
| S2H | WT (n=5)  Pitpna KO (n=5) | Two-tailed unpaired Student’s t-test | TUNEL^+^ insulin cells/: t=12.25, P<0.0001 |
| S2I | WT (n=5)  Pitpna KO (n=5) | Two-tailed unpaired Student’s t-test | TUNEL^+^ Gcg cells: t=0.7184, P=0.4929 |
| S2K | WT (n=5)  Pitpna KO (n=5) | Two-tailed unpaired Student’s t-test | Ki67^+^ Gcg cells/: t=1.989, P=0.0701 |
| S3A | si-Ctrl (n=4)  si-Pitpna (n=4) | Two-tailed unpaired Student’s t-test | 5.5 mM: t=0.8533, P=0.4262  10 mM: t=3.683, P=0.0103  25 mM: t=2.744, P=0.0335 |
| S3B | si-Ctrl (n=4)  si-Pitpna (n=4) | Two-tailed unpaired Student’s t-test | t=5.906, P=0.0010 |
| S3C | si-Ctrl (n=3)  si-Pitpna (n=3) | Two-tailed unpaired Student’s t-test | t=6.319, P=0.0032 |
| S3D | si-Ctrl (n=4)  si-Pitpna (n=4) | Two-tailed unpaired Student’s t-test | 5.5 mM: t=1.65, P=0.15  10 mM: t=1.611, P=0.1584  25 mM: t=3.303, P=0.0163 |
| S3E | si-Ctrl (n=3)  si-Pitpna (n=3) | Two-tailed unpaired Student’s t-test | 5.5 mM: t=7.728, P=0.0015  10 mM: t=6.81, P=0.0024  25 mM: t=3.39, P=0.0275 |
| S3F | si-Ctrl (n=4)  si-Pitpna (n=4) | Two-tailed unpaired Student’s t-test | t=6.764, P=0.0005 |
| S3G | si-Ctrl (n=3)  si-Pitpna (n=3) | Two-tailed unpaired Student’s t-test | t=6.555, P=0.0028 |
| S3H | si-Ctrl (n=3)  si-Pitpna (n=3) | Two-tailed unpaired Student’s t-test | 5.5 mM: t=0.5385, P=0.6188  10 mM: t=2.709, P=0.0536  25 mM: t=3.975, P=0.0165 |
| S4D | Wild-type (n=7)  *Ins-*Cre, *Pitpna^flox/flox^* (n=7) | Two-tailed unpaired Student’s t-test | Orthodox: t=0.4214, P=0.6809  Condensed: t=1.502, P=0.1589  Intermediate: t=1.495, P=0.1608  Mixed: t=2.465, P=0.0298  Swollen: t=2.493, P=0.0283 |
| S5C | sh-Ctrl (n=7)  sh-PITPNA (n=7)  OE-PITPNA (n=7) | Ordinary one-way ANOVA  Turkey’s multiple comparisons test | Orthodox Summary: F=7.842, P=0.0036  Multiple comparison:  sh-Ctrl vs sh-PITPNA: P=0.0266  sh-Ctrl vs OE-PITPNA: P=0.6246  sh-PITPNA vs OE-PITPNA: P=0.0036  Condensed Summary: F=8.321, P=0.0028  Multiple comparison:  sh-Ctrl vs sh-PITPNA: P=0.0079  sh-Ctrl vs OE-PITPNA: P=0.0053  sh-PITPNA vs OE-PITPNA: P=0.9808  Intermediate Summary: F=9.148, P=0.0018  Multiple comparison:  sh-Ctrl vs sh-PITPNA: P=0.4725  sh-Ctrl vs OE-PITPNA: P=0.0016  sh-PITPNA vs OE-PITPNA: P=0.0217  Mixed Summary: F=38.71, P<0.0001  Multiple comparison:  sh-Ctrl vs sh-PITPNA: P<0.0001  sh-Ctrl vs OE-PITPNA: P=0.0451  sh-PITPNA vs OE-PITPNA: P<0.0001  Swollen Summary: F=4.309, P=0.0296  Multiple comparison:  sh-Ctrl vs sh-PITPNA: =0.0237  sh-Ctrl vs OE-PITPNA: P=0.4757  sh-PITPNA vs OE-PITPNA: P=0.2209 |
| S5D | Control (n=3)  OE-PITPNA (n=3) | Two-tailed unpaired Student’s t-test | PI 32:0: t=0.913, P=0.4129  PI 34:0: t=0.3226, P=0.7631  PI 34:1: t=0.5446, P=0.615  PI 34:2: t=0.2573, P=0.8096  PI 36:0: t=0.0592, P=0.9556  PI 36:1: t=0.00529, P=0.996  PI 36:2: t=1.132, P=0.3209  PI 36:3: t=0.8823, P=0.4275  PI 38:1: t=0.145, P=0.8917  PI 38:2: t=0.419, P=0.6968  PI 38:3: t=1.252, P=0.2786  PI 38:4: t=1.2228, P=0.2868  PI 38:5: t=1.032, P=0.3604  PI 40:4: t=1.138, P=0.3185  PI 40:5: t=1.193, P=0.2988 |
| S5E | sh-Control (n=4)  sh-PITPNA (n=4) | Two-tailed unpaired Student’s t-test | PI 32:0: t=0.7343, P=0.4957  PI 34:0: t=0.1111, P=0.9151  PI 34:1: t=4035, P=0.7005  PI 34:2: t=0.0838, P=0.9359  PI 36:0: t=0.0561, P=0.957  PI 36:1: t=0.849, P=0.4285  PI 36:2: t=0.42, P=0.6891  PI 36:3: t=0.3753, P=0.7203  PI 38:1: t=0.4944, P=0.642  PI 38:2: t=0.177, P=0.8653  PI 38:3: t=0.4125, P=0.6925  PI 38:4: t=1.052, P=0.3332  PI 38:5: t=1.038, P=0.3394  PI 40:4: t=0.3128, P=0.765  PI 40:5: t=0.4397, P=0.6756 |
| S6A | T2D (n=2)  T2D-PITPNA OE (n=2) | Two-way repeated-measure ANOVA  Post-hoc multiple comparisons test (Sidak's) | HP-21324-01T2D Summary: F=20.79, P=0.0449  Multiple comparison:  0 min: P=0.2638  15 min: P=0.0116  HP-21263-01T2D Summary: F=5.738, P=0.0536  Multiple comparison:  0 min: P=0.9830  15 min: P=0.0071  HP-22044-01T2D Summary: F=0.9998, P=0.4227  Multiple comparison:  0 min: P=0.8376  15 min: P=0.7799  R386 Summary: F=140.4, P=0.007  Multiple comparison:  0 min: P=0.4399  15 min: P=0.0046 |
| S6B | T2D (n=3-4)  T2D PITPNA OE (n=3-4) | Two-tailed unpaired Student’s t-test | IRE1a: t=0.6064, P=0.5664  CHOP: t=7.15, P=0.002  ERO1: t=0.3418, P=0.7442  Pdi: t=2.683, P=0.0364  Bip: t=3.495, P=0.0129  PI 40:5: t=1.193, P=0.2988 |
| S6C | T2D (n=4)  T2D PITPNA OE (n=4) | Two-tailed unpaired Student’s t-test | PROINS: t=3.498, P=0.0129 |
