## Supplementary material for "Restoration of PITPNA in Type 2 diabetic human islets reverses pancreatic beta-cell dysfunction": TABLE S4

| **TABLE S4. List of qRT-PCR primer sequences** | | | |  |
| --- | --- | --- | --- | --- |
| **Gene name** | | **Forward** | | **Reverse** |
| Mouse *Argonaute RISC catalytic subunit 2* (*Ago2*) | | ACCATGTACTCGGGAGCC | | TGCTCCACAATTTCCCTGTTCA |
| Mouse *Cell adhesion molecule 1* (*Cadm1*) | | ATGGCGAGTGCTGTGCTG | | TACGTGGAGGAACCAGGACT |
| Mouse *Chromogranin A* (*Cga*) | | AGCCAGACTACAGACCCACT | | GGACGCACTTCATCACCTTG |
| Mouse *Chromogranin B* (*Cgb*) | | GCAAGTCCTGAAGAAGAGTGG | | CATCGGCTGGGTCTCTTAGC |
| Mouse *Carboxypeptidase E* (*Cpe*) | | CGGCATCTCCTTCGAGTACC | | AATTCAGGTTCACCCGGCTC |
| Mouse *Gephyrin* (*Gphn*) | | GTCACTCCAGAGGCCACAAA | | TGGAGAGAGAGGAGGTGGTG |
| Mouse *Phosphatidylinositol transfer protein, alpha* (*Pitpna*) | | TATCGGGTCATCCTGCCTGT | | AGGTGGGTACTTTGCTCTGTA |
| Mouse *Proprotein convertase 1/3* (*Pcsk1*) | CCTCCTACAGCAGTGGTGATTACA | | GGGTCTCTGTGCAGTCATTGT | |
| Mouse *Proprotein convertase 2* (*Pcsk2*) | AGACAATGGGAAGACGGTTG | | TTGAAGCATAGCCGTCACAG | |
| Mouse *beta-Actin* | | GGCTGTATTCCCCTCCATCG | | CCAGTTGGTAACAATGCCATGT |
| Mouse *Glyceraldehyde-3-phosphate dehydrogenase* (*Gapdh*) | | AGGTCGGTGTGAACGGATTTG | | GGGGTCGTTGATGGCAACA |
