## Supplementary material for "Restoration of PITPNA in Type 2 diabetic human islets reverses pancreatic beta-cell dysfunction": TABLE S3

| **A. PI sh-Ctrl vs sh-PITPNA** | | | | | | | | | | | | | | | | | |
| --- | --- | --- | --- | --- | --- | --- | --- | --- | --- | --- | --- | --- | --- | --- | --- | --- | --- |
| **Peak Areas PI NH4 + adducts** | | | | | | | | | | | | | | | | | |
|  | **Internal Std 37:4 100 ng** | **SUM** | **32 0** | **34 0** | **34 1** | **34 2** | **36 0** | **36 1** | **36 2** | **36 3** | **38 1** | **38 2** | **38 3** | **38 4** | **38 5** | **40 4** | **40 5** |
| **sh-Ctrl-1** | 158,803 | 3,823,865 | 3,645 | 27,054 | 118,209 | 1,311 | 18,218 | 169,356 | 283,159 | 85,101 | 559 | 50,575 | 351,440 | 2,325,288 | 235,395 | 99,368 | 55,187 |
| **sh-Ctrl-2** | 119,696 | 5,457,864 | 14,307 | 53,092 | 324,851 | 17,291 | 44,295 | 445,756 | 630,504 | 147,752 | 1,115 | 125,838 | 726,389 | 2,237,170 | 373,734 | 204,311 | 111,461 |
| **sh-Ctrl-3** | 54,618 | 2,959,733 | 45,980 | 57,175 | 187,371 | 51,636 | 57,206 | 233,548 | 296,831 | 78,379 | 36,906 | 100,987 | 370,359 | 1,093,706 | 141,600 | 102,499 | 105,550 |
| **sh-Ctrl-4** | 520,299 | 13,555,546 |  | 58,456 | 272,250 | 139,826 | 130,919 | 746,013 | 907,587 | 311,244 |  | 258,104 | 1,803,483 | 6,856,959 | 1,284,116 | 393,656 | 392,933 |
| **sh-PITPNA-1** | 117,990 | 6,407,499 | 20,013 | 69,200 | 455,337 | 51,115 | 49,205 | 606,281 | 805,930 | 184,003 | 3,086 | 160,519 | 870,970 | 2,461,174 | 296,834 | 245,892 | 127,940 |
| **sh-PITPNA-2** | 133,614 | 4,945,333 | 18,614 | 62,156 | 379,351 | 48,923 | 50,025 | 592,299 | 758,415 | 182,741 | 5,804 | 154,266 | 904,268 | 1,077,970 | 347,449 | 257,361 | 105,691 |
| **sh-PITPNA-3** | 91,076 | 2,832,248 | 24,025 | 41,551 | 129,070 | 36,628 | 74,576 | 281,598 | 241,855 | 75,207 | 33,037 | 98,312 | 373,309 | 1,132,952 | 100,185 | 100,763 | 89,179 |
| **sh-PITPNA-4** | 361,538 | 8,300,578 | 33,444 | 58,651 | 322,110 | 85,536 | 82,461 | 589,592 | 629,764 | 195,492 | 26,191 | 146,347 | 1,018,230 | 4,312,831 | 459,060 | 169,170 | 171,698 |
| **Normalized to Internal Standard** | | | | | | | | | | | | | | | | | |
|  |  | **SUM** | **32 0** | **34 0** | **34 1** | **34 2** | **36 0** | **36 1** | **36 2** | **36 3** | **38 1** | **38 2** | **38 3** | **38 4** | **38 5** | **40 4** | **40 5** |
| **sh-Ctrl-1** |  | 2407.93 | 2.30 | 17.04 | 74.44 | 0.83 | 11.47 | 106.65 | 178.31 | 53.59 | 0.35 | 31.85 | 221.31 | 1464.26 | 148.23 | 62.57 | 34.75 |
| **sh-Ctrl-2** |  | 4559.76 | 11.95 | 44.36 | 271.40 | 14.45 | 37.01 | 372.41 | 526.75 | 123.44 | 0.93 | 105.13 | 606.86 | 1869.04 | 312.23 | 170.69 | 93.12 |
| **sh-Ctrl-3** |  | 5418.93 | 84.18 | 104.68 | 343.05 | 94.54 | 104.74 | 427.60 | 543.46 | 143.50 | 67.57 | 184.90 | 678.08 | 2002.45 | 259.25 | 187.66 | 193.25 |
| **sh-Ctrl-4** |  | 2605.34 |  | 11.24 | 52.33 | 26.87 | 25.16 | 143.38 | 174.44 | 59.82 |  | 49.61 | 346.62 | 1317.89 | 246.80 | 75.66 | 75.52 |
| **sh-PITPNA-1** |  | 5430.52 | 16.96 | 58.65 | 385.91 | 43.32 | 41.70 | 513.84 | 683.05 | 155.95 | 2.62 | 136.04 | 738.17 | 2085.91 | 251.57 | 208.40 | 108.43 |
| **sh-PITPNA-2** |  | 3701.20 | 13.93 | 46.52 | 283.91 | 36.62 | 37.44 | 443.29 | 567.61 | 136.77 | 4.34 | 115.46 | 676.77 | 806.78 | 260.04 | 192.61 | 79.10 |
| **sh-PITPNA-3** |  | 3109.78 | 26.38 | 45.62 | 141.72 | 40.22 | 81.88 | 309.19 | 265.55 | 82.58 | 36.27 | 107.95 | 409.89 | 1243.97 | 110.00 | 110.64 | 97.92 |
| **sh-PITPNA-4** |  | 2295.91 | 9.25 | 16.22 | 89.09 | 23.66 | 22.81 | 163.08 | 174.19 | 54.07 | 7.24 | 40.48 | 281.64 | 1192.91 | 126.97 | 46.79 | 47.49 |

TABLE S3. MASS SPECTROMETRY DATA SUMMARY

| **B. PI sh-Ctrl vs sh-PITPNA** | | | | | | | | | | | | | | | | | |
| --- | --- | --- | --- | --- | --- | --- | --- | --- | --- | --- | --- | --- | --- | --- | --- | --- | --- |
| **Peak Areas PI NH4 + adducts** | | | | | | | | | | | | | | | | | |
|  | **Internal Std 37:4 100 ng** | **SUM** | **32 0** | **34 0** | **34 1** | **34 2** | **36 0** | **36 1** | **36 2** | **36 3** | **38 1** | **38 2** | **38 3** | **38 4** | **38 5** | **40 4** | **40 5** |
| **Ctrl-1** | 54,618 | 2,959,733 | 45,980 | 57,175 | 187,371 | 51,636 | 57,206 | 233,548 | 296,831 | 78,379 | 36,906 | 100,987 | 370,359 | 1,093,706 | 141,600 | 102,499 | 105,550 |
| **Ctrl-2** | 119,431 | 3,814,892 | 14,301 | 96,752 | 172,856 | 44,408 | 134,015 | 600,714 | 306,438 | 107,545 | 84,003 | 192,180 | 479,385 | 1,199,539 | 166,843 | 130,843 | 85,072 |
| **Ctrl-3** | 319,983 | 7,608,801 | 62,182 | 106,914 | 540,023 | 125,094 | 106,656 | 903,202 | 664,940 | 252,951 | 45,311 | 237,126 | 861,905 | 2,943,549 | 438,247 | 154,019 | 166,681 |
| **OE-PITPNA-1** | 48,461 | 4,174,248 | 59,405 | 53,327 | 201,290 | 38,765 | 61,914 | 271,169 | 432,567 | 105,552 | 34,617 | 107,021 | 692,398 | 1,539,005 | 210,645 | 190,188 | 176,386 |
| **OE-PITPNA-2** | 46,215 | 3,434,017 | 44,626 | 59,730 | 177,088 | 43,470 | 53,586 | 266,283 | 384,276 | 96,470 | 41,165 | 124,267 | 527,070 | 1,207,429 | 138,303 | 140,560 | 129,696 |
| **OE-PITPNA-3** | 427,680 | 7,694,870 | 39,883 | 79,522 | 284,599 | 83,841 | 61,475 | 341,682 | 676,632 | 189,501 | 22,352 | 131,956 | 944,690 | 3,985,353 | 491,269 | 162,105 | 200,011 |
| **Normalized to Internal Standard** | | | | | | | | | | | | | | | | | |
|  |  | **SUM** | **32 0** | **34 0** | **34 1** | **34 2** | **36 0** | **36 1** | **36 2** | **36 3** | **38 1** | **38 2** | **38 3** | **38 4** | **38 5** | **40 4** | **40 5** |
| **Ctrl-1** |  | 5418.93 | 84.18 | 104.68 | 343.05 | 94.54 | 104.74 | 427.60 | 543.46 | 143.50 | 67.57 | 184.90 | 678.08 | 2002.45 | 259.25 | 187.66 | 193.25 |
| **Ctrl-2** |  | 3194.23 | 11.97 | 81.01 | 144.73 | 37.18 | 112.21 | 502.98 | 256.58 | 90.05 | 70.34 | 160.91 | 401.39 | 1004.38 | 139.70 | 109.56 | 71.23 |
| **Ctrl-3** |  | 2377.87 | 19.43 | 33.41 | 168.77 | 39.09 | 33.33 | 282.27 | 207.80 | 79.05 | 14.16 | 74.11 | 269.36 | 919.91 | 136.96 | 48.13 | 52.09 |
| **OE-PITPNA-1** |  | 8613.66 | 122.58 | 110.04 | 415.37 | 79.99 | 127.76 | 559.56 | 892.61 | 217.81 | 71.43 | 220.84 | 1428.78 | 3175.77 | 434.67 | 392.46 | 363.98 |
| **OE-PITPNA-2** |  | 7430.46 | 96.56 | 129.24 | 383.18 | 94.06 | 115.95 | 576.18 | 831.49 | 208.74 | 89.07 | 268.89 | 1140.46 | 2612.61 | 299.26 | 304.14 | 280.63 |
| **OE-PITPNA-3** |  | 1799.21 | 9.33 | 18.59 | 66.54 | 19.60 | 14.37 | 79.89 | 158.21 | 44.31 | 5.23 | 30.85 | 220.89 | 931.85 | 114.87 | 37.90 | 46.77 |

| **C. PIP sh-Ctrl vs sh-PITPNA** | | | | | | | | | | | | | | | | | |
| --- | --- | --- | --- | --- | --- | --- | --- | --- | --- | --- | --- | --- | --- | --- | --- | --- | --- |
| **PIP Peak Areas NH4+ Adducts** | |  |  |  |  |  |  |  |  |  |  |  |  |  |  |  |  |
|  | **Internal Std 37:4, 20 ng** | **SUM** | **32 0** | **34 0** | **34 1** | **34 2** | **36 0** | **36 1** | **36 2** | **36 3** | **36 4** | **38 2** | **38 3** | **38 4** | **38 5** | **40 4** | **40 5** |
| **sh-Ctrl-1** | 51,241 | 481,159 | 3,781 |  | 29,152 |  |  | 24,937 | 49,368 | 34,543 |  |  | 56,611 | 208,404 | 47,592 | 10,312 | 16,458 |
| **sh-Ctrl-2** | 35,036 | 590,746 | 12,488 |  | 27,560 |  |  | 22,676 | 41,001 | 33,188 |  |  | 70,749 | 300,986 | 55,278 | 10,943 | 15,876 |
| **sh-Ctrl-3** | 30,142 | 226,346 |  | 3,937 | 25,434 | 30,617 | 5,645 | 21,923 | 29,401 | 7,330 | 8,056 | 7,802 | 20,554 | 46,087 | 6,189 | 6,745 | 6,628 |
| **sh-Ctrl-4** | 104,648 | 742,751 |  |  | 18,709 | 145,594 | 2,537 | 27,116 | 33,587 | 14,558 | 35,303 | 7,395 | 61,312 | 340,602 | 38,925 | 8,456 | 8,656 |
| **sh-PITPNA-1** | 31,214 | 447,909 | 18,636 |  | 17,302 |  |  | 19,109 | 31,992 | 26,311 |  |  | 45,405 | 226,109 | 42,743 | 9,276 | 11,026 |
| **sh-PITPNA-2** | 33,931 | 461,734 | 19,324 |  | 17,305 |  |  | 19,374 | 32,601 | 25,091 |  |  | 57,043 | 229,952 | 40,362 | 8,826 | 11,857 |
| **sh-PITPNA-3** | 62,712 | 467,814 |  | 11,018 | 42,112 | 36,737 | 7,329 | 45,239 | 62,788 | 19,832 | 16,847 | 15,178 | 42,734 | 124,894 | 17,904 | 13,015 | 12,186 |
| **sh-PITPNA-4** | 111,164 | 817,132 |  |  | 35,026 | 211,547 | 4,489 | 46,505 | 47,320 | 392 | 19,609 | 11,615 | 73,714 | 316,364 | 31,261 | 9,993 | 9,297 |
| **Normalized to Internal Standard** | | | | | | | | | | | | | | | | | |
|  |  | **SUM** | **32 0** | **34 0** | **34 1** | **34 2** | **36 0** | **36 1** | **36 2** | **36 3** | **36 4** | **38 2** | **38 3** | **38 4** | **38 5** | **40 4** | **40 5** |
| **sh-Ctrl-1** |  | 187.80 | 1.48 |  | 11.38 |  |  | 9.73 | 19.27 | 13.48 |  |  | 22.10 | 81.34 | 18.58 | 4.02 | 6.42 |
| **sh-Ctrl-2** |  | 337.22 | 7.13 |  | 15.73 |  |  | 12.94 | 23.41 | 18.95 |  |  | 40.39 | 171.82 | 31.56 | 6.25 | 9.06 |
| **sh-Ctrl-3** |  | 150.19 |  | 2.61 | 16.88 | 20.32 | 3.75 | 14.55 | 19.51 | 4.86 | 5.35 | 5.18 | 13.64 | 30.58 | 4.11 | 4.48 | 4.40 |
| **sh-Ctrl-4** |  | 141.95 |  |  | 3.58 | 27.83 | 0.48 | 5.18 | 6.42 | 2.78 | 6.75 | 1.41 | 11.72 | 65.09 | 7.44 | 1.62 | 1.65 |
| **sh-PITPNA-1** |  | 286.99 | 11.94 |  | 11.09 |  |  | 12.24 | 20.50 | 16.86 |  |  | 29.09 | 144.88 | 27.39 | 5.94 | 7.06 |
| **sh-PITPNA-2** |  | 272.16 | 11.39 |  | 10.20 |  |  | 11.42 | 19.22 | 14.79 |  |  | 33.62 | 135.54 | 23.79 | 5.20 | 6.99 |
| **sh-PITPNA-3** |  | 149.19 |  | 3.51 | 13.43 | 11.72 | 2.34 | 14.43 | 20.02 | 6.32 | 5.37 | 4.84 | 13.63 | 39.83 | 5.71 | 4.15 | 3.89 |
| **sh-PITPNA-4** |  | 147.01 |  |  | 6.30 | 38.06 | 0.81 | 8.37 | 8.51 | 0.07 | 3.53 | 2.09 | 13.26 | 56.92 | 5.62 | 1.80 | 1.67 |

| **D. PIP Ctrl vs OE-PITPNA** | | | | | | | | | | | | | | | | | |
| --- | --- | --- | --- | --- | --- | --- | --- | --- | --- | --- | --- | --- | --- | --- | --- | --- | --- |
| **PIP Peak Areas NH4+ Adducts** | | | | | | | | | | | | | | | | | |
|  | **Internal Std 37:4, 20 ng** | **SUM** | **32 0** | **34 0** | **34 1** | **34 2** | **36 0** | **36 1** | **36 2** | **36 3** | **36 4** | **38 2** | **38 3** | **38 4** | **38 5** | **40 4** | **40 5** |
| **Ctrl-1** | 90,610 | 1,559,416 |  | 26,359 | 167,011 | 49,736 | 23,188 | 220,582 | 244,615 | 57,000 | 54,576 | 46,311 | 195,781 | 394,279 | 79,980 |  |  |
| **Ctrl-2** | 29,895 | 225,112 |  | 9,659 | 16,615 | 8,012 | 6,362 | 33,976 | 31,401 | 16,153 | 10,023 | 11,259 | 23,131 | 47,431 | 11,090 |  |  |
| **Ctrl-3** | 35,395 | 421,402 |  | 4,790 | 26,755 | 144,112 | 4,811 | 36,846 | 51,656 | 11,416 | 11,167 | 12,371 | 25,146 | 66,135 | 9,878 | 10,155 | 6,164 |
| **OE-PITPNA-1** | 24,064 | 1,529,392 |  | 21,265 | 109,161 | 25,895 | 20,344 | 153,847 | 223,080 | 54,644 | 57,849 | 50,001 | 231,713 | 516,372 | 65,222 |  |  |
| **OE-PITPNA-2** | 26,574 | 1,610,130 |  | 21,818 | 135,714 | 36,188 | 21,266 | 211,463 | 307,309 | 66,949 | 55,157 | 52,748 | 243,633 | 382,430 | 75,455 |  |  |
| **OE-PITPNA-3** | 36,628 | 911,196 |  | 1,048 | 15,337 | 97,397 | 13,094 | 93,187 | 122,465 | 33,599 | 23,468 | 21,003 | 89,036 | 292,763 | 52,211 | 23,016 | 33,573 |
| **Normalized to Internal Standard** | | | | | | | | | | | | | | | | | |
|  |  | **SUM** | **32 0** | **34 0** | **34 1** | **34 2** | **36 0** | **36 1** | **36 2** | **36 3** | **36 4** | **38 2** | **38 3** | **38 4** | **38 5** | **40 4** | **40 5** |
| **Ctrl-1** |  | 344.20 |  | 5.82 | 36.86 | 10.98 | 5.12 | 48.69 | 53.99 | 12.58 | 12.05 | 10.22 | 43.21 | 87.03 | 17.65 |  |  |
| **Ctrl-2** |  | 150.60 |  | 6.46 | 11.12 | 5.36 | 4.26 | 22.73 | 21.01 | 10.81 | 6.71 | 7.53 | 15.48 | 31.73 | 7.42 |  |  |
| **Ctrl-3** |  | 238.11 |  | 2.71 | 15.12 | 81.43 | 2.72 | 20.82 | 29.19 | 6.45 | 6.31 | 6.99 | 14.21 | 37.37 | 5.58 | 5.74 | 3.48 |
| **OE-PITPNA-1** |  | 1271.11 |  | 17.67 | 90.73 | 21.52 | 16.91 | 127.87 | 185.41 | 45.42 | 48.08 | 41.56 | 192.58 | 429.17 | 54.21 |  |  |
| **OE-PITPNA-2** |  | 1211.79 |  | 16.42 | 102.14 | 27.24 | 16.01 | 159.15 | 231.28 | 50.39 | 41.51 | 39.70 | 183.36 | 287.82 | 56.79 |  |  |
| **OE-PITPNA-3** |  | 497.55 |  | 0.57 | 8.37 | 53.18 | 7.15 | 50.88 | 66.87 | 18.35 | 12.81 | 11.47 | 48.62 | 159.86 | 28.51 | 12.57 | 18.33 |
